# Landscape-Based Mutational Sensitivity Cartography and Network Community Analysis of the SARS-CoV-2 Spike Protein Structures: Quantifying Functional Effects of the Circulating Variants

**DOI:** 10.1101/2021.05.18.444742

**Authors:** Gennady M. Verkhivker, Steve Agajanian, Deniz Yazar Oztas, Grace Gupta

## Abstract

Structural and biochemical studies SARS-CoV-2 spike mutants with the enhanced infectivity have attracted significant attention and offered several mechanisms to explain the experimental data. In this study, we used an integrative computational approach to examine molecular mechanisms underlying functional effects of the D614G mutation by exploring atomistic modeling of the SARS-CoV-2 spike proteins as allosteric regulatory machines. We combined atomistic simulations, deep mutational scanning and sensitivity mapping together with the network-based community analysis to examine structures of the native and mutant SARS-CoV-2 spike proteins in different functional states. Conformational dynamics and analysis of collective motions in the SARS-CoV-2 spike proteins demonstrated that the D614 position anchors a key regulatory cluster that dictates functional transitions between open and closed states. Using mutational scanning and sensitivity analysis of the spike residues, we identified the evolution of stability hotspots in the SARS-CoV-2 spike structures of the mutant trimers. The results offer support to the reduced shedding mechanism of as a driver of the increased infectivity triggered by the D614G mutation. By employing the landscape-based network community analysis of the SARS-CoV-2 spike proteins, our results revealed that the D614G mutation can promote the increased number of stable communities in the open form by enhancing the stability of the inter-domain interactions. This study provides atomistic view of the interactions and stability hotspots in the SARS-CoV-2 spike proteins, offering a useful insight into the molecular mechanisms of the D614G mutation that can exert its functional effects through allosterically induced changes on stability of the residue interaction networks.

## Introduction

Understanding of the molecular principles driving the coronavirus disease 2019 (COVID-19) associated with the severe acute respiratory syndrome (SARS)^1–5^ has been at the focal point of biomedical research since the start of the pandemic a year ago. SARS-CoV-2 infection is transmitted when the viral spike (S) glycoprotein binds to the host cell receptor, leading to the entry of S protein into host cells and membrane fusion.^6–8^ Structural and biochemical studies have shown that the mechanism of virus infection may involve spontaneous conformational transformations of the SARS-CoV-2 S protein between a spectrum of closed and receptor-accessible open forms, where RBD continuously switches between “down” and “up” positions where the latter can promote binding with the host receptor ACE2.^9–11^

The full-length SARS-CoV-2 S protein consists of two main domains, amino (N)-terminal S1 subunit and carboxyl (C)-terminal S2 subunit. The subunit S1 includes an N-terminal domain (NTD), the receptor-binding domain (RBD), and two structurally conserved subdomains (SD1 and SD2). The S2 subunit is an evolutionary conserved modlue that contains upstream helix (UH), an N-terminal hydrophobic fusion peptide (FP), fusion peptide proximal region (FPPR), heptad repeat 1 (HR1), central helix region (CH), connector domain (CD), heptad repeat 2 (HR2), transmembrane domain (TM) and cytoplasmic tail (CT).^12^ The S1 regions serve as dynamic protective shield of the fusion machine whereby upon proteolytic activation at the S1/S2 and dissociation of S1 from S2, structural rearrangements in S2 mediate the fusion of the viral and cellular membranes.^13–19^ The cryo-EM structures of the SARS-CoV-2 S proteins characterized distinct conformational arrangements of the S protein trimers in the prefusion form that are manifested by a dynamic equilibrium between the closed (“RBD-down”) and the receptor-accessible open (“RBD-up”) form required for the S protein fusion to the viral membrane.^20–29^ A population-shift between a spectrum of closed states that includes a structurally rigid closed form and more dynamic closed states can precede a transition to the fully open S conformation.^30–34^ The cryo-EM structures and biophysical tomography characterized the structures of the SARS-CoV-2 S trimers in situ on the virion surface and confirmed a population shift between different functional states, showing that conformational transitions can proceed through an obligatory intermediate in which all three RBD domains are in the closed conformations and are oriented towards the viral particle membrane.^35,36^ Conformational events associated with ACE2 binding may include progression from a compact closed form of the SARS-CoV-2 S protein that becomes weakened after furin cleavage between the S1 and S2 domains to the partially open states and fully open and ACE2-bound form priming the protein for fusion activation.^37^ Cryo-EM structures of SARS-CoV-2 S protein in the presence and absence of ACE2 receptor demonstrated that pH-dependent refolding region (residues 824-858) at the interdomain interface displayed dramatic structural rearrangements and mediated coordinated movements of the entire trimer, giving rise to a single 1 RBD-up conformation at pH 5.5 while all-down closed conformation was favorable at lower pH.^38^

SARS-CoV-2 S mutants with the enhanced infectivity profile including D614G mutational variant have attracted an enormous attention in the scientific community following the evidence of the mutation enrichment via epidemiological surveillance, resulting in proliferation of experimental data and a considerable variety of the proposed mechanisms explaining functional observations.^39–41^ The biochemical studies suggested a phenotypic advantage and the enhanced infectivity conferred by the D614G mutation.^42^ The D614G mutation can act by shifting the population of the SARS-CoV-2 S trimer from the closed form (53% of the equilibrium) in the native spike protein to a widely-open topology of the “up” protomers in the D614G mutant with 36% of the population adopting a single open protomer, 39% with two open protomers and 20% with all three protomers in the open conformation.^43^ The cryo-EM structures of the S-D614 and S-G614 ectodomains and the structure of the cleaved S-G614 ectodomain showed the increased population of the 1-RBD-up open form compared to the closed state in the S-GSAS/D614G structure.^44^ These experiments suggested that the D614G mutation in the SD2 domain can induce allosteric effect leading to structural shifts between the up and down RBD conformations.^44^ The electron microscopy analysis also revealed the higher 84% percentage of the 1-up RBD conformation in the S-G614 protein.^45^ The retroviruses pseudotyped with S-G614 showed a markedly greater infectivity than the S-D614 protein that was correlated with a reduced S1 shedding, greater stability of the S-G614 mutant and more significant incorporation of the S protein into the pseudovirion.^46^ It was also proposed that D614 S mutation may allosterically impact RBD-ACE2 binding and alter the transitions of “up” and “down” forms.^47^ A mechanistic explanation for the increased stability of the highly infective mutant was proposed based on the cryo-EM structures of a full-length S-G614 trimer with three distinct conformations representing a closed conformation, an intermediate closed form and a partially open 1 RBD-up conformation.^48^ According to this study, D614G eliminates a salt bridge between D614 of one subunit and K854 of the adjacent subunit but can promote ordering of the partly disordered loop (residues 620-640) which can strengthen the intra- and inter-domain interactions and enhance the stability of the mutated S protein. This study provided support to the reduced shedding mechanism induced by the D614G mutation that inhibits a premature dissociation of the S1 subunit which eventually leads to the increased number of functional spikes and stronger infectivity.^48^ Structure-based protein design and cryo-EM structure determination established that D614G and D614N mutations can result in the increased stability due to a decrease in the premature shedding of the S1 domain.^49^ Nonetheless, a consensus view on the exact mechanism underlying the functional effects and increased infectivity of S-D614G spike mutant is yet to be established. Several prevalent mechanisms are actively debated in the field offered to explain diverse experimental data, including D614G-induced modulation of cleavage efficiency of S protein; “openness” scenario advocating mutation-induced shift to the open states favorable for RBD-ACE2 interaction; “density” hypothesis suggesting a more efficient S incorporation into the virion; and “stability” mechanism that implicates mutation-induced enhancement in the association and stability of prefusion spike trimers as a driving force of greater infectivity.^42^

Molecular dynamics (MD) simulations recently probed the effects of the D614G mutation, suggesting that mutational variant favors an open conformation in which S-G614 protein maintains the symmetry in the number of persistent contacts between the three protomers.^50,51^ The normal mode analyses combined with Markov model and computation of transition probabilities characterized the dynamics of the S protein and mutational variants, predicting the increase in the open state occupancy for the more infectious D614G mutation due to the increased flexibility of the closed state and the enhanced stability of the open spike form.^52^ Computer-based mutagenesis and energy analysis of the thermodynamic stability of the S-D614 and S-G614 proteins in the closed and partially open conformations showed that local interactions near D614 position are energetically frustrated and may create an unfavorable environment that is stabilized in the S-G614 mutant through strengthening of the inter-protomer association between S1 and S2 regions.^53^ Using time-independent component analysis (tICA) and protein graph connectivity network, another computational study identified the hotspot residues that may exhibit long-distance coupling with the RBD opening, showing that the D614G could exert allosteric effect on the flexibility of the RBD regions.^54^ Structure-based physical model showed that the D614G mutation may induce a packing defect in S1 that promotes closer association and stronger interactions with S2 subunit, thereby supporting the reduced shedding hypothesis.^55^ Computational modeling and MD simulations have been instrumental in predicting dynamics and function of SARS-CoV-2 glycoproteins.^56–64^ The growing body of computational modeling studies investigating dynamics and molecular mechanisms of S mutational variants produced interesting but often inconsistent data that fit different mechanisms.

In this study, we used examine molecular mechanisms underlying the functional effects of the D614G mutation in different states of the SARS-CoV-2 S protein. We combined MD simulations with the ensemble-based mutational scanning and energetic sensitivity cartography analysis to simulate cryo-EM structures of the SARS-CoV-2 D614 and G614 proteins. By employing mutational sensitivity mapping and network-based community analysis for characterization of protein stability and allosteric interactions, we show that the D614G mutation can improve local stability of the spike protein and promote the larger number of stable communities in the open form by also enhancing the S1-S2 inter-domain interactions. This study offers support to the reduced shedding hypothesis of the D614G mutant function and provides a novel insight into the molecular mechanisms of the D614G mutation by examining SARS-CoV-2 S protein as an allosteric regulatory machine.

## Materials and Methods

### Structure Preparation and Analysis

Atomistic simulations were performed for the cryo-EM structure of SARS-CoV-2 S-GSAS/D614 in the closed-all down state (pdb id 7KDG), S-GSAS/D614 in the 1 RBD-up open state (pdb id 7KDH), SARS-CoV-2 S-GSAS/G614 mutant ectodomain in the closed form (pdb id 7KDK), SARS-CoV-2 S-GSAS/G614 mutant in the 1 RBD-up open state (pdb id 7KDL) as well as furin cleaved SARS-CoV-2 spike mutant S-RRAR/G614 in the closed form (pdb id 7KDI) and 1 RBD-up open state (pdb id 7KDJ) (Table 1, Figure 1).

**Figure 1.**
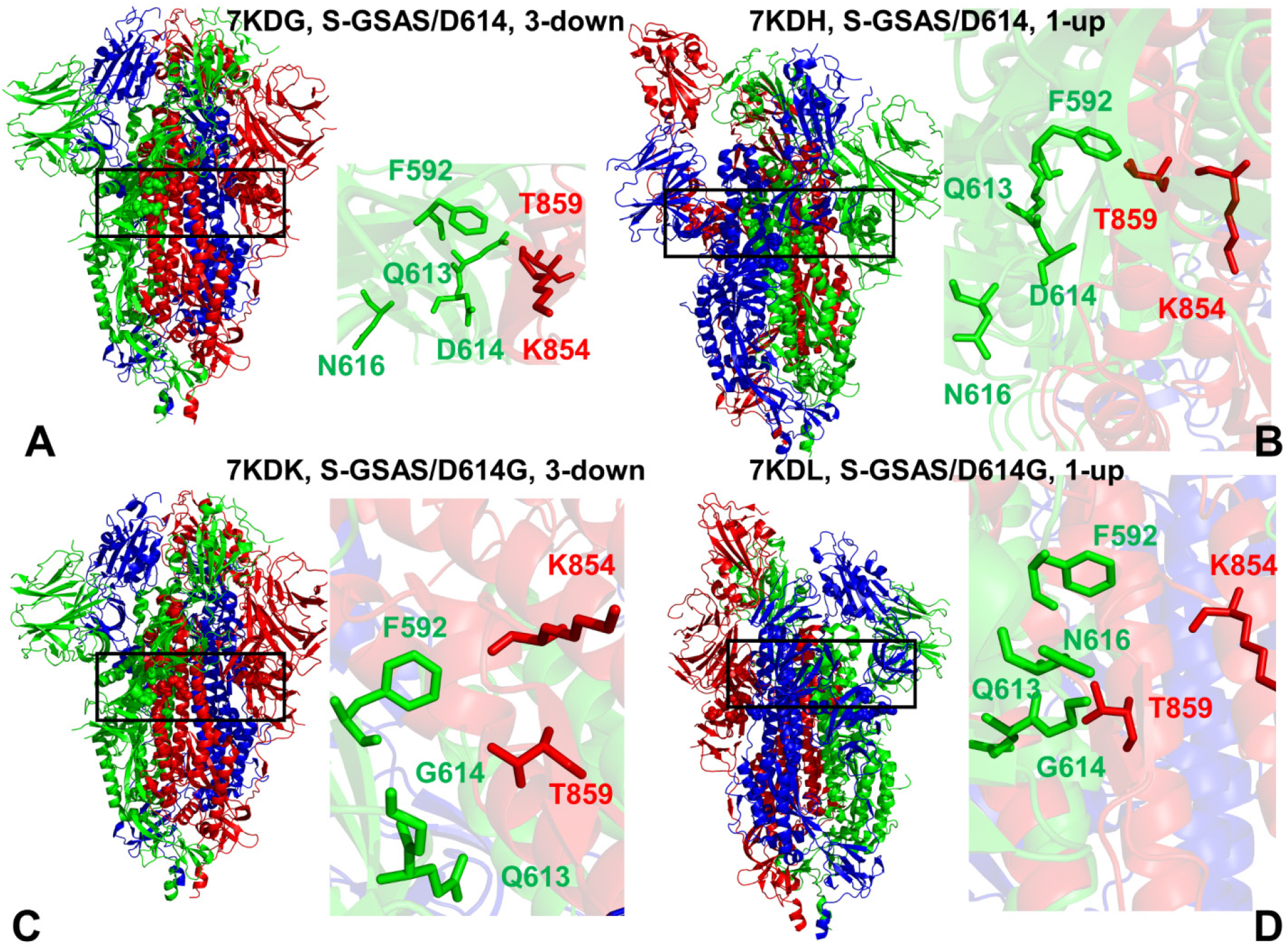
Cryo-EM structures of the SARS-CoV-2 S trimer structures used in this study. (A) The cryo-EM structure of SARS-CoV-2 S-GSAS/D614 in the closed state (pdb id 7KDG). The structure is in ribbons with protomers A,B,C are colored in green, red and blue. A close-up of the key residues and interactions near D614 position. D614, Q613, N616 of protomer A (green sticks) and K854, T859 of the adjacent protomer B are in red sticks. (B) S-GSAS/D614 in the 1 RBD-up open states (pdb id 7KDH). A close-up of the key residues and interactions near D614 position is shown. (C) The cryo-EM structure of SARS-CoV-2 S-GSAS/G614 mutant ectodomain in the closed form (pdb id 7KDK). A close-up of the key residues and interactions is shown. (D) The cryo-EM structure the SARS-CoV-2 S-GSAS/G614 mutant in the 1 RBD-up open states (pdb id 7KDL). A close-up of the key residues and interactions near D614 position is shown and residues are annotated as in panel (A).

**Table 1.** Structures of SARS-CoV2 spike protein structures examined in this study.

| PDB code | Description | RBDs position |
| --- | --- | --- |
| 6XR8 | SARS-CoV-2 Spike Protein Trimer D614 locked closed | 3 down |
| 7KDG | SARS-CoV-2 Spike Protein Trimer (S-GSAS) D614 | 3 down |
| 7KDH | SARS-CoV-2 Spike Protein Trimer (S-GSAS) D614 | 1 up |
| 7KDK | SARS-CoV-2 Spike Protein Trimer (S-GSAS) G614 | 3 down |
| 7KDL | SARS-CoV-2 Spike Protein Trimer (S-GSAS) G614 | 1 up |
| 7KRQ | SARS-CoV-2 Spike Protein Trimer G614 locked closed | 3 down |
| 7KRS | SARS-CoV-2 Spike Protein Trimer G614 intermediate closed | 1-half up |
| 7KRR | SARS-CoV-2 Spike Protein Trimer G614 open | 1 up |

All structures were obtained from the Protein Data Bank.^65,66^ Hydrogen atoms and missing residues were initially added and assigned according to the WHATIF program web interface.^67,68^ The structures were further pre-processed through the Protein Preparation Wizard (Schrödinger, LLC, New York, NY) and included the check of bond order, assignment and adjustment of ionization states, formation of disulphide bonds, removal of crystallographic water molecules and co-factors, capping of the termini, assignment of partial charges, and addition of possible missing atoms and side chains that were not assigned in the initial processing with the WHATIF program. The cryo-EM structures of the SARS-CoV-2 S proteins contained a number of missing loops of various lengths that were dispersed throughout the structure and required a rigorous template-based loop modeling and reconstruction. The modeled missing loops were located in the NTD regions (residues 1-26, 70-79,144-164,173-185,246-262, 294-304), RBD regions (residues 469-488) CTD2 regions (residues 621-640), the cleavage site at the S1/S2 boundary (residues 677-689) and residues in the S2 subunit (residues 828-853). The missing loops in the studied cryo-EM structures of the SARS-CoV-2 S protein were reconstructed and optimized using template-based loop prediction approaches ModLoop,^69^ ArchPRED server^70^ and further confirmed by FALC (Fragment Assembly and Loop Closure) program.^71^ The side chain rotamers were refined and optimized by SCWRL4 tool.^72^ The protein structures were then optimized using atomic-level energy minimization with a composite physics and knowledge-based force fields using 3Drefine method.^73^

In addition to the experimentally resolved glycan residues present in the cryo-EM structures of studied SARS-CoV-2 S proteins, the glycosylated microenvironment for simulation trajectories was mimicked by using the structurally resolved glycan conformations for 16 out of 22 most occupied N-glycans in each protomer as determined in the cryo-EM structures of the SARS-CoV-2 spike S trimer in the closed state (K986P/V987P,) (pdb id 6VXX) and open state (pdb id 6VYB) and the cryo-EM structure SARS-CoV-2 spike trimer (K986P/V987P) in the open state (pdb id 6VSB).

### MD Simulations

The structures of the SARS-CoV-2 S trimers were simulated in a box size of 85 Å × 85 Å × 85 Å with buffering distance of 12 Å. Assuming normal charge states of ionizable groups corresponding to pH = 7, sodium (Na+) and chloride (Cl-) counter-ions were added to achieve charge neutrality and a salt concentration of 0.15 M NaCl was maintained. All Na^+^ and Cl^−^ ions were placed at least 8 Å away from any protein atoms and from each other. All-atom MD simulations were performed for an N, P, T ensemble in explicit solvent using NAMD 2.13 package^74^ with CHARMM36 force field.^75^ Long-range non-bonded van der Waals interactions were computed using an atom-based cutoff of 12 Å with switching van der Waals potential beginning at 10 Å. Long-range electrostatic interactions were calculated using the particle mesh Ewald method^76^ with a real space cut-off of 1.0 nm and a fourth order (cubic) interpolation. SHAKE method was used to constrain all bonds associated with hydrogen atoms. Simulations were run using a leap-frog integrator with a 2 fs integration time step. Energy minimization after addition of solvent and ions was carried out using the steepest descent method for 100,000 steps. All atoms of the complex were first restrained at their crystal structure positions with a force constant of 10 Kcal mol^−1^ Å^−2^. Equilibration was done in steps by gradually increasing the system temperature in steps of 20K starting from 10K until 310 K and at each step 1ns equilibration was done keeping a restraint of 10 Kcal mol-1 Å-2 on the protein C_α_ atoms. After the restrains on the protein atoms were removed, the system was equilibrated for additional 10 ns. An NPT production simulation was run on the equilibrated structures for 500 ns keeping the temperature at 310 K and constant pressure (1 atm). In simulations, the Nose–Hoover thermostat^77^ and isotropic Martyna–Tobias–Klein barostat^78^ were used to maintain the temperature at 310 K and pressure at 1 atm respectively.

### Mutational Scanning and Sensitivity Mapping

To compute protein stability changes in the SARS-CoV-2 S structures, we conducted a systematic alanine scanning of protein residues in the SARS-CoV-2 trimer mutants as well as mutational sensitivity analysis at the mutational site for both SARS-CoV-2 S-D614 and SARS-CoV-2 S-G614 structures. Mutational scanning of protein residues was performed using BeAtMuSiC approach.^79–81^ If a free energy change between a mutant and the wild type (WT) proteins ΔΔG= ΔG (MT)-ΔG (WT) > 0, the mutation is destabilizing, while when ΔΔG <0 the respective mutation is stabilizing. BeAtMuSiC approach is based on statistical potentials describing the pairwise inter-residue distances, backbone torsion angles and solvent accessibilities, and considers the effect of the mutation on the strength of the interactions at the interface and on the overall stability of the complex. We leveraged rapid calculations based on statistical potentials to compute the ensemble-averaged alanine scanning computations and mutational sensitivity analysis at D614 and G614 positions using equilibrium samples from reconstructed simulation trajectories.

### Dynamic-Based Modeling of Residue Interaction Network and Community Analysis

A graph-based representation of protein structures^82^ is used to represent residues as network nodes and the inter-residue edges to describe residue interactions. We constructed the residue interaction networks using both dynamic correlations^83^ and coevolutionary residue couplings^84^ that provide complementary descriptions of the allosteric interactions and yield robust network signatures of long-range couplings and communications. The details of network construction were described in our previous studies.^85^ The ensemble of shortest paths is determined from matrix of communication distances by the Floyd-Warshall algorithm.^86^ Network graph calculations were performed using the python package NetworkX.^87^

The Girvan-Newman algorithm^88–90^ is used to identify local communities. An improvement of Girvan-Newman method was implemented where all highest betweenness edges are removed at each step of the protocol. The algorithmic details of this modified scheme were presented in our recent study.^91,92^ The network parameters were computed using the python package NetworkX^87^ and Cytoscape package for network analysis.^93,94^ A community-based analysis and modularity assessment of allosteric interaction networks is based on the notion that groups of residues that form local interacting communities are expected to be highly correlated and can switch their conformational states cooperatively. As a result, long-range allosteric interaction signals can be transmitted through a hierarchical chain of local communities on the dynamic interaction networks. The community analysis and a comparative evaluation of the communities in the SARS-CoV-2 S-D614 and S-G614 structures are used as proxy for measuring global protein stability and communication efficiency in performing allosteric functions. This analysis provides also a global metric to measure changes in connectivity and interaction between subdomains and inter-protomer association in the SARS-CoV-2 S structures.^95^

## Results and Discussion

### MD Simulations of the SARS-CoV-2 Spike Proteins in the Closed and Open Forms of the SARS-CoV-2 S Trimer

The conformational dynamics profiles of the SARS-CoV-2 S-GSAS/D614 and SARS-CoV-2 S-GSAS/G614 showed a more flexible S1 subunit and very stable S2 subunit in both closed form (Figure 2A) and 1-up open state (Figure 2B). The thermal fluctuations of both S1 and S2 regions were smaller in the closed state with RMSF < 1.0 Å. In the closed state, the RBD (residues 331-528) and CTD1 (residues 528-591) corresponded to the most stable regions in the S1 subunit, while UH (residues 736-781) and CH (residues 986-1035) were the most stable regions in the S2 subunit (Figure 2A). Only marginally larger fluctuations were seen in the CTD2 region (residues 592-686) that connects S1 and S2 subunits. Our analysis showed that the conformational dynamics profiles were generally similar for the closed forms of both the SARS-CoV-2 S-GSAS/D614 and S-GSAS/G614 mutant (Figure 2A). Molecular simulations did not detect any radical changes in the dynamics profile of the D614G mutant, indicating that the effect of mutation on the conformational dynamics could be subtle and amount to a number of small local changes dispersed throughout the protein structure. A significant increase in the conformational mobility of the S-D614 and S-D614G open states was seen for the RBD-up protomer (Figure 2B). The “up” protomer can undergo fluctuations up to RMSF = 2.5-3.5 Å also revealing signs of moderately smaller fluctuations and greater stability for the open form of the S-G614 mutant (Figure 2B). Of some notice were moderately greater residue displacements in the NTD regions of the S-D614 and S-G614 open structures, including the NTDs of the two closed-down protomers (Figure 2B, D). In general, the dynamic profiles of the S-D614 and S-G614 trimers remained similar in the native and mutant open forms.

**Figure 2.**
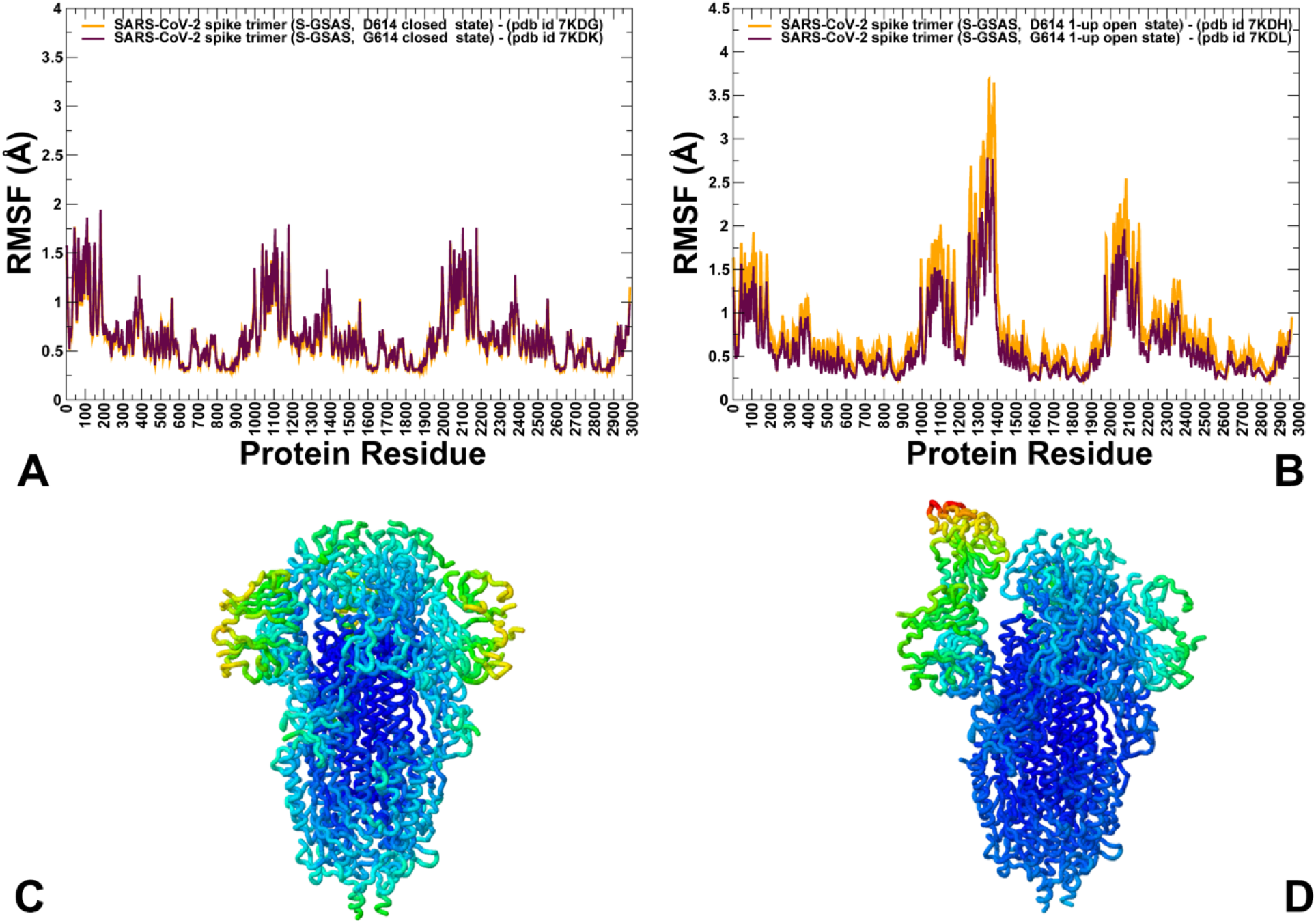
Conformational dynamics of the SARS-CoV-2 spike (S) protein trimer mutants. (A) The root mean square fluctuations (RMSF) profiles obtained from simulations of the cryo-EM structure of SARS-CoV-2 S-GSAS/D614 in the closed-all down state (pdb id 7KDG) and SARS-CoV-2 S-GSAS/G614 mutant ectodomain in the closed form (pdb id 7KDK). (B) The RMSF profiles for S-GSAS/D614 in the 1 RBD-up open state (pdb id 7KDH) and SARS-CoV-2 S-GSAS/G614 mutant in the 1 RBD-up open states (pdb id 7KDL). The profiles are shown for protomer chains A,B and C in sequential order. (C) Structural map of the conformational mobility profiles for the SARS-CoV-2 S-GSAS/D614 trimer in the closed form. (D) Structural map of the conformational mobility profiles for the SARS-CoV-2 S-GSAS/D614 trimer in the partially open 1 RBD-up state. The colored-coded maps show the rigidity-flexibility changes in blue-to-red spectrum with the most stable regions in blue and most flexible residues in red.

Several previous studies suggested that the molecular basis of the increased infectivity for the D614G mutant may be explained via “openness” hypothesis^50–52^ according to which D614G modification can alter conformational dynamics signatures of the S protein and promote switching to the open conformation, positioning a single or multiple RBDs to make contacts with the ACE2 receptor. Although our results showed no substantial alteration of the dynamic signatures caused by the D614G mutation, simulations pointed to a marginally greater stabilization of the open form for the S-G614 mutant which is consistent with previous assertions.^50–52^ To provide a more detailed analysis of the intra- and inter-protomer contacts in the closed and open states, we computed the ensemble-averaged numbers of these contacts for the SARS-CoV-2 S-GSAS/D614 and SARS-CoV-2 S-GSAS/G614 structures (Tables S1-S4) using a contacts-based Prodigy approach.^96,97^ This analysis showed that the number of inter-protomer contacts is consistently greater in the closed state of the S-D614 and S-G614 proteins as compared to the partially open state. According to this assessment, D614 and Q613 residues can anchor the intra-protomer clusters with V597, S596 and T315 of the same protomer, while establishing contacts with T859, L861 and S735 residues of the adjacent protomers (Tables S1,S2). It is worth noting that in agreement with several computational studies advocating for the “openness” mechanism^50^ we also found a preferentially symmetric nature of the inter-protomer contacts in the closed state, while the number of inter-protomer contacts can be considerably reduced in the partially open form of S-D614 conformation (Tables S1,S2). These changes could become less pronounced in the S-G614 form, but the total number of the inter-protomer contacts is consistently reduced in the open state regardless of the mutational status of D614 position (Tables S3,S4). A close inspection of the local dynamics profiles revealed only minor flexibility changes near the mutational site, where the loss of favorable interactions with T859 of the adjacent protomer may be compensated through the intra-protomer contacts especially for the up protomer in the open state. It was proposed that a more flexible closed state would favor the opening of the S-D614 spike protein, while a more rigid open state of D614G would shift the conformational equilibrium towards the open state.^52^ Our results revealed small and largely synchronous dynamic changes in the closed and open forms of the D614G mutant, showing no indication of a dramatic alteration of the dynamic signatures to suggest an obvious trigger for the dynamic preferences of the D614G mutant towards partially open form as was proposed in a computational study.^52^

We also performed MD simulations of the SARS-CoV-2 S-D614 timer in the locked closed form and structures of the S-G614 mutant in the closed, intermediate and open forms (Figure S1) to examine mutation-induced control of conformational dynamics, functional motions and protein stability in the major functional states. The thermal fluctuations of both S1 and S2 subunits are very small in the locked state of the S-D614 trimer (Figure S2A). The mobility of the RBD residues was largely suppressed and only peripheral regions of the NTD showed some level of residual mobility while thermal stabilization of the structurally ordered FPPR motif (residues 828-853) was evident. Despite similarities in the conformational mobility profiles of the S-D614 and S-G614 mutant forms, there are some important changes. In the closed form of the S-G614 mutant, the FPPR segments (residues 823-862) and loops (residues 620-640) are fully ordered, while in the S-D614 locked closed state only FPPR region is structured (Figure S1A). Nonetheless, a more significant mobility could be observed for the S-G614 mutant, where both NTD and RBD regions experienced larger fluctuations (Figure S2B). In the intermediate conformation with a partly open single RBD, the loops are disordered in the RBD-shifted module and FPPR remain fully ordered (Figure S1B). For this S-G614 mutant form, there is further expansion of higher mobility regions in the NTD and RBD subunits (Figure S2C). Upon attaining the 1-up RBD open conformation (Figure S1C), S-G614 mutant featured only moderately increased mobility in the RBD of the up protomer, while the two closed protomers exhibited the enhanced rigidity with the essentially immobilized NTD and RBD regions (Figure S2D). While the covariance matrixes of residue fluctuations highlighted a general similarity of correlated motions, a more detailed comparison could point to stronger couplings between S1 and S2 domains in the 1-up open state of the S-G614 trimer (Figure S2E-G). Moreover, we also noticed the increased density of both the intra- and inter-protomer correlated motions in the open state (Figure S2G), suggesting that long-range couplings between functional regions could become stronger in the open mutant form. We argue that by enhancing the inter-domain communications in the open state, the D614G mutation can improve stability of the S-G614 trimer and limit shedding of the S1 subunit.

We characterized collective motions and determined the hinge regions in the SARS-CoV-2 S-D614 and S-G614 structures (Figure 3) using principal component analysis (PCA) of MD trajectories with the CARMA package.^98^ The regulatory regions that control collective motions are conserved as the key hinge positions are shared between different functional states and remained intact in the S-D614 trimer and G614 mutant trimer (Figure 3). The local minima regions in each protomer are localized around K310-F318 and S591-V618 regions in the S1 domain where the first cluster can modulate NTD-RBD motions while the second hinge center is responsible for collective dynamics between RBD and S2 regions.

**Figure 3.**
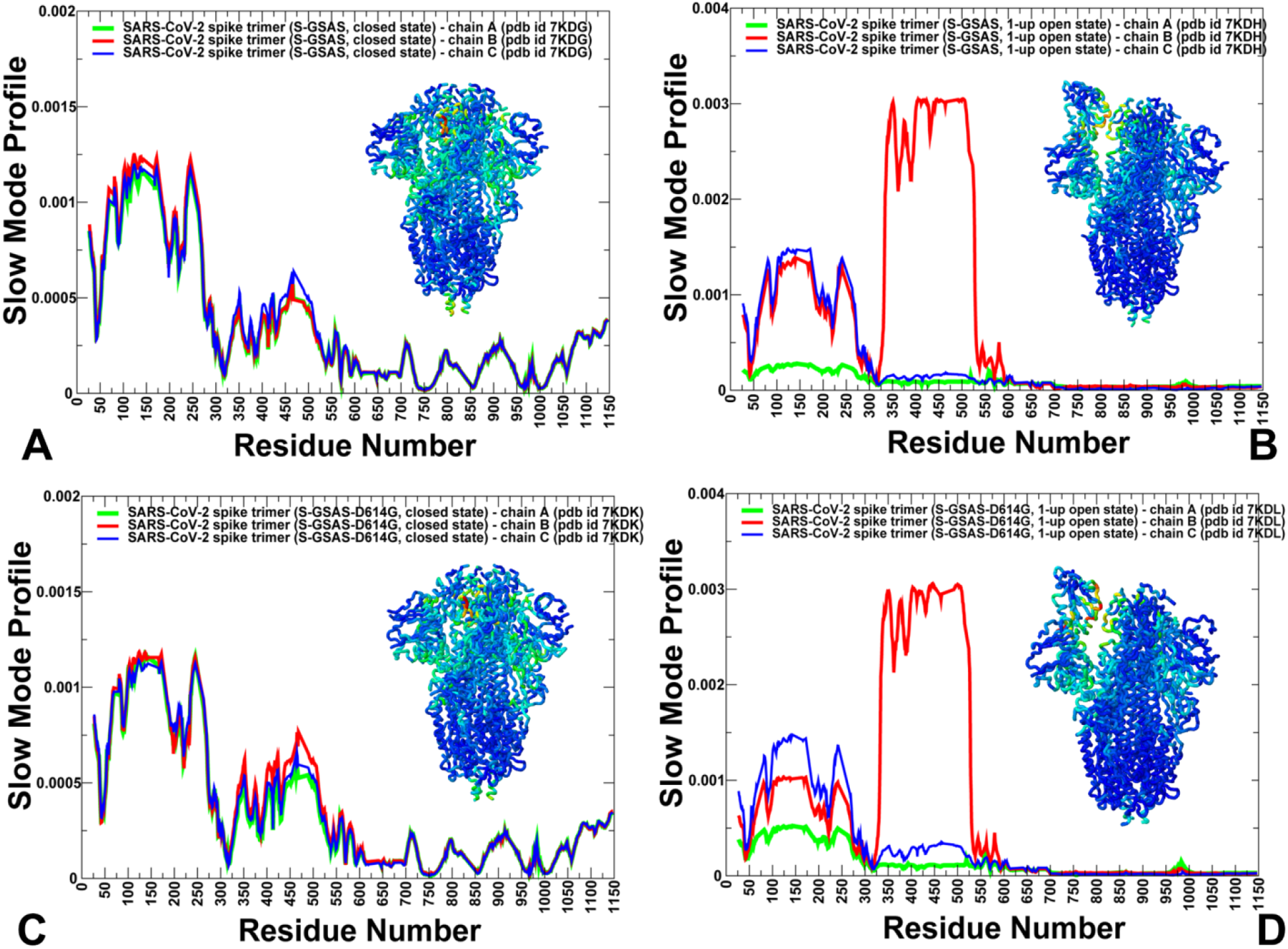
Functional dynamics of the SARS-Cov-2 S-GSAS/D614 structures in the closed and open forms. The essential mobility profiles are averaged over the three lowest frequency modes. (A) The essential mobility profiles for the SARS-Cov-2 S-GSAS/D614 in the closed form (pdb id 7KDG). (B) The slow mode profile for the SARS-Cov-2 S-GSAS/D614 structure in the 1-up open state (pdb id 7KDH). (C) The essential mobility profiles for the SARS-CoV-2 S-GSAS/G614 structure in the closed form (pdb id 7KDK). (D) The slow mode profile for the SARS-Cov-2 S-GSAS/G614 structure in the 1-up open state (pdb id 7KDL). The profiles for protomer chains A,B and C are shown in green, red and blue lines respectively. The inset on each panel shows structural maps of the SARS-CoV-2 S trimers colored by the main-chain deformability where high deformability regions are colored in red.

These observations are in line with experimental studies indicating that the D614G mutational site may be located in the immobilized structural region of the SD2 domain where local environment of D614 combined with β strand formed by residues 311–319 may correspond to a hinge center governing motions of NTD and RBD, as well as isolating the motions in S1 from the S2 subunit.^44^ Importantly, the key hinge residues F318, S591, F592, Y855, I856 residues correspond to the local minima of the essential mobility profile for each of the protomer and are involved in the inter-protomer interactions (Figure 3). These sites are located in the close proximity of D614 and could form several interaction clusters in both the closed and open states of the SARS-CoV-2 S-D614 and SARS-CoV-2 S-G614 structures. Structural mapping of high deformability regions associated with collective motions showed the broader distribution of these sites in the closed form and a strong but localized density for the partially open state. The structural conservation of the inter-protomer hinge centers is seen in the slow mode profiles pointing to the immobilization of the same regulatory positions that control collective dynamics across distinct states of the D614 and G614 S trimers (Figure 3).

A comparative analysis of the functional dynamics in the locked/closed, intermediate and open forms of the S-G614 mutant trimer provided an additional insight into spectrum of collective motions that can be modulated by mutations (Figure S3). In the locked closed form of the S-D614 trimer, the movements of the RBD and NTD regions were mostly suppressed (Figure S3A). Despite conservation of hinge sites, the collective dynamics in the S-G614 mutant trimer revealed its unique important signatures. In the closed and intermediate states, the functional dynamics profile of the S-G614 mutant trimer displayed moderate but appreciable displacements of the RBD regions in all protomers (Figure S3B,C). These dynamic preferences of the RBDs may lead to more frequent shifts between the closed and open forms and lower the kinetic barriers for the transitions between functional forms in the S-G614 mutant. Interestingly, in the open state of the S-G614 mutant trimer only the “up” protomer featured large displacements of the NTD and RBD region, while these regions in the closed-down protomers become mostly immobilized (Figure S3D). The observed pattern of collective motions in the S-G614 mutant would imply that the RBD regions may experience functional displacements in the closed and intermediate forms that promote transitions to the open form. However, once the S-G614 trimer attains the 1-up open form, the closed protomers could become rigidified with their RBD movements being largely suppressed. This may force the open RBD to return to the down conformation unless it becomes fully engaged by the ACE2 receptor as was proposed.^48^ Our findings are also consistent with experimental studies showing that the D614G mutation confers the increased structural flexibility in the closed states, which may trigger the enhanced exchange between the open and closed forms by increasing the dissociation rate and moderating binding affinity of the S-G614 trimer for ACE2 host receptor.

### Mutational Sensitivity Cartography in the SARS-CoV-2 Spike Trimers Reveal D614G- Induced Stabilization of the Closed and Open States

Previous mutagenesis analysis suggested that the D614G mutant may improve the stability of the S protein by strengthening the inter-protomer association between S1 and S2 regions.^53^ We employed the equilibrium ensembles generated by MD simulations of the SARS-CoV-2 protein structures to perform deep mutational scanning of the spike protein residues and mutational sensitivity analysis of the D614 S trimer (Figure 4) and G614 S mutant proteins (Figure 5). Using energetic perturbation analysis, we test the shedding hypothesis suggesting that stabilization of the S-G614 protein may preclude a premature dissociation of the S1 subunit, leading to the increased number of functional spikes and stronger infectivity.^48^

**Figure 4.**
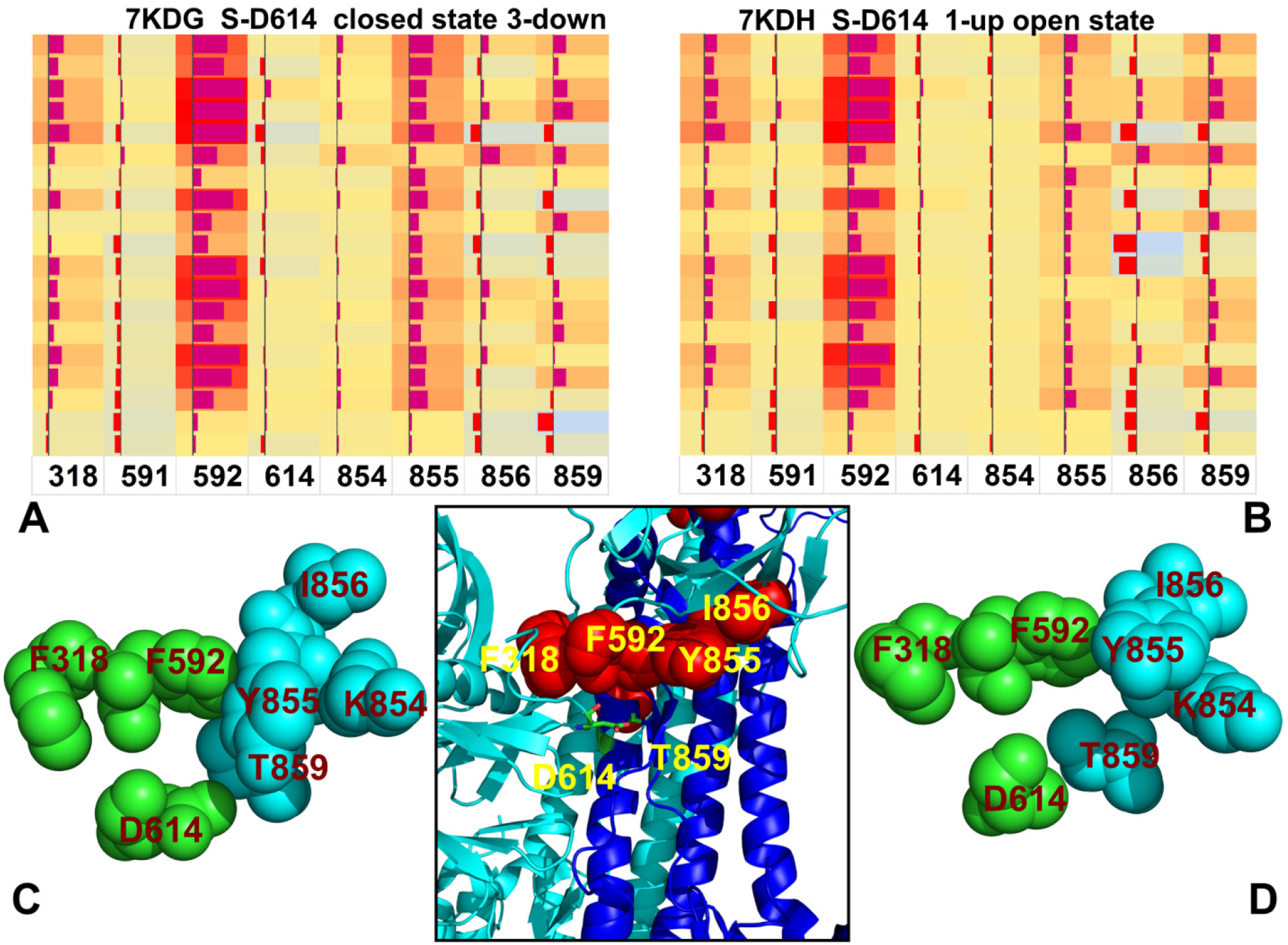
The mutational sensitivity analysis for the SARS-CoV-2 S-D614 trimer in the closed and open states. (A) The mutational scanning map of the S-D614 trimer in the closed form. (A) The mutational scanning map of the S-D614 trimer in the open form. The heatmaps show the effect of all single mutations at the hinge residue positions on the folding free energy changes. The squares on the heatmap are colored by mutational effect in a 3-colored scale from red to light blue with red indicating the largest destabilization effect. The data bars correspond to the computed binding free energy changes. (C) A close-up of the inter-protomer hinge cluster in the closed state. (D) A close-up of the inter-protomer hinge cluster in the open state. D614, F318 and F592 belong to one of the protomer and are shown in green spheres. K854, Y855, I856, and T859 belong to the other protomer and shown in cyan spheres. The central lower panel presents an overview of the inter-protomer cluster with residues in red spheres and annotated.

**Figure 5.**
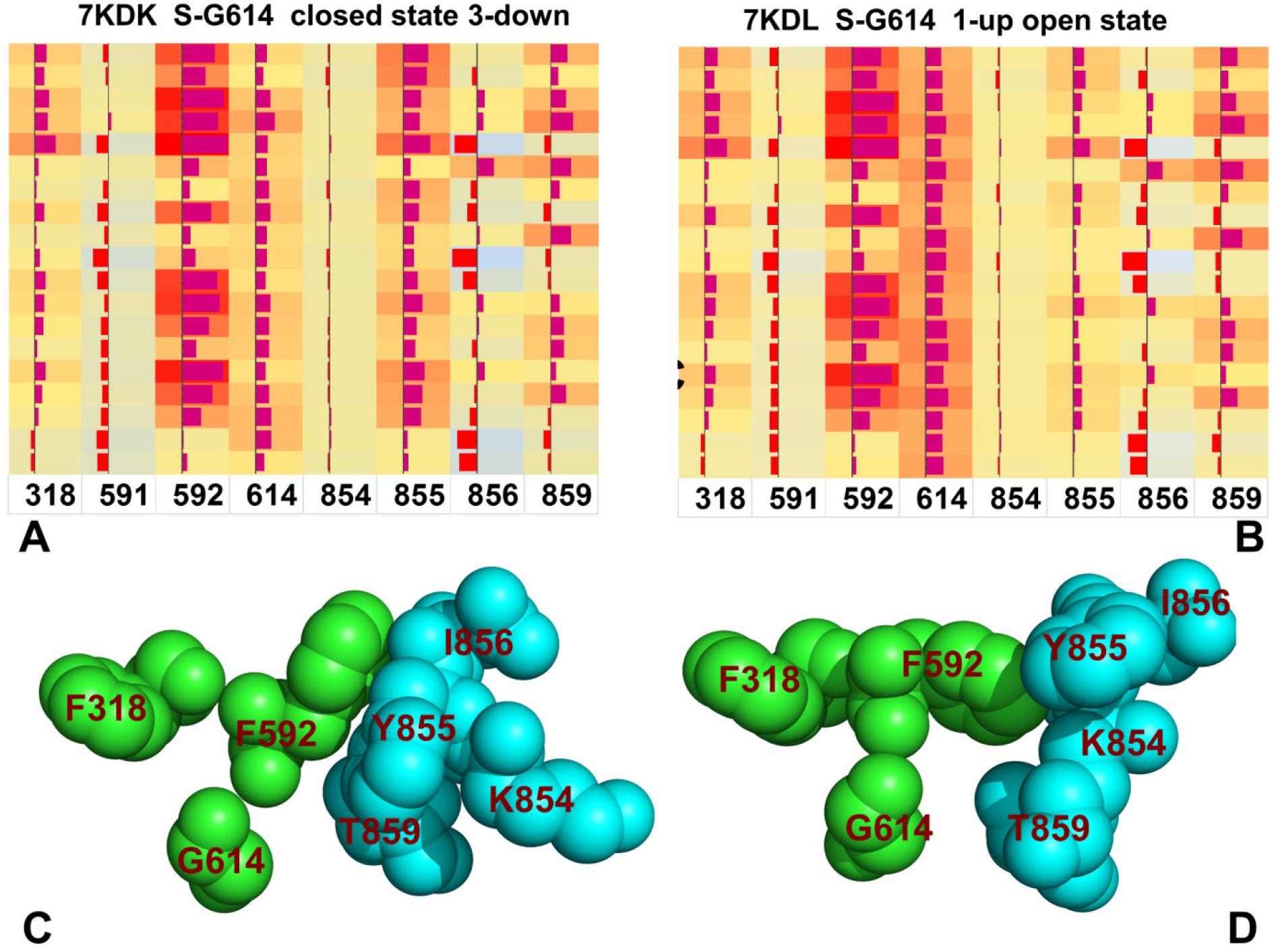
The mutational sensitivity analysis for the SARS-CoV-2 S-G614 trimer in the closed and open states. (A) The mutational scanning map of the S-G614 trimer in the closed form. (B) The mutational scanning map of the S-G614 trimer in the open form. The heatmaps show the effect of all single mutations at the hinge residue positions on the folding free energy changes. The squares on the heatmap are colored by mutational effect in a 3-colored scale from red to light blue with red indicating the largest destabilization effect. The data bars correspond to the computed binding free energy changes. (C) A close-up of the inter-protomer hinge cluster in the closed state. (D) A close-up of the inter-protomer hinge cluster in the open state. G614, F318 and F592 belong to one of the protomer and are shown in green spheres. K854, Y855, I856, and T859 belong to the other protomer and shown in cyan spheres.

We constructed mutational sensitivity heatmaps for the major hinge centers in the S-D614 closed trimer (Figure 4A) and partially open form (Figure 4B). Mutational cartography revealed several interesting trends and highlighted the energetic effect of D614 site. First, we noticed that the energetic hotspots corresponded to hinge centers F592 and Y855 that showed larger contributions in the closed state (Figure 4A). A similar heatmap was obtained for the S-D614 trimer in the open form where mutations in the Y855 hinge position caused large destabilization changes (Figure 4B). Importantly, D614 site appeared to be highly tolerable to modifications in both closed and open forms, yielding very minor and often stabilizing changes upon mutations. Structural mapping showed that hinge residues F518 and F592 in one protomer can form the inter-protomer cluster with K854, Y855, I856, and T859 hinge sites of the adjacent protomer (Figure 4C,D). D614 packs tightly with T859, K854, Y855, I856 of the other protomer, while also maintaining favorable contacts with F318 and F592 sites of the same protomer. Interestingly, D614 can effectively anchor these clusters by connecting major hinge residues (Figure 4C,D). Notably, tight packing in this cluster is preserved in both closed and open forms, which is mainly determined by the interactions between two conserved stability hotspots F592 and Y855 (Figure 4C,D). These hotspots protect structural stability of the inter-protomer hinge center and allow for mutational and energetic plasticity of other residues in the cluster. Hence, structural and mutational changes in the D614 position may be well-tolerated and accompanied by repacking of the inter-protomer hinge cluster thereby preserving functional role of this region in regulation of collective motions.

Mutational sensitivity mapping of the hinge residues in the S-G614 mutant trimer revealed a redistribution of the energetic hotspots (Figure 5A,B). In the closed form of the G614 trimer the stability hotspots corresponded to F592, Y855 and G614 sites as mutations in these positions consistently resulted in large destabilization changes (Figure 5A). Of particular interest is emergence of G614 as one of the stability hotspots that together with F592 and Y855 strengthen the inter-protomer hinge cluster (Figure 5C). Although D614G eliminates favorable interactions with T859 and K854, the tight packing is maintained between residues of the inter-protomer hinge cluster (Figure 5C). In the open form of the S-G614 trimer the stability hotspots F592 and G614 become dominant and anchor the inter-protomer hinge cluster (Figure 5B). Importantly, the heatmap showed that G614 is more stable in the open state than in the closed state which may drive conformational preference of the mutant trimer towards the 1-up form. Structural analysis of the inter-protomer cluster reflected a partial rearrangement of packing interactions where contacts between F592, G614 and Y855 residues determine stability of the hinge center (Figure 5D). The central finding of the mutational sensitivity mapping is a significant energetic plasticity of D614 in the native trimer, while in the S-G614 mutant trimer this position becomes the dominant energy hotspot of the inter-protomer hinge center.

In the closed form of the S-D614 trimer, most of D614 mutations can result in the improved stability, highlighting in particular an appreciable stabilization of the D614C, D614G, and D614N mutants (Figure 6A). Notably, the D614G mutation caused the largest energetic change as compared to other substitutions. A similar pattern was seen in the mutational profile of the open state where D614G and D614N mutations displayed favorable and comparable stabilization changes (Figure 6B).

**Figure 6.**
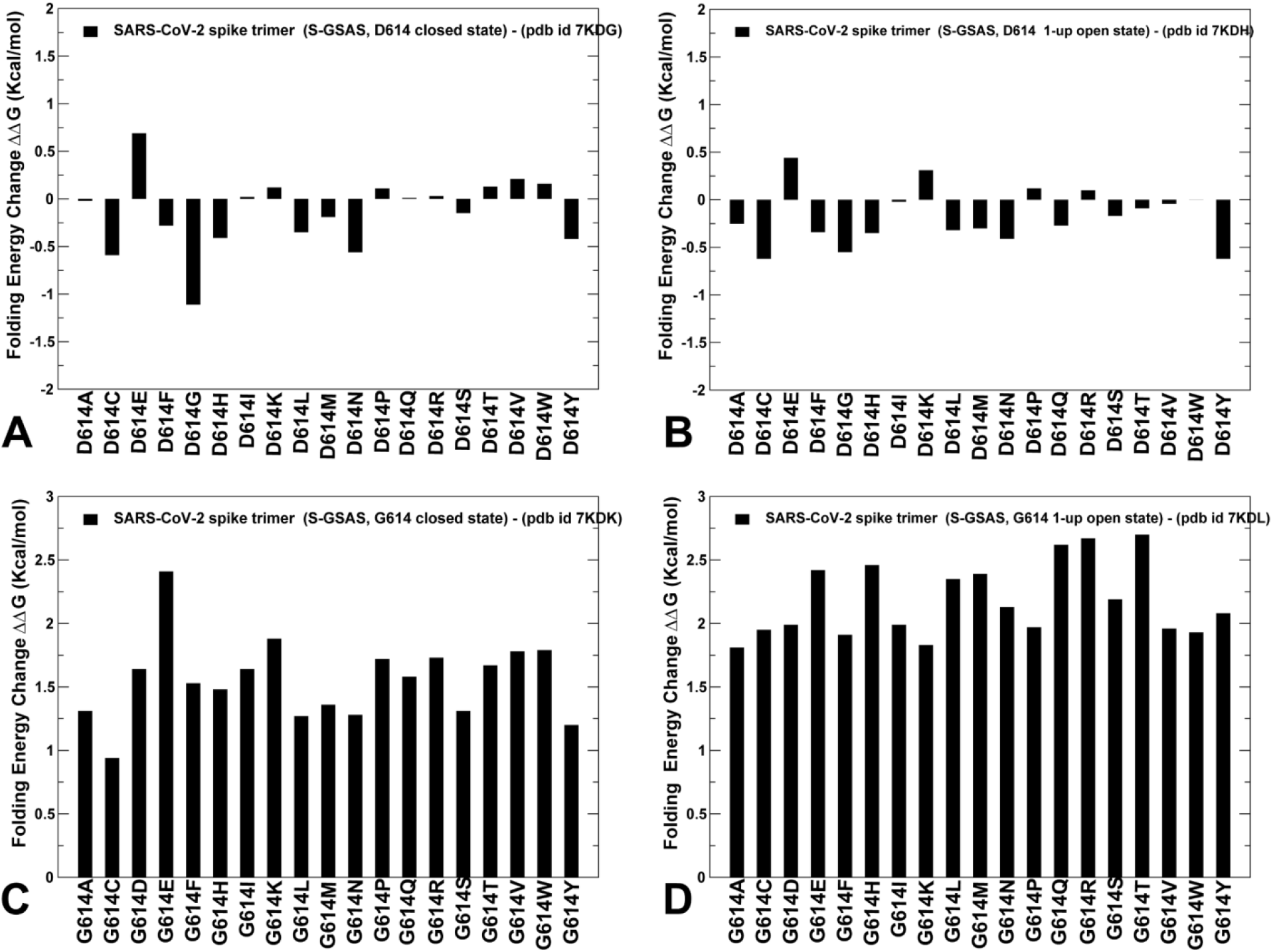
Mutational sensitivity analysis of the SARS-CoV-2 S-D614 and S-G614 trimers. Mutational sensitivity scanning of D614 position in the closed form of the S-D614 protein (A) and open form of the S-D614 protein (B). Mutational sensitivity scanning of G614 position in the closed form of the S-G614 protein (C) and S-G614 protein (D). The protein stability changes are shown in black filled bars.

These findings are consistent with the latest differential scanning fluorimetry studies showing that the D614G and D614N induced changes have a very similar effect, leading to a considerable improvement of thermal stability which may be explained by a decrease in premature shedding of S1 domain.^49^ Consistent with this experimental study, we found that D614G/N mutations can moderate the repulsive charge interactions at the interface between S1 and S2 domains and allow for the tighter inter-domain packing.^49^ Strikingly, mutational sensitivity analysis of the SARS-CoV-2 mutant structures revealed a far more significant destabilization effect of G614 substitutions in the closed and open forms (Figure 6C,D). The destabilization changes caused by mutations of G614 were greater in the open form, thereby indicating that D614G mutation may have a stronger stablization effect on the open state.

The mutational sensitivity profile of the SARS-CoV-2 S-D614 in the locked closed form showed moderate destabilization changes whereas D614G, D614C, D614H and D614N modifications resulted in the improved stability of the closed form (Figure S4A). This was particularly interesting given the overall rigidity of the locked closed S-D614 trimer, indicating that mutations in the D614 position can be accommodated even in this constrained state. In the locked closed form of S-G614 the largest destabilization changes were induced by the G614K, G614D and G614E mutations (Figure S4B), highligting the fact that the reverse G614D mutation could significantly destablize the S-G614 trimer. In the intermediate conformation, we observed larger destabilization changes for the moving protomer but also an appreciable loss of stability for the closed-down two protomers (Figure S4C). An interesting pattern of protein stability changes was seen in the 1 RBD-up open state of the S-G614 trimer (Figure S4D). The free energy changes induced by mutations of G614 in the open protomer (protomer A) were somewhat smaller, reflecting a larger mobility and openness of this protomer near the site of mutation. At the same time, the loss of protein stability upon modifications of G614 in the closed protomers were more significant, indicating the increased stabilization of the S-G614 trimer (Figure S4D). The results of the mutational scanning are consistent with the functional dynamics analysis, showing structural and energetic stability of the closed protomers in the open state of the S-G614 mutant.

We also employed the FoldX approach with the all-atom representation of protein structure^99–102^ to evaluate residue-based folding free energy contributions in the D614 and G614 S trimers (Figure 7). The protein stability ΔΔG changes were computed by averaging the results of computations over 1,000 samples obtained from MD simulation trajectories.^103,104^ In this analysis, positive folding free energy contributions indicate stability weaknesses sites and highly negative folding free energy contributions characterized sites of local stability in the S protein. For the locked closed form of the D614 S trimer, the stability hotspots are distributed across multiple regions including the NTD residues, the RBD core and the S2 domain regions (Figure 7A). It is worth noting that the most prominent stability centers were found in the RBD core and the region near the hinge hotspot cluster (residues 591-600) that featured stability peaks for hinge sites S591, F592 and G593 (Figure 7A). In both closed and open forms of the D614 S trimer, the receptor binding motif (RBM) of the RBD region (residues 470-491) showed only marginal stability as the energetic plasticity of this region is required for functional interactions with ACE2.

**Figure 7.**
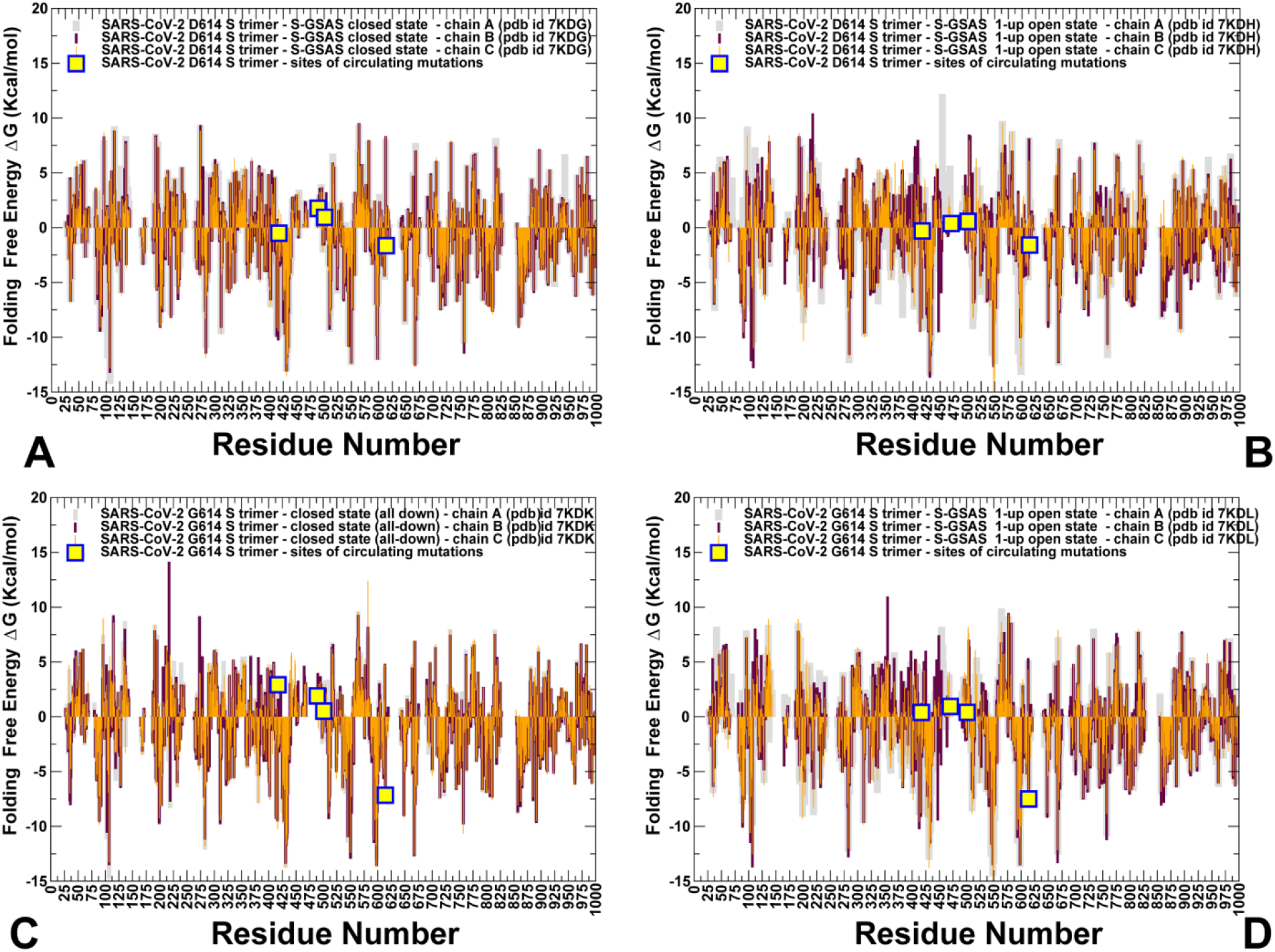
The residue-based folding stability analysis of the SARS-CoV-2 S-D614 and S-G614 trimers. The folding free energy profile for the S-D614 protein in the closed form (A), S-D614 protein in the 1-up open form (B), the S-G614 protein in the closed form (C) and S-G614 protein in the 1-up open form (D). The folding stability residue profiles are shown in color-coded bars (chain A is in grey bars, chain B in maroon bars, and chain C is in orange bars). The positions of the RBD sites K417, E484, N501, and D614/G614 that are subjected to circulating mutational variants are shown in yellow-colored filled squares.

In the open form of the D614 S trimer, the stability of the RBD core and S2 regions was partially reduced while maintaining the overall stabilizing signature in these positions (Figure 7B). By highlighting positions of circulating mutations K417, E484, N501 and D614 we noticed that D614 corresponds to a moderately stable position while other sites are marginally destabilizing (Figure 7A,B). Hence, mutations in these sites can be easily tolerated without impairing protein stability of the S trimer (Figure S1A) The overall stability pattern in the G614 S trimer remained preserved, highlighting similar regions of high structural stability (Figure 7C,D). However, the D614G mutation can incur a more significant stabilization in the site of mutation and in the neighboring regions, particularly enhancing stability of the hinge site cluster formed by F318, F592, G593, G614, K854, Y855, I856, and T859 residues (Figure 7C,D). Notably, the stability of G614 position is stronger in the open state of the S-G614 trimer (Figure 7D). Significantly, the open form of the G614 mutant showed a stronger stabilization of the RBD regions and S1-S2 inter-domain regions (Figure 7D) as compared the open form of the D614 S protein (Figure 7B).

Our findings support the notion that the D614G mutation in the SD2 domain could strengthen stability of the distal NTD and RBD regions in the open state^44,45^ and potentially promote exposure to the host receptor. These results offer support to the experimental observations that the enhanced stability of the S-D614G mutant may be linked with the mechanism of the reduced S1 shedding.^49^ Importantly, the differential stabilization of the S-G614 closed and open forms showing the energetic preferences of the mutant towards the open state may help to partly reconcile the “openness” and “S1-shedding” mechanism underpinning the D614G effects.

### Network Modeling and Community Analysis Suggest D614G-Induced Reorganization of the Residue Interaction Networks and Improved Allosteric Signaling in the Open States

Mechanistic network-based models allow for a quantitative analysis of allosteric molecular events in which conformational landscapes of protein systems can be remodeled by various perturbations such as mutations, ligand binding, or interactions with other proteins. Using this framework, the residue interaction networks in the SARS-CoV-2 spike trimer structures were built using a graph-based representation of protein structures in which residue nodes are interconnected through both dynamic^83^ and coevolutionary correlations.^84,85^ Using community decomposition, the residue interaction networks were divided into local interaction modules in which residues are densely interconnected and highly correlated, while the local communities are weakly coupled through long-range allosteric couplings. A community-based model of allosteric communications is based on the notion that groups of residues that form local interacting communities are correlated and switch their conformational states cooperatively. We explored community analysis as a network proxy for stability assessment. The number of communities for the S-D614 protein was greater in the closed form (Figure 8A) as compared to the open form (Figure 8B). In some contrast, the network analysis of the S-G614 mutant showed a subtle redistribution in the number and allocation of communities as the open state of the mutant harbored more communities (Figure 8C,D). Interestingly, the total number of local communities is moderately increased in both closed and open states of the S-G614 mutant. These results are consistent with the latest experimental data that demonstrated the improved stability of the D614G mutant as compared to the S-D614 protein allowing for reduction in a premature shedding of S1 domain.^49^

**Figure 8.**
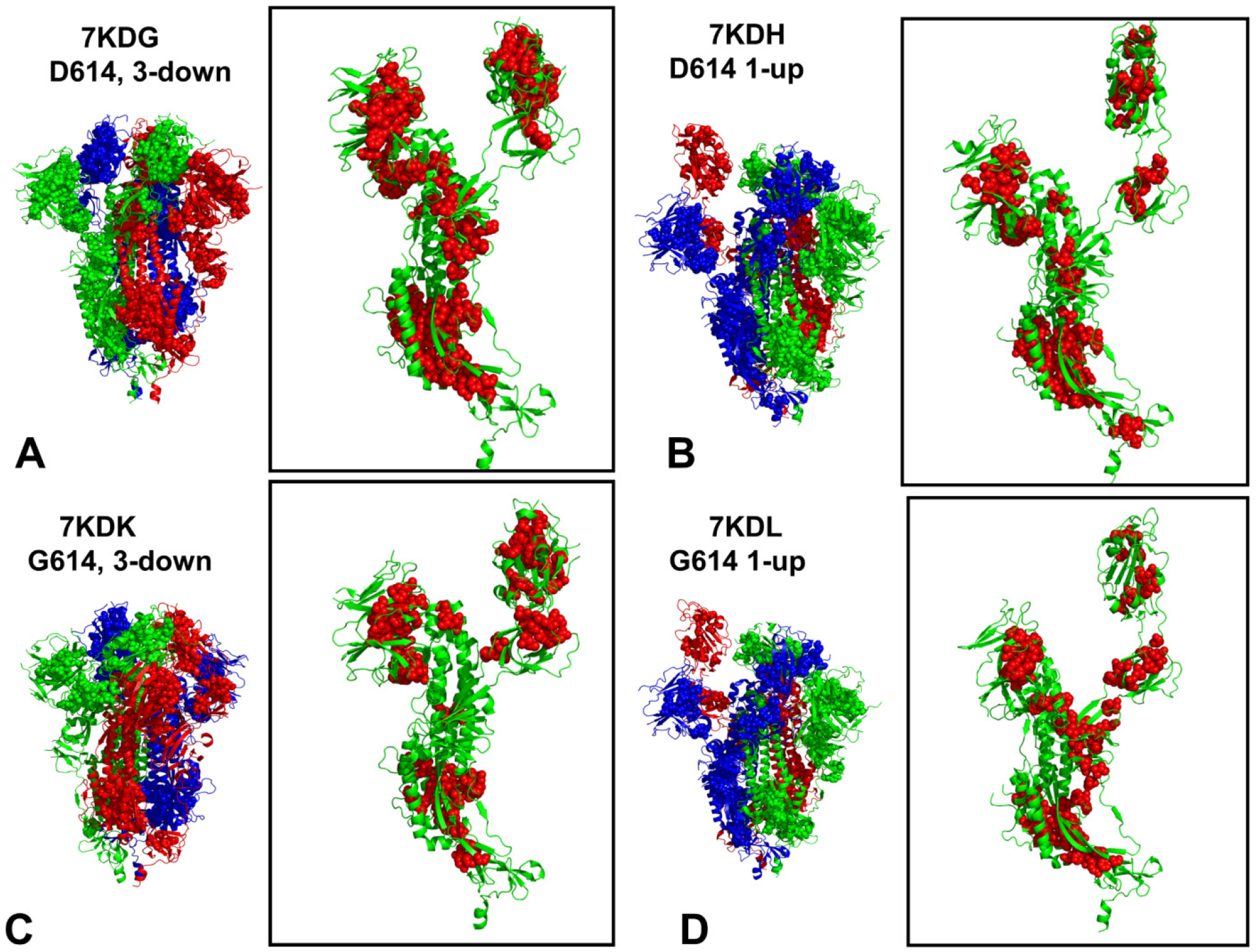
Community analysis and structural community maps in the SARS-CoV-2 S-D614 and S-D614G mutant structures. (A) Structural mapping of local communities is projected onto a single protomer for the S-GSAS/D614 in the closed-all down state (pdb id 7KDG). (B) Structural mapping of local communities for the S-GSAS/D614 in the open state (pdb id 7KDH). (C) Structural mapping of local communities in the closed form of S-GSAS/G614 (pdb id 7KDK). (D) Structural mapping of local communities for the S-GSAS/G614 in the open state (pdb id 7KDL). The cryo-EM structures of the SARS-CoV-2 S-D614 and S-G614 trimers are shown in ribbons (chain A in green, chain B in red and chain C in blue). The local communities are highlighted in spheres. The detailed close-ups of the local communities are shown for a single protomer with the protomer in green ribbons and communities in red spheres.

A more detailed inspection of local communities in the closed and partially open conformational states of the native S-D614 protein provided some important insight into the proposed mechanisms (Tables 2,3). We found that there are important changes in the communities situated in the RBD-CTD1, RBD-CTD2 and CTD1-CTD2 regions. The local modules in these regions that are unique to the closed form included Q314-S596-Q613, I693-V656-Y660, F543-L598-V576 and I666-L650-I670-T645 clusters (Table 2). In the partially open form, the state-specific communities in these regions included P579-P330-N544, F541-L552-I587 and R328-F543-P579 (Table 3). Characteristically, the key community in the closed form of the S-D614 protein is anchored by Q613, which is the immediate neighbor of D614, forming a tight stable cluster with Q314 in the NTD and S596 in CTD2. Structural mapping of local communities in the S-D614 states illustrated subtle differences in the distribution and density of stable modules (Figure 8A,B). A general comparison of structural maps indicated the better connectivity of local communities in the closed form of S-D614 forming a broad network linking the S2 regions with the NTD and RBD regions (Figure 8A,B). In addition, a number of unique communities are localized in the CTD2 region (I693-V656-Y660 and I666-L650-I670-T645), suggesting the stronger S1-S2 interfacial interactions and tighter packing between S1 and S2 domains in the closed form of the S-D614 protein. We also detected the larger number of communities in the NTD and RBD S1 regions in the closed form, indicating that the allosteric interaction network in the S1 and S1-S2 regions is stronger in the closed form, protecting all-down closed form in the S-D614 protein (Figure 8A). A weaker connectivity of local communities near S1-S2 interfaces and in the NTD/RBD regions was observed in the open state of S-D614 (Figure 8B). These findings suggested that efficiency of allosteric communication in the residue interaction network could favor the closed-down form of the S-D614 spike protein.

**Table 2.** The local interacting communities of the protomer A in the structure of SARS-CoV-2 S-GSAS/D614 in the closed state (pdb id 7KDG).

| Number | Contributing structural domains | Local Community Interacting Residues |
| --- | --- | --- |
| 1 | HR1-HR2 | A_1042_PHE A_1031_GLU A_1039_ARG |
| 2 | S2-HR1-HR2 | A_1067_TYR A_1049_LEU A_906_PHE A_909_ILE<br>A_911_VAL A_915_VAL |
| 3 | S2-HR1-HR2 | A_1054_GLN A_816_SER A_819_GLU |
| 4 | NTD | A_215_ASP A_266_TYR A_64_TRP |
| 5 | NTD | A_275_PHE A_290_ASP A_58_PHE |
| 6 | NTD | A_279_TYR A_44_ARG A_47_VAL A_49_HIS |
| 7 | CTD2-NTD | A_664_ILE A_312_ILE A_598_ILE |
| 8 | RBD | A_464_PHE A_355_ARG A_514_SER A_429_PHE<br>A_425_LEU A_426_PRO |
| 9 | RBD | A_509_ARG A_442_ASP A_438_SER |
| 10 | CTD2 | A_693_ILE A_656_VAL A_660_TYR |
| 11 | S2 | A_888_PHE A_789_TYR A_880_GLY |
| 12 | S2-HR1 | A_1001_LEU A_1005_GLN A_759_PHE |
| 13 | HR1-HR2 | A_1024_LEU A_1028_LYS A_1042_PHE A_1032_CYS |
| 14 | S2-HR1-HR2 | A_1062_PHE A_1029_MET A_1033_VAL A_1053_PRO<br>A_877_LEU |
| 15 | HR1-HR2 | A_1032_CYS A_1043_CYS A_1048_HIS A_1051_SER<br>A_1064_HIS |
| 16 | S2-HR1-HR2 | A_819_GLU A_1055_SER A_874_THR |
| 17 | HR1-HR2 | A_1075_PHE A_1096_VAL A_1110_TYR A_714_ILE |
| 18 | HR1-HR2 | A_1106_GLN A_1109_PHE A_915_VAL |
| 19 | NTD | A_231_ILE A_130_VAL A_168_PHE |
| 20 | NTD | A_193_VAL A_204_TYR A_37_TYR |
| 21 | NTD | A_285_ILE A_279_TYR A_38_TYR |
| 22 | NTD-CTD1-CTD2 | A_299_THR A_315_THR A_597_VAL |
| 23 | RBD | A_338_PHE A_342_PHE A_368_LEU |
| 24 | RBD | A_451_TYR A_401_VAL A_442_ASP |
| 25 | RBD | A_353_TRP A_398_ASP A_464_PHE |
| 26 | RBD-CTD1 | A_543_PHE A_585_LEU A_576_VAL |
| 27 | CTD2 | A_666_ILE A_650_LEU A_670_ILE A_645_THR |
| 28 | HR1 | A_977_LEU A_749_CYS A_993_ILE A_997_ILE |
| 29 | S2 | A_878_LEU A_806_LEU A_882_ILE |
| 30 | S2 | A_906_PHE A_923_ILE A_916_LEU |
| 31 | HR1 | A_1000_ARG A_977_LEU A_996_LEU |
| 32 | S2-HR1-HR2 | A_741_TYR A_1004_LEU A_962_LEU A_858_LEU |
| 33 | S2-HR1-HR2 | A_902_MET A_1050_MET A_898_PHE |
| 34 | RBD | A_342_PHE A_511_VAL A_374_PHE A_436_TRP |
| 35 | RBD | A_353_TRP A_400_PHE A_423_TYR A_512_VAL A_410_ILE |
| 36 | RBD | A_392_PHE A_515_PHE A_395_VAL |
| 37 | RBD-CTD2 | A_314_GLN A_596_SER A_613_GLN |

**Table 3.** The local interacting communities of the protomer A in the structure of SARS-CoV-2 S-GSAS/D614 in the open state (pdb id 7KDH).

| Number | Contributing structural domains | Local Community Interacting Residues |
| --- | --- | --- |
| 1 | S2-HR1 | A 731 MET A 1014 ARG A 955 ASN |
| 2 | NTD | A 101 ILE A 240 THR A 265 TYR A 92 PHE |
| 3 | S2-HR1-HR2 | A 1028 LYS A 1062 PHE A 727 LEU |
| 4 | HR1-HR2 | A 1042 PHE A 1031 GLU A 1039 ARG |
| 5 |  | A 1032 CYS A 1048 HIS A 1051 SER A 1064 HIS |
| 6 | S2-HR1-HR2 | A 1067 TYR A 1049 LEU A 909 ILE |
| 7 | S2-HR1-HR2 | A 1054 GLN A 816 SER A 819 GLU |
| 8 | HR1-HR2 | A 1102 TRP A 1081 ILE A 1135 ASN |
| 9 | NTD | A 238 PHE A 92 PHE A 267 VAL |
| 10 | NTD | A 275 PHE A 290 ASP A 58 PHE |
| 11 | NTKD | A 279 TYR A 44 ARG A 47 VAL A 49 HIS |
| 12 | NTD-CTD2 | A 299 THR A 315 THR A 597 VAL |
| 13 | CTD2 | A 664 ILE A 312 ILE A 598 ILE |
| 14 | RBD | A 418 ILE A 495 TYR A 453 TYR |
| 15 | RBD | A_464_PHE A_429_PHE A_425_LEU A_426_PRO<br>A 514 SER |
| 16 | RBD | A 454 ARG A 457 ARG A 467 ASP |
| 17 | HR1-HR2 | A 1062 PHE A 1029 MET A 1033 VAL |
| 18 | S2-HR1-HR2 | A_819_GLU A_1055_SER A_874_THR |
| 19 | HR1-HR2 | A 1106 GLN A 1109 PHE A 915 VAL |
| 20 | NTD | A 193 VAL A 204 TYR A 37 TYR |
| 21 | NTD | A 285 ILE A 279 TYR A 38 TYR |
| 22 | RBD-CTD1 | A 579 PRO A 330 PRO A 544 ASN |
| 23 | RBD | A 351 TYR A 454 ARG A 492 LEU |
| 24 | RBD-CTD1 | A 365 TYR A 387 LEU A 515 PHE A 432 CYS |
| 25 | CTD1 | A 541 PHE A 552 LEU A 587 ILE |
| 26 | S2-HR1 | A 997 ILE A 749 CYS A 993 ILE |
| 27 | S2 | A 898 PHE A 802 PHE A 797 PHE |
| 28 | S2 | A 878 LEU A 806 LEU A 882 ILE |
| 29 | S2 | A 818 ILE A 804 GLN A 935 GLN |
| 30 | S2-HR1 | A 923 ILE A 916 LEU A 906 PHE |
| 31 | S2-HR1 | A 905 ARG A 1050 MET A 898 PHE A 902 MET |
| 32 | S2-HR1-HR2 | A_1065_VAL A_1052_PHE A_802_PHE A_805_ILE<br>A 878 LEU A 927 PHE |
| 33 | RBD-CTD1 | A_328_ARG A_543_PHE A_579_PRO |
| 34 | RBD | A 342 PHE A 511 VAL A 374 PHE A 436 TRP |
| 35 | S2-HR1-HR2 | A 805 ILE A 1054 GLN A 818 ILE |

The analysis revealed the increased number of stable communities for the S-G614 mutant in both the closed form (Table 4) and partially open state (Table 5). Both forms of S-G614 mutant featured the appreciable number of communities in the CTD1 and CTD2 regions involved in stabilization of the S1-S2 interfaces (Figure 8C,D). These inter-domain local communities bridging S1 and S2 regions included T315-T299-V597, R328-S530-Q580, Q314-S596-Q613, I664-I312-I598, L611-P6645-L650, and F898-F802-F797. The stabilizing communities in the S1-S2 interfaces appeared to strengthen stability of both closed and open S-G614 forms which is consistent with the experimentally observed increased in protein stability of the D614G mutant.^49^ However, the balance in the number of communities may be shifted towards the open state of the S-G614 mutant, showing a more significant increase of local stable modules and promoting the preferential stabilization of the open mutant form. The key community near mutational site Q314-S596-Q613 is uniquely present only in the open form of the S-G614 mutant, likely pointing to state-specific rearrangements of stable interactions induced by the D614G mutation (Table 5). The important subtle rearrangements were found in the local communities localized in the N2R linker region (residues 306–334) that connects the NTD and RBD regions stacking against the CTD1 and CTD2 domains. Interestingly, the open form of the S-G614 mutant featured some of these modules (T299-T315-V597 and Q314-S596-Q613) and several additional N2R communities unique to this state (R328-F543-P579, P579-P330-N543, F338-F342-L368) that strengthen connections of the N2R linker with the NTD and RBD regions (Figure 8D, Table 5). This local subnetwork may be instrumental in bridging the individual domains in the S1 subunit and ensure a more efficient inter-connectivity between S2 regions, N2R and S1-RBD regions in the S-G614 mutant.

**Table 4.** The local interacting communities of the protomer A in the structure of SARS-CoV-2 S-GSAS/G614 in the closed state (pdb id 7KDK).

| Number | Contributing structural domains | Local Community Interacting Residues |
| --- | --- | --- |
| 1 | NTD | A 101 ILE A 190 ARG A 94 SER |
| 2 | HR1-HR2 | A 1042_PHE A 1031_GLU A 1037_SER A 1039_ARG<br>A 1032_CYS |
| 3 | S2-HR1-HR2 | A 1067_TYR A 1049_LEU A 906_PHE A 909_ILE A 911_VAL<br>A 915_VAL |
| 4 | S2-HR1-HR2 | A 1054_GLN A 816_SER A 819_GLU |
| 5 | S2-HR1-HR2 | A 1063_LEU A 724_THR A 934_ILE |
| 6 | S2-HR1-HR2 | A 1075_PHE A 1110_TYR A 714_ILE |
| 7 | NTD | A 265_TYR A 92_PHE A 240_THR |
| 8 | NTD | A 275_PHE A 290_ASP A 58_PHE |
| 9 | NTD | A 279_TYR A 44_ARG A 47_VAL A 49_HIS |
| 10 | NTD-CTD2 | A 315_THR A 299_THR A 597_VAL |
| 11 | NTD-RBD-CTD1 | A 328_ARG A 530_SER A 580_GLN |
| 12 | CTD1 | A 579_PRO A 544_ASN A 564_GLN |
| 13 | S2 | A 888_PHE A 789_TYR A 880_GLY |
| 14 | S2-HR1 | A 1001_LEU A 1005_GLN A 759_PHE |
| 15 | HR1-HR2 | A 1024_LEU A 1028_LYS A 1042_PHE |
| 16 | S2-HR1-HR2 | A 1062_PHE A 1029_MET A 1033_VAL A 1053_PRO<br>A 877_LEU |
| 17 | S2-HR1-HR2 | A 905_ARG A 1050_MET A 898_PHE A 901_GLN |
| 18 | S2-HR1-HR2 | A 819_GLU A 1055_SER A 874_THR |
| 19 | NTD | A 194_PHE A 238_PHE A 106_PHE A 117_LEU A 201_PHE<br>A 235_ILE A 86_PHE A 231_ILE A 90_VAL |
| 20 | NTD | A 193_VAL A 204_TYR A 37_TYR |
| 21 | NTD-RBD-CTD1 | A 328_ARG A 533_LEU A 578_ASP |
| 22 | RBD | A 338_PHE A 342_PHE A 368_LEU |
| 23 | RBD | A 353_TRP A 398_ASP A 464_PHE |
| 24 | RBD | A 365_TYR A 387_LEU A 515_PHE |
| 25 | RBD | A 406_GLU A 403_ARG A 495_TYR |
| 26 | RBD | A 509_ARG A 401_VAL A 451_TYR A 442_ASP A 438_SER<br>A 507_PRO |
| 27 | CTD1 | A 543_PHE A 585_LEU A 576_VAL |
| 28 | S2-HR1 | A 977_LEU A 749_CYS A 993_ILE A 997_ILE |
| 29 | S2 | A 878_LEU A 806_LEU A 882_ILE |
| 30 | HR1 | A 973_ILE A 984_LEU A 992_GLN |
| 31 | S2-HR1 | A 1000_ARG A 977_LEU A 996_LEU |
| 32 | S2-HR1 | A 741_TYR A 1004_LEU A 962_LEU A 858_LEU |
| 33 | HR1-HR2 | A 1065_VAL A 1052_PHE A 802_PHE A 927_PHE |
| 34 | NTD-RBD-CTD1 | A 328_ARG A 543_PHE A 579_PRO |
| 35 | RBD | A 342_PHE A 511_VAL A 374_PHE A 436_TRP A 347_PHE |
| 36 | RBD | A 353_TRP A 400_PHE A 423_TYR A 512_VAL A 410_ILE |
| 37 | NTD | A 104_TRP A 194_PHE A 238_PHE A 92_PHE |
| 38 | NTD-CTD2 | A 664_ILE A 312_ILE A 598_ILE |

**Table 5.** The local interacting communities of the protomer A in the structure of SARS-CoV-2 S-GSAS/G614 in the open state (pdb id 7KDL).

| Number | Contributing structural domains | Local Community Interacting Residues |
| --- | --- | --- |
| 1 | S2-HR1 | A_1001_LEU A_1005_GLN A_759_PHE |
| 2 | HR1-HR2 | A_1042_PHE A_1031_GLU A_1037_SER A_1039_ARG |
| 3 | S2-HR1-HR2 | A_1067_TYR A_1049_LEU A_906_PHE A_909_ILE<br>A_911_VAL |
| 4 | S2-HR1-HR2 | A_1054_GLN A_816_SER A_819_GLU |
| 5 | S2-HR1-HR2 | A_1075_PHE A_1110_TYR A_714_ILE |
| 6 | HR1-HR2 | A_1102_TRP A_1081_ILE A_1135_ASN |
| 7 | NTD | A_186_PHE A_264_ALA A_66_HIS |
| 8 | NTD | A_189_LEU A_210_ILE A_217_PRO |
| 9 | NTD | A_220_PHE A_288_ALA A_36_VAL |
| 10 | NTD | A_265_TYR A_92_PHE A_240_THR |
| 11 | NTD | A_265_TYR A_65_PHE A_82_PRO |
| 12 | NTD | A_275_PHE A_290_ASP A_58_PHE |
| 13 | NTD | A_279_TYR A_44_ARG A_49_HIS |
| 14 | NTD | A_299_THR A_315_THR A_597_VAL |
| 15 | CTD2 | A_664_ILE A_312_ILE A_598_ILE |
| 16 | RBD | A_342_PHE A_511_VAL A_374_PHE A_436_TRP A_347_PHE<br>A_399_SER A_509_ARG |
| 17 | RBD | A_351_TYR A_452_LEU A_454_ARG A_492_LEU |
| 18 | NTD | A_91_TYR A_35_GLY A_56_LEU |
| 19 | RBD | A_464_PHE A_429_PHE A_425_LEU A_426_PRO A_514_SER |
| 20 | RBD | A_451_TYR A_401_VAL A_497_PHE A_448_ASN |
| 21 | RBD | A_454_ARG A_457_ARG A_467_ASP |
| 22 | CTD2 | A_611_LEU A_666_ILE A_650_LEU |
| 23 | S2 | A_888_PHE A_789_TYR A_880_GLY |
| 24 | HR1-HR2 | A_1000_ARG A_977_LEU A_996_LEU |
| 25 | HR1-HR2 | A_1024_LEU A_1028_LYS A_1042_PHE A_1032_CYS |
| 26 | S2-HR1-HR2 | A_1062_PHE A_1029_MET A_1033_VAL A_1053_PRO<br>A_877_LEU |
| 27 | HR1-HR2 | A_1032_CYS A_1043_CYS A_1048_HIS A_1051_SER<br>A_1064_HIS |
| 28 | S2-HR1-HR2 | A_819_GLU A_1055_SER A_874_THR |
| 29 | HR1-HR2 | A_1115_ILE A_1104_VAL A_1119_ASN |
| 30 | S2-HR1-HR2 | A_1106_GLN A_1109_PHE A_915_VAL |
| 31 | NTD | A_193_VAL A_204_TYR A_37_TYR |
| 32 | NTD | A_104_TRP A_194_PHE A_238_PHE A_92_PHE A_267_VAL<br>A_84_LEU |
| 33 | NTD | A_238_PHE A_106_PHE A_117_LEU A_201_PHE A_235_ILE<br>A_86_PHE A_90_VAL |
| 34 | RBD-CTD1 | A_579_PRO A_330_PRO A_544_ASN |
| 35 | RBD | A_338_PHE A_342_PHE A_368_LEU |
| 36 | RBD | A_353_TRP A_398_ASP A_423_TYR A_464_PHE |
| 37 | RBD | A_365_TYR A_387_LEU A_515_PHE |
| 38 | CTD1 | A_543_PHE A_585_LEU A_576_VAL |
| 39 | S2-HR1 | A_997_ILE A_749_CYS A_993_ILE |
| 40 |  | A_878_LEU A_806_LEU A_882_ILE |
| 41 | S2 | A_906_PHE A_902_MET A_923_ILE A_916_LEU |
| 42 | S2-HR1 | A_741_TYR A_1004_LEU A_962_LEU A_858_LEU |
| 43 | S2-HR1-HR2 | A_905_ARG A_1050_MET A_898_PHE A_902_MET |
| 44 | HR1-HR2 | A_1065_VAL A_1052_PHE A_802_PHE A_927_PHE |
| 45 | HR1-HR2 | A_1081_ILE A_1115_ILE A_1137_VAL |
| 46 | NTD | A_285_ILE A_38_TYR A_279_TYR |
| 47 | NTD-RBD-CTD1 | A_328_ARG A_543_PHE A_579_PRO |
| 48 | S2 | A_898_PHE A_802_PHE A_797_PHE |
| 49 | RBD-CTD2 | A_314_GLN A_596_SER A_613_GLN |

The observed difference in the total number of local modules favoring the open mutant state is largely due emerging communities in the NTD and RBD regions. Indeed, while the open form of the S-D614 spike protein is characterized by mobile NTDs that could provide flexible access to the ACE2 receptor, the increased stabilization of the NTD and RBD regions in the S-G614 mutant may accompany strengthening of the S1-S2 interdomain interactions and could contribute to a decrease in premature shedding of S1 domain. Hence, D614G mutation may also exert allosteric effect by partly immobilizing the distal NTD and RBD regions. These findings support the previously suggested notion that the D614G mutation in the SD2 domain could allosterically strengthen stability of the distal NTD and RBD regions in the open state^44,45^ and therefore potentially promote exposure to the host receptor and greater infectivity.

Our results are more consistent with the latest experimental data^46^ by showing that the D614G mutation may strengthen the stability of the local community Q314-S596-Q613 and promote hydrogen bonding interactions between Q613 and T859 of adjacent protomer afforded by the local backbone flexibility at the mutational site. The presented results also partly reconciled several scenarios offered to explain functional effects of the D614G mutation. Indeed, the community network analysis suggested that preferential stabilization of the open mutant form can be determined by two main factors : (a) through strengthening of the S1-S2 interactions and (b) by reducing functional movements of the NTD and RBD regions exposed to binding with the host receptor. Accordingly, the D614G mutation may exert its effect through allosteric stabilization of the S1-S2 interfaces limiting shedding of the S1 domain and by reducing flexibility and enhancing thermodynamic preferences of the open state that is central aspect of the “openness”-based mechanistic scenario.

Hence, dynamic network modeling and community analysis of the S-D614 and S-G614 proteins revealed that D614G mutation can induce a partial rearrangement of the residue interaction networks and promote the larger number of stable communities in both the closed and open forms by enhancing the S1-S2 inter-domain interactions. Furthermore, the network analysis suggested a differential stabilization of the S-G614 mutant, favoring the open form in which strengthened allosteric couplings between mutational sites, S1-S2 regions and NTD/RBD regions in S1 could contribute to a decrease in premature shedding of S1 domain. Although recent studies indicated that D614G variant did not itself drive escape from antibody binding, it was found that D614G can remarkably potentiate escape mutations at some positions in certain patients, supporting an allosteric mechanism of action triggered by this mutation on dynamics and function in remote regions exposed to interactions with antibodies. In light of these newly emerging experimental data, our results argue that the D614G mutation may exert its global impact on other sites by acting as an important mediating center governing regulation of the SARS-CoV-2 machine.

## Conclusions

We combined several simulation-based approaches with dynamic network modeling and community analysis to quantify the effect of D614G mutation on dynamics, stability and network organization of the SARS-CoV-2 S proteins. The results of this study provide a novel insight into the molecular mechanisms underlying the effect of D614G mutation by examining SARS-CoV-2 S protein as an allosteric regulatory machine. Using mutational sensitivity analysis of the SARS-CoV-2 S-D614 and S-G614 proteins we demonstrated the improved stability of the D614G mutant as compared to the S-D614 protein, offering support to the reduced shedding mechanism underlying functional effects of the D614G circulating mutation. By examining the dynamic and network properties of the SARS-CoV-2 S trimer proteins, we characterized the distribution of allosteric hotspots in the S-D614 and S-G614 mutant structures revealing consolidation of the communication hotspots in the CTD1-CTD2 regions of the S-G614 mutant that may promote the greater stability and efficient allosteric couplings between S1 and S2 regions. A systematic characterization of stable local communities in multiple states of the S-D614 and S-G614 proteins, we demonstrated that D614G mutation can increase protein stability and strengthen the allosteric interaction networks in both closed and open forms. This study provides support to the reduced shedding hypothesis suggesting that D614G mutation can exert its primary effect through allosterically induced changes on stability and long-range communications in the residue interaction networks. Examining functions of the SARS-CoV-2 pike proteins through the prism of an allosterically regulated machine^105^ may prove to be useful to uncover functional mechanisms and rationalize the growing body of diverse experimental data.

## SUPPORTING INFORMATION

Figure S1 shows structures of the SARS-CoV-2 S-D614 trimer in the locked closed state, and SARS-CoV-2 S-G614 in the locked closed state, intermediate and 1 RBD-up open form. Figure S2 describes conformational mobility profiles for the SARS-CoV-2 S-D614 trimer in the locked closed state and SARS-CoV-2 S-G614 in the locked closed state, intermediate and 1 RBD-up open form. Figure S3 shows functional dynamics of these SARS-CoV-2 S-D614 and S-G614 trimer structures. Figure S4 highlights mutational sensitivity profiles of theses SARS-CoV-2 S-D614 and S-G614 trimer structures. Supporting information contains Tables S1-S4 that characterizes the inter-protomer contacts in the SARS-CoV-2 S-D614 and SARS-CoV-2 S-G614 structures in the closed and open states.

This material is available free of charge via the Internet at http://pubs.acs.org.

## AUTHOR INFORMATION

The authors declare no competing financial interest.

## Acknowledgment

This work was supported by the Kay Family Foundation Grant A20-0032.

## ABBREVIATIONS

SARS: Severe Acute Respiratory Syndrome
RBD: Receptor Binding Domain
ACE2: Angiotensin-Converting Enzyme 2 (ACE2)
NTD: N-terminal domain
RBD: receptor-binding domain
CTD1: C-terminal domain 1
CTD2: C-terminal domain 2
FP: fusion peptide
FPPR: fusion peptide proximal region
HR1: heptad repeat 1
CH: central helix region
CD: connector domain
HR2: heptad repeat 2
TM: transmembrane anchor
CT: cytoplasmic tail

## Notes

### Competing Interest Statement

The authors have declared no competing interest.

