## Supplemental Information for "Landscape-Based Mutational Sensitivity Cartography and Network Community Analysis of the SARS-CoV-2 Spike Protein Structures: Quantifying Functional Effects of the Circulating Variants"

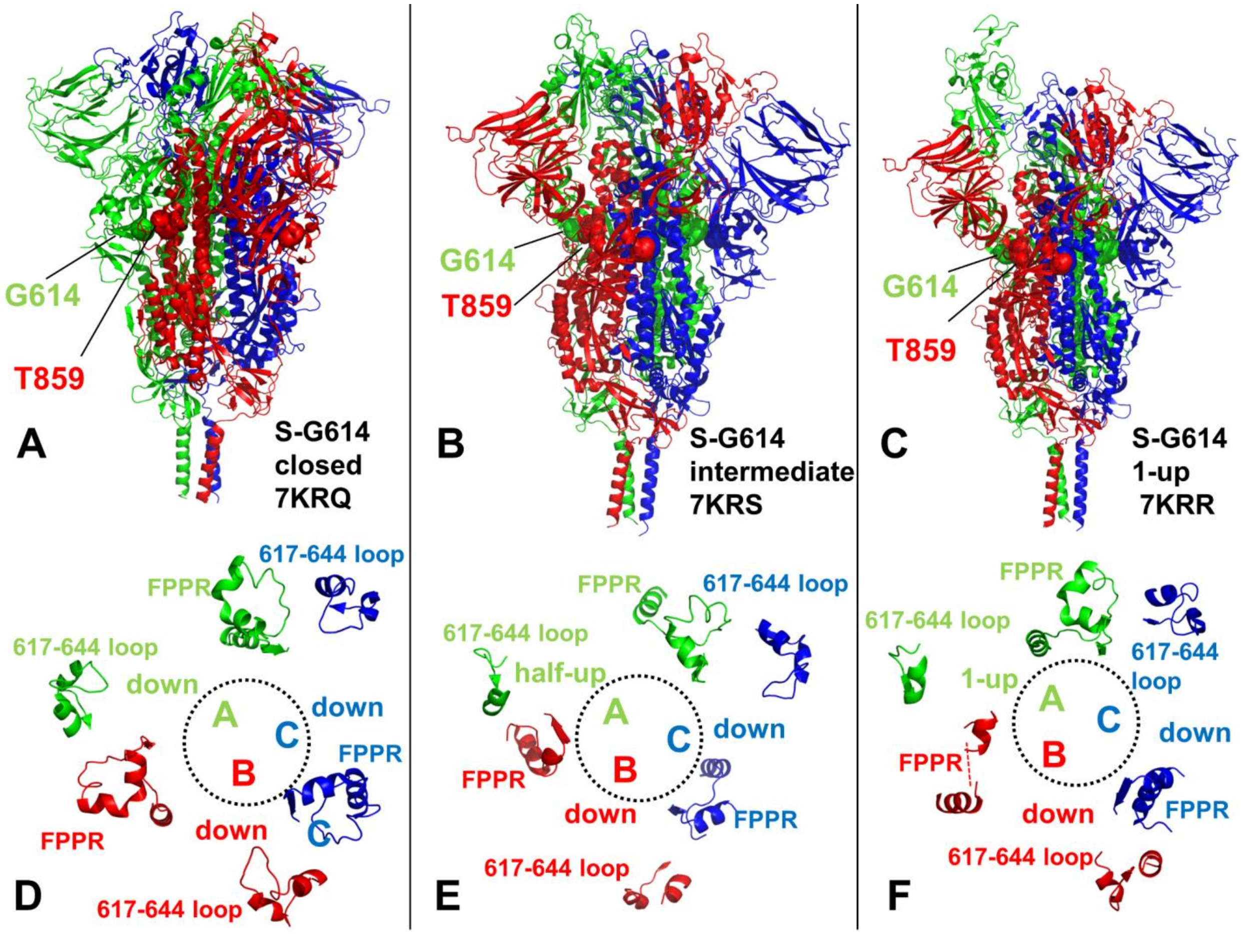


**Figure S1.** Cryo-EM structures of the SARS-CoV-2 S trimer structures used in this study. The cryo-EM structures of SARS-CoV-2 S-G614 in the closed state, pdb id 7KRQ (A), in the intermediate state, pdb id 7KRS (B) and 1 RBD-up open form, pdb id 7KRR (C). The structure is in ribbons with protomers A,B,C are colored in green, red and blue. The position of G614 and T859 are shown in spheres colored according to the chain. (D-F) Structures of the 630 loop (residues 617 to 644) and FPPR (residues 823 to 862) are shown for protomer A in green ribbons, protomer B in red ribbons and protomer C in blue ribbons.

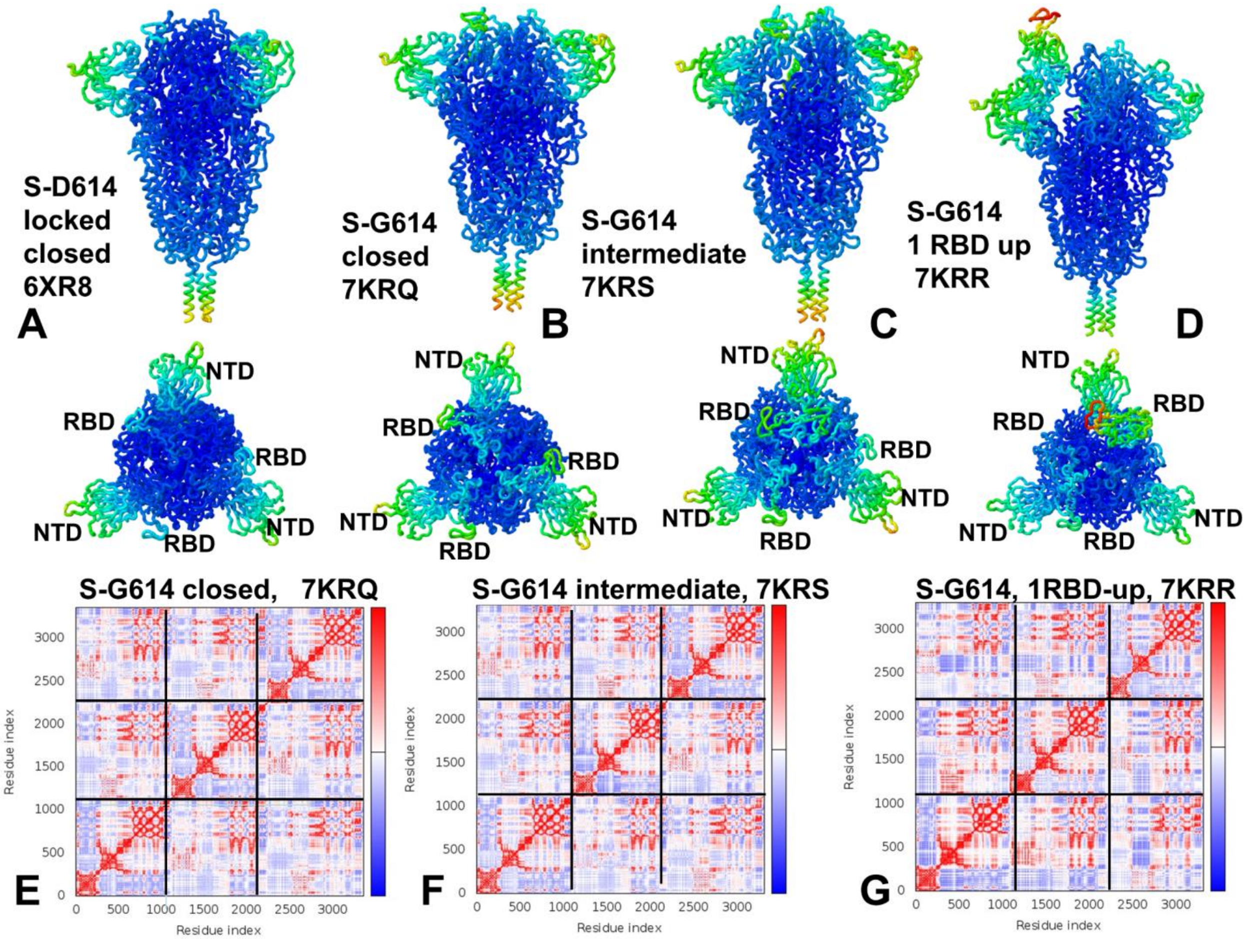


**Figure S2.** Conformational mobility profiles for the SARS-CoV-2 S-D614 trimer in the locked closed state, pdb id 6XR8 (A), SARS-CoV-2 S-G614 in the closed state, pdb id 7KRQ (A), in the intermediate state, pdb id 7KRS (B) and 1 RBD-up open form, pdb id 7KRR (C). The structures are in ribbons with the rigidity-to-flexibility scale colored from blue to red**.** Middle panel illustrates top views of the superposition of the three structures of the S-D614 and S-G614 structures. (E-G) The covariance matrix indicates coupling between pairs of residues. Cross-correlations of residue-based fluctuations vary between +1 (correlated motion; fluctuation vectors in the same direction, colored in dark red) and -1 (anti-correlated motions; fluctuation vectors in the same direction, colored in dark blue). The values > 0.5 are colored in dark red and the lower bound in the color bar indicates the value of the most anti-correlated pairs. The boxes highlight cross-correlations for the protomer A, protomer B and protomer C.

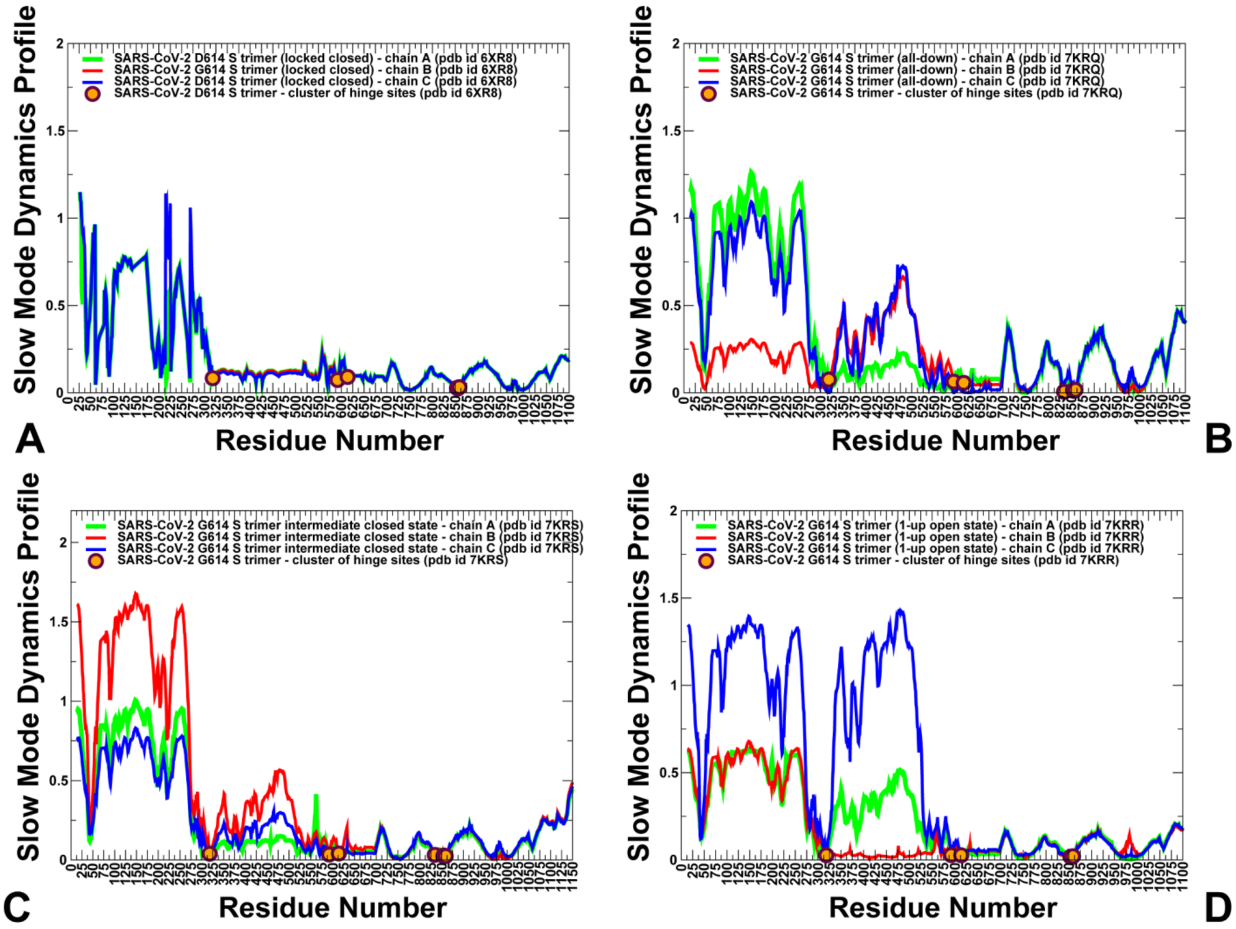


**Figure S3.** Functional dynamics of the SARS-CoV-2 S-D614 and S-G614 trimer structures. The reported essential mobility profiles were averaged over the first three major low frequency modes. (A) The essential mobility profiles for the SARS-CoV-2 S-D614 in the locked closed form (pdb id 6XR8). (B) The slow mode profile for the SARS-CoV-2 S-G614 trimer structure in the closed form (pdb id 7KRQ). (C) The slow mode profile for the SARS-CoV-2 S-G614 trimer in the intermediate closed form (pdb id 7KRS). (D) The slow mode profile for the SARS-CoV-2 S-G614 trimer structure in the 1 RBD-up open form (pdb id 7KRS). The profiles for protomer chains A, B and C are shown in green, red and blue lines, respectively. The positions of the hinge sites forming the inter-protomer cluster F318, F592, D614/G614, Y855, I856, and T859 are shown along the profiles in filled orange-colored circles.

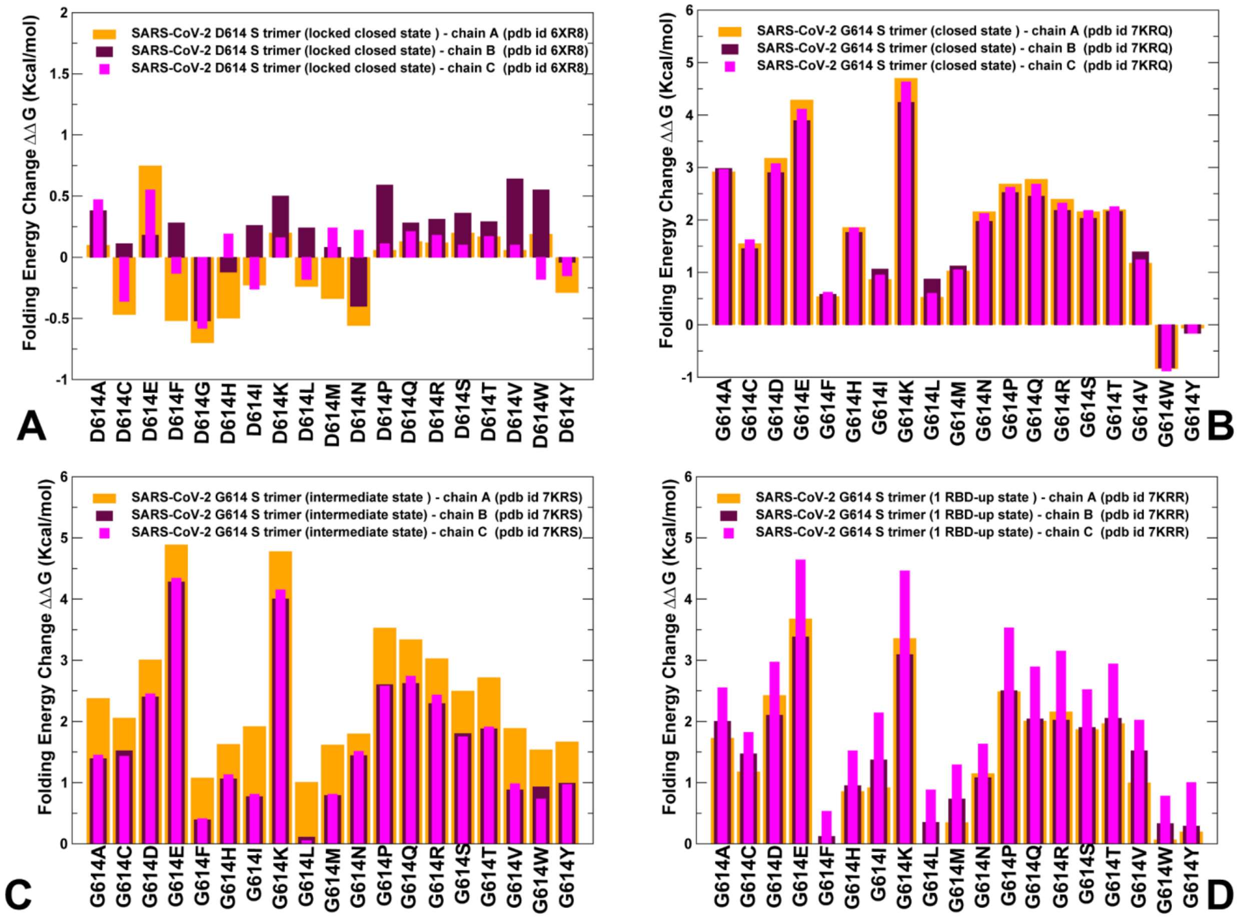


**Figure S4.** Mutational sensitivity analysis of the SARS-CoV-2 S-D614 and S-G614 trimers. Mutational sensitivity scanning of D614 position in the closed form of the S-D614 protein (A). Mutational sensitivity scanning of G614 position in the closed form of the S-G614 protein (B), intermediate state (C) and 1 RBD-up open state (D). The protein stability changes are shown for protomer A in orange-colored bars, for protomer B in maroon-colored bars, and for the protomer C in magenta-colored bars.

**Table S1.** The list of the inter-protomer contacts of the protomer A in the structure of SARS-CoV-2 S-GSAS/D614 in the closed state (pdb id 7KDG).

| S protein residue name | S protein residue number | S protein chain | S protein residue name | S protein residue number | S protein chain |
|---|---|---|---|---|---|
| PHE | 970 | A | GLN | 755 | B |
| ALA | 890 | A | VAL | 1068 | C |
| ASN | 919 | A | VAL | 1128 | C |
| MET | 900 | A | VAL | 1094 | C |
| MET | 740 | A | ARG | 319 | C |
| VAL | 705 | A | LEU | 894 | B |
| ILE | 569 | A | ASN | 960 | B |
| ASN | 907 | A | ARG | 1091 | C |
| GLN | 913 | A | GLY | 1093 | C |
| ARG | 466 | A | GLY | 232 | B |
| TYR | 369 | A | GLN | 414 | C |
| ALA | 1016 | A | ILE | 1013 | C |
| GLN | 779 | A | MET | 697 | C |
| SER | 758 | A | GLN | 965 | C |
| LEU | 864 | A | MET | 697 | C |
| ILE | 712 | A | PRO | 897 | B |
| PRO | 463 | A | GLY | 199 | B |
| ALA | 668 | A | PRO | 862 | B |
| GLY | 891 | A | LYS | 1045 | C |
| TYR | 707 | A | PHE | 898 | B |
| ALA | 890 | A | ASP | 1041 | C |
| LEU | 966 | A | ASP | 571 | C |
| ASN | 856 | A | THR | 572 | C |
| GLN | 613 | A | THR | 859 | B |
| SER | 758 | A | SER | 968 | C |
| ASN | 709 | A | PRO | 897 | B |
| SER | 45 | A | LYS | 557 | C |
| LEU | 546 | A | ASP | 979 | B |
| ILE | 670 | A | LEU | 864 | B |
| TYR | 505 | A | SER | 373 | B |
| ASN | 969 | A | GLN | 755 | B |
| VAL | 382 | A | LEU | 984 | B |
| THR | 1009 | A | ILE | 1013 | C |
| PRO | 897 | A | SER | 708 | C |
| PHE | 970 | A | GLY | 757 | B |

| | | | | | |
|---|---|---|---|---|---|
| LEU | 560 | A | GLY | 283 | B |
| TYR | 873 | A | LEU | 699 | C |
| GLY | 545 | A | ASN | 978 | B |
| ALA | 766 | A | GLN | 1010 | C |
| PHE | 855 | A | PHE | 592 | C |
| TYR | 1047 | A | ALA | 890 | B |
| PRO | 863 | A | GLY | 669 | C |
| GLU | 918 | A | GLY | 1124 | C |
| PHE | 855 | A | PRO | 589 | C |
| GLU | 988 | A | PRO | 384 | C |
| GLY | 889 | A | GLY | 1046 | C |
| ALA | 701 | A | GLN | 787 | B |
| TYR | 873 | A | GLY | 700 | C |
| VAL | 1068 | A | GLY | 891 | B |
| LYS | 386 | A | ARG | 983 | B |
| GLN | 1002 | A | TYR | 756 | B |
| GLN | 762 | A | GLN | 965 | C |
| LEU | 981 | A | LYS | 386 | C |
| LEU | 865 | A | MET | 697 | C |
| GLN | 1036 | A | VAL | 1040 | C |
| ALA | 892 | A | PRO | 1069 | C |
| GLY | 798 | A | ILE | 1130 | C |
| GLN | 314 | A | LEU | 861 | B |
| TYR | 873 | A | SER | 698 | C |
| PHE | 1089 | A | ASN | 914 | B |
| GLU | 773 | A | GLU | 1017 | C |
| GLN | 52 | A | LEU | 754 | B |
| ASP | 427 | A | VAL | 987 | C |
| ARG | 1019 | A | ALA | 1016 | C |
| LEU | 1145 | A | LEU | 1145 | B |
| TYR | 1047 | A | THR | 887 | B |
| ASP | 1041 | A | LEU | 1034 | B |
| GLY | 889 | A | ASP | 1041 | C |
| GLY | 999 | A | PHE | 759 | B |
| LEU | 390 | A | SER | 982 | B |
| PRO | 1079 | A | TYR | 917 | B |
| GLN | 872 | A | LEU | 699 | C |
| ALA | 372 | A | ASP | 405 | C |
| GLN | 913 | A | PRO | 1079 | C |
| GLU | 918 | A | VAL | 1128 | C |
| PHE | 43 | A | PHE | 559 | C |

| | | | | | |
|---|---|---|---|---|---|
| ASN | 703 | A | ILE | 788 | B |
| ASP | 1041 | A | GLY | 889 | B |
| SER | 884 | A | VAL | 705 | C |
| ASP | 571 | A | LEU | 966 | B |
| GLU | 465 | A | GLY | 232 | B |
| ILE | 1013 | A | ILE | 1013 | C |
| LEU | 864 | A | ILE | 670 | C |
| ALA | 668 | A | THR | 866 | B |
| PHE | 565 | A | LYS | 41 | B |
| GLY | 669 | A | PRO | 863 | B |
| SER | 735 | A | GLN | 314 | C |
| GLY | 757 | A | PHE | 970 | C |
| GLN | 895 | A | SER | 708 | C |
| VAL | 42 | A | GLY | 566 | C |
| LYS | 557 | A | PHE | 43 | B |
| GLY | 891 | A | GLY | 1046 | C |
| TYR | 873 | A | MET | 697 | C |
| ARG | 567 | A | PHE | 43 | B |
| GLN | 563 | A | PHE | 43 | B |
| PHE | 592 | A | LEU | 858 | B |
| ASP | 40 | A | PHE | 562 | C |
| THR | 167 | A | ARG | 466 | C |
| LEU | 518 | A | LYS | 41 | B |
| ARG | 567 | A | ASP | 979 | B |
| SER | 1123 | A | ASN | 914 | B |
| TYR | 904 | A | GLY | 1093 | C |
| ALA | 890 | A | TYR | 1067 | C |
| VAL | 1128 | A | GLU | 918 | B |
| ARG | 466 | A | THR | 167 | B |
| PHE | 970 | A | ASP | 994 | B |
| ILE | 1013 | A | ARG | 1019 | B |
| LEU | 894 | A | ARG | 1107 | C |
| ARG | 567 | A | VAL | 976 | B |
| LEU | 861 | A | GLN | 314 | C |
| PRO | 230 | A | TYR | 396 | C |
| TYR | 707 | A | ILE | 794 | B |
| TYR | 707 | A | ASP | 796 | B |
| GLU | 1072 | A | ALA | 892 | B |
| GLN | 895 | A | ILE | 712 | C |
| PRO | 897 | A | ASN | 709 | C |
| ASN | 1074 | A | GLN | 895 | B |

| | | | | | |
|---|---|---|---|---|---|
| ARG | 403 | A | ALA | 372 | B |
| MET | 900 | A | THR | 1077 | C |
| GLN | 564 | A | LYS | 41 | B |
| MET | 740 | A | PRO | 589 | C |
| GLY | 669 | A | THR | 866 | B |
| THR | 1009 | A | THR | 1009 | B |
| ASP | 405 | A | ALA | 372 | B |
| GLN | 957 | A | ARG | 765 | B |
| SER | 968 | A | GLY | 757 | B |
| TRP | 886 | A | TYR | 1047 | C |
| ILE | 666 | A | LEU | 864 | B |
| GLY | 971 | A | ASP | 994 | B |
| SER | 982 | A | GLY | 545 | C |
| TYR | 38 | A | GLN | 563 | C |
| ALA | 570 | A | SER | 975 | B |
| TYR | 38 | A | LEU | 560 | C |
| GLN | 755 | A | SER | 968 | C |
| PRO | 1069 | A | LEU | 894 | B |
| GLY | 283 | A | LEU | 560 | C |
| PHE | 562 | A | VAL | 42 | B |
| ILE | 468 | A | GLN | 115 | B |
| GLN | 563 | A | GLY | 283 | B |
| PHE | 559 | A | PHE | 43 | B |
| SER | 698 | A | TYR | 873 | B |
| TYR | 707 | A | ILE | 896 | B |
| ILE | 896 | A | ILE | 712 | C |
| LEU | 560 | A | PHE | 43 | B |
| ARG | 983 | A | VAL | 382 | C |
| ARG | 983 | A | LYS | 386 | C |
| PHE | 759 | A | SER | 1003 | C |
| THR | 912 | A | PHE | 1121 | C |
| THR | 883 | A | VAL | 705 | C |
| LYS | 41 | A | HIS | 519 | C |
| LYS | 202 | A | LEU | 518 | C |
| ILE | 233 | A | PHE | 464 | C |
| GLN | 784 | A | ASP | 1041 | C |
| VAL | 42 | A | PHE | 562 | C |
| LYS | 786 | A | GLY | 700 | C |
| LEU | 984 | A | TYR | 380 | C |
| GLU | 1017 | A | GLU | 773 | B |
| GLN | 1010 | A | LEU | 1012 | B |

| | | | | | |
|---|---|---|---|---|---|
| LYS | 986 | A | ASP | 427 | B |
| GLY | 1046 | A | GLY | 891 | B |
| LYS | 964 | A | SER | 758 | B |
| MET | 869 | A | GLY | 669 | C |
| ARG | 1091 | A | ASN | 907 | B |
| ILE | 569 | A | VAL | 963 | B |
| ILE | 896 | A | TYR | 707 | C |
| ASN | 907 | A | ARG | 1107 | C |
| SER | 968 | A | SER | 758 | B |
| MET | 697 | A | LEU | 864 | B |
| VAL | 987 | A | PRO | 412 | B |
| PHE | 562 | A | LYS | 41 | B |
| TYR | 396 | A | PRO | 230 | B |
| LEU | 865 | A | LEU | 699 | C |
| SER | 968 | A | ASP | 571 | C |
| VAL | 1008 | A | GLN | 1010 | C |
| GLU | 918 | A | VAL | 1129 | C |
| ARG | 1019 | A | ILE | 1013 | C |
| ARG | 983 | A | GLY | 381 | C |
| GLN | 913 | A | PHE | 1121 | C |
| TYR | 380 | A | LEU | 984 | B |
| ILE | 468 | A | LYS | 113 | B |
| GLY | 971 | A | TYR | 756 | B |
| GLN | 787 | A | ALA | 701 | C |
| ASN | 709 | A | ASP | 796 | B |
| TYR | 789 | A | ALA | 701 | C |
| ASN | 764 | A | GLN | 314 | C |
| ARG | 466 | A | GLN | 115 | B |
| LYS | 790 | A | SER | 704 | C |
| GLN | 613 | A | LEU | 861 | B |
| GLY | 880 | A | VAL | 705 | C |
| PHE | 592 | A | PHE | 855 | B |
| ALA | 647 | A | PRO | 862 | B |
| ILE | 882 | A | TYR | 707 | C |
| TYR | 789 | A | ASN | 703 | C |
| GLU | 516 | A | ASP | 228 | B |
| THR | 415 | A | TYR | 369 | B |
| ASP | 979 | A | ARG | 567 | C |
| GLN | 1002 | A | LEU | 1001 | B |
| SER | 375 | A | ARG | 408 | C |
| SER | 968 | A | GLN | 755 | B |

| | | | | | |
|---|---|---|---|---|---|
| VAL | 1128 | A | ASN | 919 | B |
| GLY | 566 | A | ARG | 44 | B |
| VAL | 1040 | A | SER | 1030 | B |
| MET | 697 | A | MET | 869 | B |
| GLY | 199 | A | PRO | 463 | C |
| ILE | 1013 | A | ALA | 1016 | B |
| TYR | 756 | A | GLN | 965 | C |
| LYS | 41 | A | PHE | 562 | C |
| ASN | 282 | A | LEU | 560 | C |
| ALA | 570 | A | SER | 967 | B |
| TYR | 707 | A | PRO | 897 | B |
| THR | 866 | A | GLY | 669 | C |
| MET | 740 | A | PHE | 592 | C |
| PHE | 43 | A | GLY | 566 | C |
| ASP | 568 | A | PHE | 855 | B |
| ASP | 1041 | A | SER | 1030 | B |
| ALA | 713 | A | ALA | 893 | B |
| GLN | 895 | A | ASN | 1074 | C |
| PHE | 1121 | A | ASN | 914 | B |
| TYR | 200 | A | ARG | 355 | C |
| ARG | 319 | A | ASP | 745 | B |
| GLY | 381 | A | ARG | 983 | B |
| ILE | 794 | A | TYR | 707 | C |
| ALA | 668 | A | LEU | 864 | B |
| PHE | 562 | A | ASP | 40 | B |
| GLN | 314 | A | THR | 768 | B |
| GLY | 381 | A | ILE | 973 | B |
| GLY | 381 | A | LEU | 984 | B |
| ASP | 571 | A | VAL | 976 | B |
| LEU | 1034 | A | VAL | 1040 | C |
| LEU | 1141 | A | LEU | 1141 | C |
| ASP | 571 | A | SER | 975 | B |
| GLN | 762 | A | GLN | 1010 | C |
| ASP | 1041 | A | ALA | 890 | B |
| ILE | 569 | A | VAL | 47 | B |
| ILE | 569 | A | LYS | 964 | B |
| TYR | 200 | A | SER | 514 | C |
| VAL | 1040 | A | GLU | 1031 | B |
| PRO | 862 | A | ALA | 647 | C |
| ASN | 703 | A | GLN | 787 | B |
| TYR | 789 | A | VAL | 705 | C |

| | | | | | |
|---|---|---|---|---|---|
| GLN | 762 | A | TYR | 1007 | C |
| GLY | 667 | A | PRO | 863 | B |
| ASP | 40 | A | HIS | 519 | C |
| ASP | 198 | A | ASP | 428 | C |
| THR | 961 | A | GLN | 762 | B |
| PHE | 464 | A | TYR | 200 | B |
| ASN | 856 | A | ALA | 570 | C |
| TYR | 756 | A | GLY | 971 | C |
| TYR | 1067 | A | ALA | 890 | B |
| GLY | 700 | A | ILE | 788 | B |
| ALA | 701 | A | ILE | 788 | B |
| SER | 982 | A | THR | 547 | C |
| SER | 373 | A | TYR | 505 | C |
| GLN | 755 | A | GLY | 971 | C |
| ASN | 317 | A | THR | 739 | B |
| THR | 739 | A | ARG | 319 | C |
| THR | 1009 | A | THR | 1009 | C |
| GLY | 548 | A | ASN | 978 | B |
| PHE | 1095 | A | MET | 900 | B |
| LYS | 417 | A | ASN | 370 | B |
| LYS | 41 | A | LEU | 518 | C |
| ILE | 233 | A | ARG | 466 | C |
| GLY | 669 | A | MET | 869 | B |
| MET | 869 | A | LEU | 699 | C |
| SER | 514 | A | TYR | 200 | B |
| PRO | 230 | A | ARG | 357 | C |
| ARG | 319 | A | THR | 739 | B |
| GLN | 965 | A | GLY | 757 | B |
| ASP | 427 | A | LYS | 986 | C |
| LYS | 558 | A | ASN | 282 | B |
| PRO | 589 | A | PHE | 855 | B |
| GLY | 667 | A | LEU | 864 | B |
| LEU | 984 | A | GLY | 381 | C |
| PHE | 759 | A | GLN | 965 | C |
| VAL | 987 | A | GLY | 413 | B |
| LEU | 1141 | A | GLU | 1144 | B |
| LYS | 786 | A | LYS | 1045 | C |
| GLY | 566 | A | PHE | 43 | B |
| THR | 961 | A | SER | 758 | B |
| LYS | 1045 | A | ALA | 890 | B |
| GLY | 283 | A | GLN | 563 | C |

| | | | | | |
|---|---|---|---|---|---|
| GLY | 232 | A | ARG | 466 | C |
| TYR | 707 | A | PHE | 797 | B |
| THR | 415 | A | ASP | 985 | C |
| LYS | 557 | A | SER | 45 | B |
| VAL | 42 | A | HIS | 519 | C |
| GLY | 283 | A | LYS | 558 | C |
| ASN | 914 | A | SER | 1123 | C |
| TYR | 505 | A | ALA | 372 | B |
| TRP | 886 | A | ARG | 1107 | C |
| LEU | 864 | A | CYS | 662 | C |
| LYS | 854 | A | ASP | 568 | C |
| ARG | 408 | A | TYR | 369 | B |
| GLU | 516 | A | LYS | 41 | B |
| GLY | 700 | A | LYS | 786 | B |
| ASP | 985 | A | SER | 383 | C |
| TYR | 904 | A | ARG | 1107 | C |
| LYS | 386 | A | ASP | 985 | B |
| THR | 696 | A | MET | 869 | B |
| ARG | 1039 | A | ARG | 1039 | B |
| PRO | 897 | A | SER | 711 | C |
| PHE | 43 | A | ARG | 567 | C |
| TYR | 917 | A | ILE | 1130 | C |
| MET | 900 | A | PHE | 1095 | C |
| PHE | 562 | A | TYR | 38 | B |
| ASN | 196 | A | LYS | 462 | C |
| PHE | 43 | A | PHE | 565 | C |
| GLU | 516 | A | TYR | 200 | B |
| VAL | 705 | A | GLN | 895 | B |
| ARG | 983 | A | LEU | 390 | C |
| PHE | 1121 | A | GLN | 913 | B |
| VAL | 382 | A | ARG | 983 | B |
| GLY | 413 | A | ASP | 985 | C |
| ASP | 994 | A | GLY | 971 | C |
| LYS | 462 | A | ASN | 196 | B |
| LYS | 1045 | A | GLY | 891 | B |
| GLU | 702 | A | TYR | 789 | B |
| ARG | 983 | A | LEU | 517 | C |
| SER | 711 | A | GLN | 895 | B |
| LEU | 560 | A | THR | 284 | B |
| ASP | 737 | A | ASN | 317 | C |
| VAL | 976 | A | ASP | 571 | C |

| | | | | | |
|---|---|---|---|---|---|
| ASP | 796 | A | SER | 708 | C |
| SER | 975 | A | ASP | 571 | C |
| ASP | 40 | A | GLN | 563 | C |
| ILE | 1013 | A | ILE | 1013 | B |
| LYS | 558 | A | GLY | 283 | B |
| SER | 711 | A | PRO | 897 | B |
| GLN | 895 | A | VAL | 705 | C |
| LEU | 560 | A | ASN | 282 | B |
| GLN | 787 | A | GLU | 702 | C |
| TYR | 396 | A | ASP | 228 | B |
| THR | 385 | A | THR | 415 | C |
| ALA | 701 | A | TYR | 789 | B |
| LYS | 790 | A | ASN | 703 | C |
| ALA | 890 | A | TYR | 1047 | C |
| ASP | 198 | A | PRO | 426 | C |
| LEU | 966 | A | ALA | 570 | C |
| ASN | 317 | A | ASP | 737 | B |
| PRO | 1069 | A | ALA | 892 | B |
| ASP | 228 | A | GLU | 516 | C |
| ARG | 1107 | A | TRP | 886 | B |
| ARG | 355 | A | TYR | 200 | B |
| GLU | 1111 | A | SER | 1123 | C |
| ARG | 1107 | A | ASN | 907 | B |
| GLU | 132 | A | ILE | 468 | C |
| ASN | 703 | A | TYR | 789 | B |
| LEU | 1012 | A | ILE | 1013 | C |
| GLU | 988 | A | CYS | 379 | C |
| PHE | 565 | A | PHE | 43 | B |
| ARG | 765 | A | GLN | 957 | C |
| SER | 383 | A | ARG | 983 | B |
| TYR | 369 | A | THR | 415 | C |
| THR | 768 | A | GLN | 314 | C |
| GLU | 1031 | A | PHE | 1042 | C |
| LYS | 854 | A | PHE | 592 | C |
| PRO | 1069 | A | GLY | 891 | B |
| THR | 859 | A | PHE | 592 | C |
| GLU | 702 | A | LYS | 790 | B |
| ALA | 713 | A | LEU | 894 | B |
| TYR | 756 | A | PHE | 970 | C |
| SER | 383 | A | GLU | 988 | B |
| ASN | 370 | A | LYS | 417 | C |

| | | | | | |
|---|---|---|---|---|---|
| GLU | 918 | A | PHE | 1089 | C |
| GLN | 563 | A | ARG | 44 | B |
| LEU | 390 | A | ARG | 983 | B |
| GLN | 965 | A | PHE | 759 | B |
| CYS | 671 | A | LEU | 864 | B |
| ASN | 914 | A | PHE | 1121 | C |
| PRO | 1079 | A | MET | 900 | B |
| LYS | 1038 | A | GLN | 1036 | B |
| GLU | 988 | A | VAL | 382 | C |
| ILE | 896 | A | SER | 711 | C |
| PHE | 759 | A | GLN | 1002 | C |
| LEU | 699 | A | ILE | 870 | B |
| GLN | 314 | A | SER | 735 | B |
| VAL | 705 | A | THR | 883 | B |
| PHE | 592 | A | THR | 859 | B |
| LYS | 854 | A | ILE | 569 | C |
| SER | 735 | A | GLN | 613 | C |
| VAL | 1040 | A | GLY | 889 | B |
| CYS | 662 | A | LEU | 864 | B |
| LEU | 864 | A | ALA | 668 | C |
| PRO | 792 | A | TYR | 707 | C |
| GLY | 757 | A | SER | 968 | C |
| ASP | 389 | A | SER | 982 | B |
| LYS | 386 | A | LEU | 984 | B |
| LEU | 865 | A | ALA | 668 | C |
| ARG | 1107 | A | LEU | 894 | B |
| LEU | 984 | A | SER | 383 | C |
| VAL | 1128 | A | TYR | 917 | B |
| ILE | 666 | A | PRO | 862 | B |
| LEU | 864 | A | ILE | 666 | C |
| MET | 869 | A | MET | 697 | C |
| PHE | 898 | A | TYR | 707 | C |
| LYS | 41 | A | GLN | 564 | C |
| GLU | 988 | A | LYS | 378 | C |
| ARG | 1019 | A | GLU | 1017 | C |
| ARG | 1091 | A | GLN | 913 | B |
| ILE | 973 | A | GLY | 381 | C |
| ASP | 198 | A | PRO | 463 | C |
| VAL | 42 | A | GLN | 563 | C |
| PHE | 759 | A | GLY | 999 | C |
| PRO | 463 | A | ASP | 198 | B |

| | | | | | |
|---|---|---|---|---|---|
| LEU | 1145 | A | GLU | 1144 | B |
| THR | 1077 | A | PRO | 897 | B |
| PHE | 759 | A | THR | 1006 | C |
| ARG | 1107 | A | TYR | 904 | B |
| VAL | 42 | A | PHE | 565 | C |
| LYS | 386 | A | SER | 982 | B |
| SER | 1030 | A | VAL | 1040 | C |
| PRO | 862 | A | GLY | 667 | C |
| PHE | 592 | A | MET | 740 | B |
| MET | 697 | A | LEU | 865 | B |
| ILE | 788 | A | GLY | 700 | C |
| GLU | 1072 | A | LEU | 894 | B |
| LEU | 699 | A | LEU | 865 | B |
| ASN | 370 | A | GLY | 416 | C |
| ILE | 714 | A | LEU | 894 | B |
| PRO | 230 | A | ARG | 355 | C |
| ASP | 198 | A | PHE | 464 | C |
| THR | 887 | A | TYR | 1047 | C |
| ASN | 856 | A | ASP | 568 | C |
| LEU | 894 | A | ALA | 713 | C |
| LYS | 790 | A | GLU | 702 | C |
| GLU | 1031 | A | ARG | 1039 | C |
| ILE | 1130 | A | GLN | 920 | B |
| ALA | 899 | A | TYR | 707 | C |
| GLN | 1005 | A | GLN | 1002 | C |
| GLU | 1144 | A | LEU | 1141 | C |
| PRO | 897 | A | ILE | 712 | C |
| ALA | 1016 | A | ARG | 1019 | B |
| VAL | 705 | A | TYR | 789 | B |
| ALA | 893 | A | ALA | 713 | C |
| ASN | 710 | A | PRO | 897 | B |
| VAL | 785 | A | LEU | 699 | C |
| LEU | 894 | A | GLU | 1072 | C |
| VAL | 705 | A | ALA | 879 | B |
| SER | 758 | A | LYS | 964 | C |
| ASN | 1108 | A | TRP | 886 | B |
| PHE | 970 | A | PHE | 759 | B |
| ALA | 890 | A | LYS | 1045 | C |
| GLN | 1010 | A | ALA | 766 | B |
| GLU | 702 | A | GLN | 787 | B |
| GLN | 1010 | A | GLN | 762 | B |

| | | | | | |
|---|---|---|---|---|---|
| SER | 1030 | A | ASP | 1041 | C |
| ARG | 355 | A | PRO | 230 | B |
| ILE | 569 | A | SER | 967 | B |
| GLN | 314 | A | ASN | 764 | B |
| TYR | 200 | A | PHE | 464 | C |
| GLN | 613 | A | SER | 735 | B |
| TYR | 707 | A | SER | 884 | B |
| ALA | 890 | A | PRO | 1069 | C |
| SER | 711 | A | ILE | 896 | B |
| ALA | 372 | A | TYR | 505 | C |
| LYS | 786 | A | ALA | 701 | C |
| ILE | 569 | A | LYS | 854 | B |
| ARG | 319 | A | MET | 740 | B |
| PHE | 1089 | A | TYR | 917 | B |
| THR | 385 | A | ASP | 985 | B |
| GLN | 913 | A | PRO | 1090 | C |
| THR | 1006 | A | GLN | 762 | B |
| THR | 883 | A | TYR | 707 | C |
| ILE | 1130 | A | LYS | 921 | B |
| ALA | 892 | A | GLU | 1072 | C |
| TYR | 789 | A | GLU | 702 | C |
| GLN | 755 | A | PHE | 970 | C |
| VAL | 963 | A | ALA | 570 | C |
| MET | 900 | A | ILE | 712 | C |
| GLU | 702 | A | ILE | 788 | B |
| ALA | 372 | A | ARG | 403 | C |
| ASN | 703 | A | LYS | 790 | B |
| GLN | 1005 | A | THR | 1006 | C |
| ASP | 985 | A | LYS | 386 | C |
| PHE | 1042 | A | GLU | 1031 | B |
| TYR | 917 | A | PHE | 1089 | C |
| GLY | 416 | A | TYR | 369 | B |
| GLN | 895 | A | ALA | 713 | C |
| TYR | 707 | A | PRO | 792 | B |
| ILE | 788 | A | GLU | 702 | C |
| ARG | 44 | A | GLN | 563 | C |
| THR | 547 | A | ASN | 978 | B |
| THR | 572 | A | ASN | 856 | B |
| GLU | 1031 | A | VAL | 1040 | C |
| PRO | 225 | A | PHE | 562 | C |
| SER | 383 | A | ASP | 985 | B |

| | | | | | |
|---|---|---|---|---|---|
| LYS | 378 | A | GLU | 988 | B |
| VAL | 987 | A | GLN | 414 | B |
| THR | 415 | A | THR | 385 | B |
| LYS | 786 | A | LEU | 699 | C |
| TYR | 917 | A | PRO | 1079 | C |
| GLN | 965 | A | SER | 758 | B |
| PHE | 759 | A | PHE | 970 | C |
| TYR | 1047 | A | TRP | 886 | B |
| LEU | 894 | A | ILE | 714 | C |
| PRO | 715 | A | LEU | 894 | B |
| GLN | 115 | A | ILE | 468 | C |
| LEU | 1001 | A | GLN | 1002 | C |
| TYR | 917 | A | VAL | 1129 | C |
| GLY | 1046 | A | GLY | 889 | B |
| LEU | 699 | A | VAL | 785 | B |
| MET | 900 | A | ALA | 1078 | C |
| LEU | 517 | A | ARG | 983 | B |
| PHE | 1089 | A | GLN | 913 | B |
| ALA | 570 | A | LEU | 966 | B |
| ASP | 796 | A | TYR | 707 | C |
| PHE | 464 | A | ASP | 198 | B |
| GLN | 563 | A | ASP | 40 | B |
| VAL | 1068 | A | ALA | 890 | B |
| ALA | 701 | A | LYS | 786 | B |
| ARG | 983 | A | SER | 383 | C |
| SER | 1123 | A | GLU | 1111 | B |
| ILE | 788 | A | ALA | 701 | C |
| VAL | 1040 | A | GLY | 1035 | B |
| TYR | 396 | A | TYR | 200 | B |
| LEU | 858 | A | PHE | 592 | C |
| ASP | 1041 | A | GLN | 784 | B |
| ASP | 796 | A | ASN | 709 | C |
| VAL | 976 | A | THR | 572 | C |
| ARG | 1039 | A | ARG | 1039 | C |
| TYR | 707 | A | ILE | 882 | B |
| TYR | 707 | A | GLN | 895 | B |
| HIS | 519 | A | VAL | 42 | B |
| ASN | 914 | A | PHE | 1089 | C |
| ARG | 319 | A | ASP | 737 | B |
| LEU | 984 | A | LYS | 386 | C |
| GLY | 891 | A | PRO | 1069 | C |

| | | | | | |
|---|---|---|---|---|---|
| SER | 884 | A | TYR | 707 | C |
| PRO | 1069 | A | ALA | 890 | B |
| VAL | 1129 | A | GLU | 918 | B |
| GLY | 566 | A | VAL | 42 | B |
| VAL | 705 | A | SER | 884 | B |
| ASP | 228 | A | LEU | 518 | C |
| ASP | 985 | A | THR | 415 | B |
| LYS | 1045 | A | LYS | 786 | B |
| GLY | 889 | A | VAL | 1040 | C |
| GLU | 988 | A | TYR | 380 | C |
| ALA | 570 | A | ASN | 856 | B |
| THR | 430 | A | ARG | 983 | B |
| GLN | 1002 | A | GLN | 1002 | C |
| TYR | 756 | A | GLN | 1002 | C |
| THR | 572 | A | VAL | 976 | B |
| LEU | 699 | A | ILE | 788 | B |
| ARG | 765 | A | THR | 961 | C |
| ILE | 788 | A | ASN | 703 | C |
| PRO | 863 | A | GLY | 667 | C |
| ASP | 228 | A | TYR | 396 | C |
| TRP | 353 | A | GLY | 232 | B |
| SER | 967 | A | ASP | 571 | C |
| ALA | 668 | A | LEU | 865 | B |
| THR | 1006 | A | GLN | 1005 | B |
| ILE | 1013 | A | THR | 1009 | B |
| ARG | 646 | A | PRO | 862 | B |
| SER | 708 | A | ASP | 796 | B |
| GLY | 971 | A | GLN | 755 | B |
| GLY | 232 | A | GLU | 465 | C |
| LEU | 861 | A | GLN | 613 | C |
| SER | 982 | A | LYS | 386 | C |
| PRO | 412 | A | VAL | 987 | C |
| LEU | 699 | A | GLN | 872 | B |
| ASP | 571 | A | SER | 967 | B |
| LEU | 984 | A | VAL | 382 | C |
| VAL | 1094 | A | TYR | 904 | B |
| LEU | 864 | A | GLY | 669 | C |
| LEU | 518 | A | LYS | 202 | B |
| THR | 859 | A | GLN | 613 | C |
| ASP | 568 | A | LYS | 854 | B |
| GLY | 232 | A | PHE | 464 | C |

| | | | | | |
|---|---|---|---|---|---|
| ILE | 712 | A | ILE | 896 | B |
| GLN | 895 | A | TYR | 707 | C |
| LEU | 865 | A | GLY | 669 | C |
| CYS | 379 | A | GLU | 988 | B |
| ARG | 466 | A | ILE | 233 | B |
| SER | 967 | A | ILE | 569 | C |
| GLY | 889 | A | LYS | 1045 | C |
| TYR | 200 | A | TYR | 396 | C |
| GLY | 1093 | A | TYR | 904 | B |
| THR | 1027 | A | ARG | 1039 | C |
| VAL | 705 | A | ALA | 893 | B |
| ALA | 570 | A | LYS | 964 | B |
| PHE | 970 | A | TYR | 756 | B |
| GLU | 224 | A | LEU | 560 | C |
| PRO | 897 | A | TYR | 707 | C |
| ARG | 1039 | A | GLU | 1031 | B |
| PHE | 562 | A | GLU | 224 | B |
| ILE | 231 | A | ARG | 466 | C |
| VAL | 382 | A | GLU | 988 | B |
| PRO | 862 | A | ILE | 666 | C |
| THR | 961 | A | ARG | 765 | B |
| GLU | 224 | A | PHE | 562 | C |
| ALA | 713 | A | ILE | 896 | B |
| LEU | 1145 | A | LEU | 1145 | C |
| PRO | 792 | A | VAL | 705 | C |
| GLN | 755 | A | ASN | 969 | C |
| GLY | 757 | A | GLN | 965 | C |
| VAL | 976 | A | ARG | 567 | C |
| GLN | 913 | A | ARG | 1091 | C |
| PHE | 855 | A | THR | 588 | C |
| SER | 1003 | A | PHE | 759 | B |
| THR | 1006 | A | PHE | 759 | B |
| PRO | 665 | A | LEU | 864 | B |
| GLY | 669 | A | LEU | 864 | B |
| PHE | 43 | A | LYS | 557 | C |
| GLY | 798 | A | TYR | 707 | C |
| PHE | 592 | A | LYS | 854 | B |
| SER | 975 | A | ALA | 570 | C |
| LYS | 113 | A | ILE | 468 | C |
| LEU | 894 | A | PRO | 715 | C |
| PHE | 43 | A | LYS | 558 | C |

| | | | | | |
|---|---|---|---|---|---|
| GLN | 787 | A | ASN | 703 | C |
| GLN | 1002 | A | GLN | 1002 | B |
| TYR | 789 | A | SER | 704 | C |
| ILE | 788 | A | LEU | 699 | C |
| THR | 866 | A | ALA | 668 | C |
| GLY | 1046 | A | ALA | 890 | B |
| GLY | 700 | A | TYR | 873 | B |
| ASP | 994 | A | PHE | 970 | C |
| VAL | 1129 | A | TYR | 917 | B |
| TRP | 886 | A | ASN | 1108 | C |
| PRO | 863 | A | ALA | 668 | C |
| ALA | 372 | A | GLU | 406 | C |
| TYR | 38 | A | PHE | 562 | C |
| PHE | 565 | A | VAL | 42 | B |
| ALA | 520 | A | LYS | 41 | B |
| GLN | 762 | A | THR | 961 | C |
| VAL | 705 | A | LYS | 790 | B |
| GLY | 667 | A | PRO | 862 | B |
| GLN | 1036 | A | LYS | 1038 | C |
| VAL | 987 | A | ASP | 427 | B |
| THR | 1077 | A | MET | 900 | B |
| ASP | 745 | A | ARG | 319 | C |
| SER | 1147 | A | LEU | 1145 | C |
| GLN | 965 | A | TYR | 756 | B |
| PHE | 797 | A | TYR | 707 | C |
| GLY | 413 | A | VAL | 987 | C |
| VAL | 705 | A | PRO | 792 | B |
| THR | 284 | A | LEU | 560 | C |
| VAL | 963 | A | ILE | 569 | C |
| ILE | 896 | A | SER | 708 | C |
| GLU | 918 | A | SER | 1123 | C |
| ALA | 879 | A | VAL | 705 | C |
| ARG | 567 | A | ARG | 44 | B |
| PRO | 862 | A | ARG | 646 | C |
| THR | 547 | A | SER | 982 | B |
| LYS | 558 | A | PHE | 43 | B |
| GLN | 965 | A | GLN | 762 | B |
| ALA | 570 | A | VAL | 963 | B |
| SER | 967 | A | ALA | 570 | C |
| ILE | 1013 | A | LEU | 1012 | B |
| ALA | 706 | A | GLN | 895 | B |

| | | | | | |
|---|---|---|---|---|---|
| TYR | 904 | A | VAL | 1094 | C |
| HIS | 519 | A | LYS | 41 | B |
| ILE | 1130 | A | TYR | 917 | B |
| LEU | 754 | A | GLN | 52 | C |
| GLN | 414 | A | TYR | 369 | B |
| LEU | 699 | A | LYS | 786 | B |
| PHE | 464 | A | ILE | 233 | B |
| PRO | 897 | A | ASN | 710 | C |
| TYR | 707 | A | ALA | 899 | B |
| GLN | 762 | A | THR | 1006 | C |
| GLY | 199 | A | PHE | 464 | C |
| ASP | 568 | A | ASN | 856 | B |
| ASN | 282 | A | LYS | 558 | C |
| LYS | 964 | A | ALA | 570 | C |
| LYS | 41 | A | GLN | 563 | C |
| TYR | 369 | A | GLY | 416 | C |
| ALA | 570 | A | VAL | 976 | B |
| LEU | 1141 | A | LEU | 1145 | B |
| ALA | 713 | A | GLN | 895 | B |
| VAL | 976 | A | ALA | 570 | C |
| GLU | 988 | A | SER | 383 | C |
| GLN | 895 | A | ALA | 706 | C |
| ARG | 466 | A | ILE | 231 | B |
| GLY | 1035 | A | VAL | 1040 | C |
| ASP | 737 | A | PHE | 592 | C |
| GLY | 545 | A | SER | 982 | B |
| ILE | 788 | A | SER | 704 | C |
| SER | 968 | A | TYR | 756 | B |
| TYR | 904 | A | ILE | 712 | C |
| PHE | 374 | A | ARG | 408 | C |
| PRO | 1090 | A | GLN | 913 | B |
| PHE | 43 | A | GLN | 563 | C |
| SER | 758 | A | THR | 961 | C |
| TYR | 917 | A | VAL | 1128 | C |
| ASP | 571 | A | SER | 968 | B |
| GLN | 913 | A | PHE | 1089 | C |
| SER | 704 | A | TYR | 789 | B |
| PHE | 592 | A | ASN | 856 | B |
| ASN | 978 | A | GLY | 548 | C |
| LEU | 1145 | A | LEU | 1141 | B |
| LYS | 790 | A | VAL | 705 | C |

| | | | | | |
|---|---|---|---|---|---|
| ASN | 960 | A | ILE | 569 | C |
| PHE | 464 | A | GLY | 199 | B |
| GLY | 1093 | A | GLN | 913 | B |
| TYR | 707 | A | THR | 883 | B |
| GLN | 1002 | A | GLN | 1005 | B |
| ALA | 1078 | A | MET | 900 | B |
| LEU | 864 | A | GLY | 667 | C |
| SER | 982 | A | ASP | 389 | C |
| LYS | 1045 | A | GLY | 889 | B |
| TYR | 917 | A | ALA | 1080 | C |
| MET | 697 | A | TYR | 873 | B |
| THR | 547 | A | LEU | 981 | B |
| GLN | 895 | A | SER | 711 | C |
| SER | 708 | A | GLN | 895 | B |
| SER | 704 | A | LYS | 790 | B |
| ASN | 856 | A | PHE | 592 | C |
| ALA | 893 | A | VAL | 705 | C |
| LEU | 981 | A | THR | 547 | C |
| TYR | 756 | A | SER | 968 | C |
| ARG | 1039 | A | THR | 1027 | B |
| MET | 900 | A | PRO | 1079 | C |
| SER | 704 | A | ILE | 788 | B |
| GLN | 1002 | A | PHE | 759 | B |
| LEU | 1012 | A | GLN | 1010 | C |
| GLY | 891 | A | VAL | 1068 | C |
| ARG | 44 | A | GLY | 566 | C |
| ASP | 737 | A | ARG | 319 | C |
| VAL | 705 | A | GLY | 880 | B |
| PHE | 562 | A | PRO | 225 | B |
| LEU | 699 | A | MET | 869 | B |
| VAL | 1094 | A | MET | 900 | B |
| TYR | 707 | A | GLY | 798 | B |
| VAL | 42 | A | ARG | 567 | C |
| LYS | 41 | A | ALA | 520 | C |
| LYS | 41 | A | PHE | 565 | C |
| SER | 708 | A | ILE | 896 | B |
| ILE | 712 | A | MET | 900 | B |
| ASP | 985 | A | THR | 385 | C |
| PHE | 592 | A | GLY | 857 | B |
| GLN | 115 | A | ARG | 466 | C |
| GLY | 669 | A | LEU | 865 | B |

| | | | | | |
|---|---|---|---|---|---|
| ILE | 1130 | A | GLY | 798 | B |
| SER | 383 | A | LEU | 984 | B |
| GLN | 787 | A | GLY | 700 | C |
| GLY | 857 | A | PHE | 592 | C |
| PRO | 862 | A | ALA | 668 | C |
| LEU | 864 | A | CYS | 671 | C |
| LEU | 560 | A | TYR | 38 | B |
| PHE | 464 | A | GLY | 232 | B |
| SER | 708 | A | PRO | 897 | B |
| ILE | 712 | A | TYR | 904 | B |
| TYR | 380 | A | GLU | 988 | B |
| GLN | 920 | A | ILE | 1130 | C |
| ASN | 978 | A | THR | 547 | C |
| PRO | 897 | A | THR | 1077 | C |
| ILE | 896 | A | ALA | 713 | C |
| PHE | 1089 | A | GLU | 918 | B |
| ARG | 567 | A | VAL | 42 | B |
| GLN | 563 | A | VAL | 42 | B |
| GLY | 232 | A | TRP | 353 | C |
| GLU | 1017 | A | ARG | 1019 | B |
| LEU | 1034 | A | ASP | 1041 | C |
| ASP | 428 | A | ASP | 198 | B |
| MET | 869 | A | THR | 696 | C |
| PHE | 43 | A | LEU | 560 | C |
| THR | 588 | A | PHE | 855 | B |
| TYR | 1007 | A | GLN | 762 | B |
| GLY | 1124 | A | GLU | 918 | B |
| PRO | 1079 | A | GLN | 913 | B |
| LEU | 1141 | A | LEU | 1141 | B |
| PHE | 1121 | A | THR | 912 | B |
| ILE | 468 | A | GLU | 132 | B |
| LEU | 864 | A | PRO | 665 | C |
| LYS | 386 | A | LEU | 981 | B |
| ALA | 668 | A | PRO | 863 | B |
| ASN | 978 | A | GLY | 545 | C |
| ILE | 712 | A | GLN | 895 | B |
| GLN | 563 | A | TYR | 38 | B |
| LEU | 894 | A | VAL | 705 | C |
| PRO | 384 | A | GLU | 988 | B |
| GLU | 1144 | A | LEU | 1145 | C |
| GLN | 563 | A | LYS | 41 | B |

| | | | | | |
|---|---|---|---|---|---|
| ARG | 983 | A | THR | 430 | C |
| VAL | 1040 | A | GLN | 1036 | B |
| LEU | 894 | A | PRO | 1069 | C |
| ALA | 890 | A | GLY | 1046 | C |
| GLY | 700 | A | GLN | 787 | B |
| SER | 982 | A | LEU | 390 | C |
| PRO | 426 | A | ASP | 198 | B |
| VAL | 47 | A | ILE | 569 | C |
| TYR | 200 | A | GLU | 516 | C |
| LEU | 1141 | A | LEU | 1145 | C |
| LEU | 699 | A | TYR | 873 | B |
| SER | 1123 | A | GLU | 918 | B |
| MET | 697 | A | GLN | 779 | B |
| LYS | 964 | A | ILE | 569 | C |
| ASP | 979 | A | LEU | 546 | C |
| ARG | 44 | A | ARG | 567 | C |
| LYS | 921 | A | ILE | 1130 | C |
| VAL | 1040 | A | LEU | 1034 | B |

**Table S2.** The list of the inter-protomer contacts of the protomer A in the structure of SARS-CoV-2 S-GSAS/D614 in the open state (pdb id 7KDH).

| S protein residue name | S protein residue number | S protein chain | S protein residue name | S protein residue number | S protein chain |
|---|---|---|---|---|---|
| GLN | 563 | A | ARG | 44 | B |
| ARG | 319 | A | MET | 740 | B |
| ALA | 570 | A | LEU | 966 | B |
| ARG | 44 | A | ASP | 571 | C |
| GLY | 381 | A | ARG | 983 | B |
| TYR | 873 | A | MET | 697 | C |
| ASP | 737 | A | ASN | 317 | C |
| ILE | 1130 | A | LYS | 921 | B |
| TYR | 200 | A | TYR | 396 | C |
| LEU | 984 | A | GLY | 381 | C |
| LEU | 699 | A | LEU | 865 | B |
| ASN | 703 | A | TYR | 789 | B |
| ARG | 1091 | A | ASP | 1118 | B |
| ASP | 40 | A | PHE | 562 | C |
| GLN | 784 | A | LYS | 1045 | C |
| LEU | 1034 | A | ASP | 1041 | C |
| VAL | 1094 | A | ILE | 896 | B |
| ILE | 1013 | A | LEU | 1012 | B |
| GLU | 1072 | A | ALA | 892 | B |
| GLN | 755 | A | PHE | 970 | C |
| LYS | 386 | A | LEU | 984 | B |
| SER | 967 | A | ILE | 569 | C |
| PHE | 43 | A | PHE | 559 | C |
| ARG | 995 | A | ASP | 994 | B |
| VAL | 976 | A | ARG | 567 | C |
| PHE | 1089 | A | GLN | 913 | B |
| TYR | 873 | A | LEU | 699 | C |
| GLN | 787 | A | GLY | 700 | C |
| TYR | 707 | A | PHE | 898 | B |
| LYS | 41 | A | PRO | 521 | C |
| GLY | 889 | A | VAL | 1040 | C |
| TYR | 756 | A | PHE | 970 | C |
| LEU | 894 | A | ALA | 713 | C |
| ASN | 978 | A | GLY | 545 | C |
| ARG | 1039 | A | THR | 1027 | B |

| | | | | | |
|---|---|---|---|---|---|
| PHE | 1121 | A | THR | 912 | B |
| THR | 1006 | A | GLN | 762 | B |
| PRO | 230 | A | ARG | 355 | C |
| ALA | 713 | A | LEU | 894 | B |
| ALA | 879 | A | VAL | 705 | C |
| LEU | 699 | A | GLN | 787 | B |
| GLN | 613 | A | THR | 859 | B |
| ASP | 745 | A | ARG | 319 | C |
| ALA | 701 | A | ILE | 788 | B |
| ASN | 703 | A | LYS | 790 | B |
| TYR | 756 | A | ASN | 969 | C |
| TYR | 38 | A | PHE | 562 | C |
| ILE | 714 | A | LEU | 894 | B |
| ASN | 165 | A | ILE | 468 | C |
| THR | 284 | A | LEU | 560 | C |
| ILE | 788 | A | ALA | 701 | C |
| VAL | 1068 | A | GLY | 891 | B |
| ASN | 907 | A | ARG | 1107 | C |
| MET | 900 | A | ILE | 712 | C |
| ASP | 979 | A | LEU | 546 | C |
| ILE | 896 | A | SER | 711 | C |
| PHE | 43 | A | GLY | 566 | C |
| GLY | 798 | A | ILE | 1130 | C |
| TYR | 904 | A | ARG | 1107 | C |
| PRO | 1069 | A | ALA | 892 | B |
| LEU | 1034 | A | LYS | 1045 | C |
| ASN | 394 | A | ASP | 228 | B |
| GLY | 1093 | A | GLN | 913 | B |
| GLN | 965 | A | GLY | 757 | B |
| PRO | 1069 | A | LEU | 894 | B |
| PHE | 888 | A | LYS | 1045 | C |
| ALA | 570 | A | ASN | 856 | B |
| TYR | 707 | A | ALA | 899 | B |
| LEU | 1141 | A | LEU | 1141 | C |
| THR | 866 | A | ALA | 668 | C |
| GLN | 784 | A | ASP | 1041 | C |
| VAL | 705 | A | TYR | 789 | B |
| VAL | 705 | A | ALA | 879 | B |
| GLY | 757 | A | PHE | 970 | C |
| SER | 1003 | A | PHE | 759 | B |
| LEU | 546 | A | ASP | 979 | B |

| | | | | | |
|---|---|---|---|---|---|
| LEU | 1141 | A | LEU | 1145 | C |
| THR | 588 | A | PHE | 855 | B |
| LEU | 894 | A | PRO | 715 | C |
| ALA | 570 | A | VAL | 963 | B |
| LEU | 865 | A | MET | 697 | C |
| THR | 1077 | A | MET | 900 | B |
| ILE | 1013 | A | ALA | 1016 | B |
| GLN | 895 | A | ILE | 712 | C |
| LYS | 1045 | A | GLN | 784 | B |
| LEU | 699 | A | TYR | 873 | B |
| GLY | 700 | A | GLN | 787 | B |
| LEU | 864 | A | ILE | 666 | C |
| PRO | 1079 | A | MET | 900 | B |
| GLU | 1031 | A | ARG | 1039 | C |
| ALA | 701 | A | TYR | 789 | B |
| GLU | 702 | A | GLN | 872 | B |
| ASP | 994 | A | PHE | 970 | C |
| PRO | 1069 | A | GLY | 891 | B |
| ALA | 701 | A | GLN | 787 | B |
| GLU | 918 | A | VAL | 1128 | C |
| GLN | 965 | A | PHE | 759 | B |
| GLY | 669 | A | LEU | 865 | B |
| LEU | 699 | A | ILE | 788 | B |
| LEU | 864 | A | ILE | 670 | C |
| SER | 968 | A | GLN | 755 | B |
| GLY | 667 | A | PRO | 862 | B |
| VAL | 976 | A | ALA | 570 | C |
| GLY | 757 | A | SER | 968 | C |
| SER | 383 | A | GLU | 988 | B |
| GLY | 798 | A | TYR | 707 | C |
| ILE | 896 | A | SER | 708 | C |
| ASP | 737 | A | PHE | 592 | C |
| LYS | 786 | A | GLY | 700 | C |
| SER | 708 | A | ILE | 896 | B |
| CYS | 379 | A | GLU | 988 | B |
| ASP | 571 | A | SER | 967 | B |
| GLU | 702 | A | LYS | 790 | B |
| GLY | 971 | A | ASP | 994 | B |
| GLN | 913 | A | GLY | 1093 | C |
| PHE | 1089 | A | GLU | 918 | B |
| THR | 961 | A | SER | 758 | B |

| | | | | | |
|---|---|---|---|---|---|
| PRO | 862 | A | GLY | 667 | C |
| SER | 968 | A | GLY | 757 | B |
| TYR | 789 | A | GLU | 702 | C |
| ALA | 647 | A | PRO | 862 | B |
| LEU | 517 | A | ARG | 983 | B |
| ARG | 44 | A | ARG | 567 | C |
| GLY | 232 | A | PHE | 464 | C |
| LEU | 865 | A | GLY | 669 | C |
| THR | 415 | A | ASP | 985 | C |
| PHE | 43 | A | LEU | 560 | C |
| THR | 1009 | A | THR | 1009 | B |
| ALA | 890 | A | LYS | 1045 | C |
| ALA | 520 | A | LYS | 41 | B |
| PRO | 897 | A | SER | 711 | C |
| GLN | 965 | A | TYR | 756 | B |
| GLY | 566 | A | PHE | 43 | B |
| ALA | 892 | A | PRO | 1069 | C |
| THR | 167 | A | ARG | 466 | C |
| LYS | 386 | A | ARG | 983 | B |
| GLU | 1017 | A | ARG | 1019 | B |
| GLN | 314 | A | SER | 735 | B |
| LEU | 894 | A | VAL | 705 | C |
| GLN | 787 | A | ALA | 701 | C |
| LEU | 1145 | A | LEU | 1145 | C |
| PHE | 1042 | A | GLU | 1031 | B |
| LEU | 546 | A | VAL | 976 | B |
| GLY | 971 | A | TYR | 756 | B |
| LEU | 1012 | A | ILE | 1013 | C |
| ASN | 978 | A | GLY | 548 | C |
| THR | 1027 | A | ARG | 1039 | C |
| VAL | 705 | A | THR | 791 | B |
| ASP | 796 | A | ASN | 709 | C |
| TYR | 904 | A | ILE | 712 | C |
| THR | 912 | A | PHE | 1121 | C |
| LEU | 1145 | A | SER | 1147 | B |
| GLN | 762 | A | GLN | 965 | C |
| LEU | 858 | A | PHE | 592 | C |
| THR | 883 | A | TYR | 707 | C |
| GLY | 667 | A | LEU | 864 | B |
| GLY | 667 | A | PRO | 863 | B |
| SER | 711 | A | PRO | 897 | B |

| | | | | | |
|---|---|---|---|---|---|
| VAL | 1129 | A | TYR | 917 | B |
| SER | 982 | A | THR | 547 | C |
| ARG | 44 | A | GLY | 566 | C |
| LEU | 560 | A | THR | 284 | B |
| THR | 572 | A | ASN | 856 | B |
| GLU | 1072 | A | LEU | 894 | B |
| ASN | 856 | A | ALA | 570 | C |
| THR | 430 | A | ARG | 983 | B |
| PRO | 862 | A | ALA | 647 | C |
| PRO | 1079 | A | GLN | 913 | B |
| ARG | 567 | A | PHE | 43 | B |
| THR | 1006 | A | PHE | 759 | B |
| ASN | 317 | A | GLY | 857 | B |
| ARG | 1091 | A | ASN | 907 | B |
| LYS | 1045 | A | ALA | 890 | B |
| THR | 385 | A | ASP | 985 | B |
| TYR | 707 | A | PRO | 897 | B |
| SER | 884 | A | VAL | 705 | C |
| PRO | 897 | A | THR | 1077 | C |
| LEU | 1001 | A | GLN | 1002 | C |
| ASP | 574 | A | PHE | 855 | B |
| ASP | 994 | A | GLY | 971 | C |
| LEU | 894 | A | PRO | 1069 | C |
| ASP | 1041 | A | LEU | 1034 | B |
| LYS | 558 | A | GLY | 283 | B |
| GLN | 787 | A | GLU | 702 | C |
| THR | 961 | A | GLN | 762 | B |
| ALA | 713 | A | GLN | 895 | B |
| PRO | 897 | A | ASN | 710 | C |
| ASN | 856 | A | ASP | 568 | C |
| VAL | 1094 | A | TYR | 904 | B |
| PRO | 230 | A | ARG | 466 | C |
| LEU | 1145 | A | GLU | 1144 | B |
| GLY | 381 | A | LEU | 984 | B |
| ARG | 983 | A | VAL | 382 | C |
| ASP | 1041 | A | SER | 1030 | B |
| LYS | 386 | A | ASP | 985 | B |
| PHE | 855 | A | PHE | 592 | C |
| ILE | 670 | A | LEU | 864 | B |
| ALA | 668 | A | MET | 869 | B |
| ASP | 571 | A | VAL | 976 | B |

| | | | | | |
|---|---|---|---|---|---|
| GLY | 413 | A | LYS | 986 | C |
| PHE | 456 | A | SER | 383 | B |
| ALA | 1078 | A | MET | 900 | B |
| TYR | 38 | A | GLN | 563 | C |
| GLN | 895 | A | SER | 711 | C |
| LYS | 854 | A | ASP | 568 | C |
| ARG | 1039 | A | GLU | 1031 | B |
| VAL | 42 | A | PHE | 562 | C |
| SER | 383 | A | ARG | 983 | B |
| THR | 739 | A | ASN | 317 | C |
| ASP | 198 | A | PRO | 463 | C |
| THR | 887 | A | TYR | 1047 | C |
| VAL | 860 | A | ARG | 646 | C |
| PHE | 970 | A | GLN | 755 | B |
| TYR | 1047 | A | THR | 887 | B |
| GLN | 563 | A | ASP | 40 | B |
| ALA | 1070 | A | ALA | 892 | B |
| ILE | 712 | A | GLN | 895 | B |
| THR | 961 | A | ARG | 765 | B |
| ILE | 1130 | A | GLY | 798 | B |
| TYR | 707 | A | PHE | 797 | B |
| GLN | 913 | A | PHE | 1121 | C |
| SER | 383 | A | ASP | 985 | B |
| ASP | 737 | A | ARG | 319 | C |
| PRO | 862 | A | ARG | 646 | C |
| GLY | 857 | A | PHE | 592 | C |
| PHE | 759 | A | GLY | 999 | C |
| ASN | 709 | A | ASP | 796 | B |
| GLN | 895 | A | ALA | 713 | C |
| THR | 385 | A | THR | 415 | C |
| PRO | 1069 | A | ALA | 890 | B |
| TYR | 756 | A | ARG | 995 | C |
| SER | 758 | A | LYS | 964 | C |
| GLY | 413 | A | VAL | 987 | C |
| PHE | 759 | A | PHE | 970 | C |
| LYS | 1045 | A | GLY | 889 | B |
| PRO | 792 | A | TYR | 707 | C |
| TYR | 904 | A | VAL | 1094 | C |
| LYS | 41 | A | ALA | 520 | C |
| LYS | 964 | A | ILE | 569 | C |
| ALA | 899 | A | TYR | 707 | C |

| | | | | | |
|---|---|---|---|---|---|
| GLY | 700 | A | ILE | 788 | B |
| GLY | 891 | A | LYS | 1045 | C |
| GLN | 563 | A | GLY | 283 | B |
| THR | 1006 | A | GLN | 1005 | B |
| GLN | 1010 | A | ALA | 766 | B |
| SER | 1123 | A | ASN | 914 | B |
| ASP | 985 | A | PRO | 384 | C |
| ILE | 973 | A | GLY | 381 | C |
| LEU | 864 | A | PRO | 665 | C |
| GLU | 918 | A | SER | 1123 | C |
| ASN | 856 | A | THR | 572 | C |
| GLY | 669 | A | LEU | 864 | B |
| TYR | 707 | A | ILE | 794 | B |
| ALA | 570 | A | SER | 967 | B |
| TYR | 707 | A | ASP | 796 | B |
| ALA | 575 | A | PHE | 43 | B |
| SER | 45 | A | ARG | 567 | C |
| ASN | 234 | A | ARG | 466 | C |
| ASN | 394 | A | PRO | 230 | B |
| GLY | 232 | A | ARG | 466 | C |
| LYS | 557 | A | SER | 45 | B |
| LEU | 1034 | A | VAL | 1040 | C |
| SER | 704 | A | THR | 791 | B |
| THR | 572 | A | VAL | 976 | B |
| TYR | 707 | A | ILE | 896 | B |
| GLU | 918 | A | GLY | 1124 | C |
| PHE | 1121 | A | ASN | 914 | B |
| GLY | 548 | A | ASN | 978 | B |
| ARG | 1091 | A | GLN | 913 | B |
| GLN | 895 | A | TYR | 707 | C |
| GLN | 564 | A | LYS | 41 | B |
| TYR | 904 | A | GLY | 1093 | C |
| TYR | 917 | A | ILE | 1130 | C |
| THR | 883 | A | VAL | 705 | C |
| PHE | 592 | A | GLY | 857 | B |
| ILE | 569 | A | VAL | 47 | B |
| ASP | 571 | A | SER | 975 | B |
| TYR | 1047 | A | TRP | 886 | B |
| GLU | 988 | A | SER | 383 | C |
| ILE | 1130 | A | TYR | 917 | B |
| LYS | 41 | A | GLN | 564 | C |

| | | | | | |
|---|---|---|---|---|---|
| MET | 869 | A | GLY | 669 | C |
| PHE | 565 | A | VAL | 42 | B |
| VAL | 705 | A | GLN | 895 | B |
| GLN | 762 | A | THR | 961 | C |
| PHE | 562 | A | LYS | 41 | B |
| PRO | 589 | A | MET | 740 | B |
| ASP | 228 | A | TYR | 396 | C |
| GLN | 762 | A | SER | 1003 | C |
| LEU | 981 | A | LYS | 386 | C |
| PRO | 1140 | A | LEU | 1141 | C |
| LEU | 984 | A | VAL | 382 | C |
| LEU | 1145 | A | LEU | 1145 | B |
| THR | 1009 | A | ILE | 1013 | C |
| TYR | 756 | A | GLN | 965 | C |
| LEU | 894 | A | GLU | 1072 | C |
| SER | 982 | A | GLY | 545 | C |
| PRO | 384 | A | GLU | 988 | B |
| PHE | 759 | A | GLN | 1002 | C |
| GLN | 762 | A | GLN | 1010 | C |
| LYS | 41 | A | PHE | 562 | C |
| ALA | 570 | A | VAL | 976 | B |
| ILE | 712 | A | MET | 900 | B |
| MET | 740 | A | ARG | 319 | C |
| ILE | 666 | A | PRO | 862 | B |
| PRO | 665 | A | LEU | 864 | B |
| GLN | 563 | A | TYR | 279 | B |
| GLY | 757 | A | GLN | 965 | C |
| GLY | 891 | A | VAL | 1068 | C |
| ASP | 427 | A | GLU | 990 | C |
| GLU | 516 | A | ASP | 228 | B |
| VAL | 42 | A | GLY | 566 | C |
| SER | 1030 | A | PHE | 1042 | C |
| ARG | 983 | A | SER | 383 | C |
| GLN | 564 | A | VAL | 42 | B |
| GLN | 563 | A | VAL | 42 | B |
| VAL | 1068 | A | ALA | 890 | B |
| ALA | 1080 | A | TYR | 917 | B |
| THR | 547 | A | ASN | 978 | B |
| GLY | 889 | A | ASP | 1041 | C |
| LYS | 557 | A | PHE | 43 | B |
| TYR | 369 | A | GLN | 414 | C |

| | | | | | |
|---|---|---|---|---|---|
| ASP | 979 | A | THR | 547 | C |
| TYR | 707 | A | THR | 883 | B |
| GLU | 918 | A | PHE | 1089 | C |
| TYR | 756 | A | GLY | 971 | C |
| ARG | 1019 | A | GLU | 1017 | C |
| LYS | 790 | A | GLU | 702 | C |
| GLY | 700 | A | LYS | 786 | B |
| CYS | 671 | A | LEU | 864 | B |
| LYS | 786 | A | ALA | 701 | C |
| TYR | 1007 | A | GLN | 762 | B |
| GLN | 314 | A | THR | 768 | B |
| ILE | 712 | A | ILE | 896 | B |
| LEU | 560 | A | PHE | 43 | B |
| SER | 1123 | A | GLU | 1111 | B |
| VAL | 705 | A | PRO | 792 | B |
| LYS | 790 | A | SER | 704 | C |
| TYR | 707 | A | GLY | 798 | B |
| ALA | 706 | A | GLN | 895 | B |
| MET | 869 | A | MET | 697 | C |
| ARG | 44 | A | GLN | 563 | C |
| ASN | 969 | A | GLN | 755 | B |
| GLY | 199 | A | PHE | 464 | C |
| ARG | 765 | A | THR | 961 | C |
| GLU | 224 | A | PHE | 562 | C |
| GLU | 1031 | A | PHE | 1042 | C |
| PRO | 863 | A | GLY | 669 | C |
| PHE | 43 | A | LYS | 558 | C |
| ASP | 796 | A | SER | 708 | C |
| GLU | 773 | A | GLU | 1017 | C |
| ALA | 713 | A | ILE | 896 | B |
| ARG | 1107 | A | GLN | 913 | B |
| GLY | 999 | A | PHE | 759 | B |
| ALA | 701 | A | LYS | 786 | B |
| ASN | 317 | A | MET | 740 | B |
| GLN | 1010 | A | LEU | 1012 | B |
| PHE | 855 | A | ASP | 568 | C |
| ASP | 1041 | A | GLY | 889 | B |
| GLN | 115 | A | ARG | 466 | C |
| ASN | 710 | A | PRO | 897 | B |
| GLN | 913 | A | PRO | 1090 | C |
| GLY | 1124 | A | GLU | 918 | B |

| | | | | | |
|---|---|---|---|---|---|
| VAL | 785 | A | LEU | 699 | C |
| ASP | 796 | A | TYR | 707 | C |
| TYR | 789 | A | SER | 704 | C |
| GLN | 957 | A | ARG | 765 | B |
| ALA | 972 | A | GLN | 755 | B |
| LEU | 699 | A | LYS | 786 | B |
| GLN | 1036 | A | LYS | 1038 | C |
| TYR | 369 | A | ARG | 408 | C |
| ALA | 890 | A | VAL | 1040 | C |
| TYR | 200 | A | ARG | 355 | C |
| ASP | 1041 | A | GLN | 784 | B |
| PHE | 1089 | A | TYR | 917 | B |
| ASP | 994 | A | ARG | 995 | C |
| GLN | 895 | A | SER | 708 | C |
| PHE | 565 | A | PHE | 43 | B |
| LEU | 966 | A | ALA | 570 | C |
| GLN | 913 | A | PHE | 1089 | C |
| TYR | 789 | A | VAL | 705 | C |
| LEU | 864 | A | ALA | 668 | C |
| ILE | 794 | A | TYR | 707 | C |
| GLU | 1111 | A | SER | 1123 | C |
| ILE | 870 | A | LEU | 699 | C |
| PRO | 897 | A | SER | 708 | C |
| SER | 704 | A | TYR | 789 | B |
| GLN | 755 | A | ASN | 969 | C |
| GLN | 563 | A | TYR | 38 | B |
| PHE | 898 | A | TYR | 707 | C |
| ARG | 1039 | A | ARG | 1039 | C |
| ALA | 890 | A | GLY | 1046 | C |
| VAL | 976 | A | THR | 572 | C |
| ARG | 983 | A | GLY | 381 | C |
| LYS | 790 | A | VAL | 705 | C |
| LYS | 41 | A | GLN | 563 | C |
| ARG | 567 | A | ARG | 44 | B |
| LEU | 560 | A | TYR | 38 | B |
| GLN | 913 | A | ARG | 1091 | C |
| PRO | 412 | A | VAL | 987 | C |
| SER | 45 | A | LYS | 557 | C |
| GLY | 1046 | A | ALA | 890 | B |
| VAL | 1122 | A | ASN | 914 | B |
| LYS | 964 | A | SER | 758 | B |

| | | | | | |
|---|---|---|---|---|---|
| TYR | 369 | A | THR | 415 | C |
| ILE | 712 | A | PRO | 897 | B |
| LEU | 560 | A | GLY | 283 | B |
| MET | 697 | A | MET | 869 | B |
| LEU | 984 | A | TYR | 380 | C |
| THR | 547 | A | SER | 982 | B |
| GLN | 755 | A | SER | 968 | C |
| ASP | 1118 | A | ARG | 1091 | C |
| LEU | 984 | A | THR | 385 | C |
| GLU | 1092 | A | GLN | 913 | B |
| VAL | 382 | A | GLU | 988 | B |
| VAL | 1040 | A | SER | 1030 | B |
| TYR | 756 | A | SER | 968 | C |
| PHE | 855 | A | PRO | 589 | C |
| LEU | 865 | A | ALA | 668 | C |
| LEU | 864 | A | GLY | 667 | C |
| GLN | 1002 | A | THR | 998 | B |
| SER | 982 | A | LEU | 390 | C |
| TRP | 886 | A | ASN | 1108 | C |
| ASP | 40 | A | HIS | 519 | C |
| GLY | 1046 | A | GLY | 891 | B |
| THR | 167 | A | ILE | 468 | C |
| VAL | 42 | A | GLN | 563 | C |
| THR | 1009 | A | THR | 1009 | C |
| GLY | 413 | A | ASP | 985 | C |
| SER | 1030 | A | VAL | 1040 | C |
| ASP | 198 | A | PHE | 464 | C |
| TYR | 1047 | A | ALA | 890 | B |
| ALA | 668 | A | PRO | 863 | B |
| TYR | 707 | A | PRO | 792 | B |
| ALA | 668 | A | THR | 866 | B |
| GLU | 516 | A | TYR | 200 | B |
| ASN | 1108 | A | TRP | 886 | B |
| VAL | 1094 | A | MET | 900 | B |
| PRO | 897 | A | ILE | 712 | C |
| GLN | 755 | A | GLY | 971 | C |
| PHE | 43 | A | LYS | 557 | C |
| GLN | 314 | A | ASN | 764 | B |
| GLU | 1017 | A | GLU | 773 | B |
| TYR | 200 | A | PHE | 464 | C |
| MET | 900 | A | ALA | 1078 | C |

| | | | | | |
|---|---|---|---|---|---|
| ALA | 372 | A | ARG | 403 | C |
| SER | 968 | A | SER | 758 | B |
| GLY | 891 | A | PRO | 1069 | C |
| MET | 697 | A | TYR | 873 | B |
| TRP | 886 | A | TYR | 1047 | C |
| VAL | 42 | A | PHE | 565 | C |
| PRO | 230 | A | TYR | 396 | C |
| GLY | 669 | A | THR | 866 | B |
| TYR | 917 | A | VAL | 1128 | C |
| ALA | 668 | A | LEU | 864 | B |
| ARG | 1107 | A | TRP | 886 | B |
| ASN | 914 | A | SER | 1123 | C |
| PHE | 592 | A | PHE | 855 | B |
| ASP | 427 | A | VAL | 987 | C |
| PRO | 1090 | A | GLN | 913 | B |
| MET | 869 | A | ALA | 668 | C |
| THR | 547 | A | ASP | 979 | B |
| GLU | 1144 | A | LEU | 1141 | C |
| ALA | 893 | A | VAL | 705 | C |
| MET | 697 | A | LEU | 865 | B |
| TYR | 789 | A | ALA | 701 | C |
| PHE | 970 | A | TYR | 756 | B |
| GLY | 566 | A | VAL | 42 | B |
| SER | 982 | A | LEU | 546 | C |
| GLY | 889 | A | GLY | 1046 | C |
| SER | 708 | A | GLN | 895 | B |
| ASN | 282 | A | LEU | 560 | C |
| SER | 968 | A | TYR | 756 | B |
| TYR | 707 | A | GLN | 895 | B |
| ASP | 985 | A | THR | 385 | C |
| TYR | 707 | A | ILE | 882 | B |
| VAL | 1068 | A | ALA | 892 | B |
| SER | 967 | A | ASP | 571 | C |
| CYS | 662 | A | LEU | 864 | B |
| MET | 900 | A | THR | 1077 | C |
| SER | 982 | A | LYS | 386 | C |
| ILE | 231 | A | ARG | 466 | C |
| GLN | 787 | A | ASN | 703 | C |
| VAL | 705 | A | THR | 883 | B |
| ARG | 983 | A | LEU | 390 | C |
| GLY | 283 | A | LEU | 560 | C |

| | | | | | |
|---|---|---|---|---|---|
| PRO | 521 | A | LYS | 41 | B |
| PRO | 897 | A | TYR | 707 | C |
| THR | 859 | A | PHE | 592 | C |
| ILE | 1130 | A | GLN | 920 | B |
| GLU | 1072 | A | ALA | 893 | B |
| HIS | 519 | A | VAL | 42 | B |
| ARG | 567 | A | VAL | 976 | B |
| GLY | 232 | A | GLU | 465 | C |
| ARG | 646 | A | THR | 866 | B |
| ASN | 978 | A | THR | 547 | C |
| GLY | 1093 | A | TYR | 904 | B |
| LYS | 776 | A | LYS | 947 | C |
| GLY | 1046 | A | GLY | 889 | B |
| ARG | 1107 | A | TYR | 904 | B |
| SER | 708 | A | PRO | 897 | B |
| ASN | 703 | A | ILE | 788 | B |
| SER | 758 | A | THR | 961 | C |
| GLN | 1005 | A | GLN | 1002 | C |
| VAL | 963 | A | ALA | 570 | C |
| LEU | 699 | A | VAL | 785 | B |
| ARG | 765 | A | GLN | 957 | C |
| TYR | 369 | A | GLY | 416 | C |
| ILE | 788 | A | ASN | 703 | C |
| LYS | 790 | A | ASN | 703 | C |
| LYS | 386 | A | LEU | 981 | B |
| TYR | 396 | A | PRO | 230 | B |
| MET | 900 | A | PRO | 1079 | C |
| VAL | 705 | A | LYS | 790 | B |
| GLN | 895 | A | ASN | 1074 | C |
| PRO | 1079 | A | TYR | 917 | B |
| GLN | 920 | A | ILE | 1130 | C |
| GLY | 700 | A | TYR | 873 | B |
| VAL | 47 | A | ILE | 569 | C |
| VAL | 1128 | A | TYR | 917 | B |
| GLY | 880 | A | VAL | 705 | C |
| PHE | 970 | A | PHE | 759 | B |
| PHE | 562 | A | TYR | 38 | B |
| MET | 740 | A | PHE | 592 | C |
| PHE | 43 | A | PHE | 565 | C |
| THR | 739 | A | ARG | 319 | C |
| PHE | 592 | A | MET | 740 | B |

| | | | | | |
|---|---|---|---|---|---|
| LEU | 894 | A | ILE | 714 | C |
| ARG | 983 | A | LEU | 517 | C |
| ILE | 569 | A | LYS | 964 | B |
| THR | 696 | A | MET | 869 | B |
| HIS | 519 | A | ASP | 40 | B |
| ALA | 890 | A | VAL | 1068 | C |
| ILE | 788 | A | GLY | 700 | C |
| ALA | 668 | A | LEU | 865 | B |
| GLN | 115 | A | SER | 469 | C |
| PHE | 562 | A | PRO | 225 | B |
| VAL | 705 | A | ALA | 893 | B |
| VAL | 1040 | A | GLU | 1031 | B |
| GLN | 563 | A | PHE | 43 | B |
| ILE | 1013 | A | ILE | 1013 | B |
| GLN | 314 | A | LEU | 861 | B |
| PHE | 797 | A | TYR | 707 | C |
| TYR | 1067 | A | ALA | 890 | B |
| GLN | 895 | A | ALA | 706 | C |
| ARG | 1039 | A | ARG | 1039 | B |
| VAL | 1040 | A | GLY | 1035 | B |
| SER | 758 | A | SER | 968 | C |
| GLY | 283 | A | GLN | 563 | C |
| LYS | 964 | A | ALA | 570 | C |
| LEU | 865 | A | LEU | 699 | C |
| ILE | 788 | A | LEU | 699 | C |
| MET | 900 | A | PHE | 1095 | C |
| ILE | 882 | A | TYR | 707 | C |
| ASP | 198 | A | PRO | 426 | C |
| VAL | 382 | A | LEU | 984 | B |
| LEU | 1141 | A | LEU | 1141 | B |
| SER | 1003 | A | GLN | 762 | B |
| VAL | 705 | A | LEU | 894 | B |
| ILE | 896 | A | ILE | 712 | C |
| LYS | 41 | A | HIS | 519 | C |
| PRO | 225 | A | PHE | 562 | C |
| PHE | 759 | A | GLN | 965 | C |
| ALA | 520 | A | ASP | 228 | B |
| SER | 967 | A | ALA | 570 | C |
| ARG | 355 | A | TYR | 200 | B |
| ALA | 570 | A | SER | 975 | B |
| GLN | 762 | A | THR | 1006 | C |

| | | | | | |
|---|---|---|---|---|---|
| PHE | 759 | A | THR | 1006 | C |
| LEU | 864 | A | GLY | 669 | C |
| PRO | 863 | A | ALA | 668 | C |
| LEU | 390 | A | ARG | 983 | B |
| ARG | 765 | A | ALA | 958 | C |
| GLY | 891 | A | GLY | 1046 | C |
| LEU | 699 | A | MET | 869 | B |
| SER | 383 | A | LEU | 984 | B |
| ASP | 571 | A | SER | 968 | B |
| GLN | 115 | A | ASP | 467 | C |
| GLN | 1010 | A | GLN | 762 | B |
| LYS | 558 | A | ASN | 282 | B |
| PHE | 43 | A | GLN | 563 | C |
| ARG | 567 | A | VAL | 42 | B |
| VAL | 963 | A | ILE | 569 | C |
| HIS | 519 | A | LYS | 41 | B |
| GLU | 1144 | A | LEU | 1145 | C |
| SER | 698 | A | TYR | 873 | B |
| MET | 697 | A | LEU | 864 | B |
| ARG | 983 | A | LYS | 386 | C |
| ASP | 985 | A | SER | 383 | C |
| VAL | 382 | A | ARG | 983 | B |
| ILE | 569 | A | VAL | 963 | B |
| SER | 982 | A | ASP | 389 | C |
| THR | 768 | A | GLN | 314 | C |
| LEU | 560 | A | ASN | 282 | B |
| TYR | 789 | A | ASN | 703 | C |
| ALA | 890 | A | TYR | 1047 | C |
| VAL | 42 | A | HIS | 519 | C |
| ILE | 569 | A | SER | 967 | B |
| ASP | 571 | A | LEU | 966 | B |
| ARG | 1107 | A | ASN | 907 | B |
| GLN | 872 | A | LEU | 699 | C |
| PHE | 1089 | A | ASN | 914 | B |
| LYS | 41 | A | PHE | 565 | C |
| SER | 758 | A | GLN | 965 | C |
| TYR | 917 | A | PRO | 1079 | C |
| GLU | 918 | A | VAL | 1129 | C |
| ILE | 1013 | A | ILE | 1013 | C |
| PRO | 792 | A | VAL | 705 | C |
| GLN | 1002 | A | PHE | 759 | B |

| | | | | | |
|---|---|---|---|---|---|
| GLY | 669 | A | MET | 869 | B |
| ALA | 668 | A | PRO | 862 | B |
| PHE | 1121 | A | GLN | 913 | B |
| GLN | 1010 | A | VAL | 1008 | B |
| SER | 1030 | A | ASP | 1041 | C |
| ASP | 198 | A | ASP | 428 | C |
| SER | 704 | A | ILE | 788 | B |
| ASN | 703 | A | GLN | 787 | B |
| ILE | 1013 | A | THR | 1009 | B |
| ALA | 892 | A | GLU | 1072 | C |
| GLU | 702 | A | ILE | 788 | B |
| GLY | 545 | A | ASN | 978 | B |
| PHE | 559 | A | PHE | 43 | B |
| ASN | 907 | A | ARG | 1091 | C |
| GLN | 1002 | A | GLN | 1002 | B |
| PRO | 715 | A | LEU | 894 | B |
| ALA | 890 | A | ASP | 1041 | C |
| ALA | 890 | A | PRO | 1069 | C |
| THR | 1077 | A | PRO | 897 | B |
| ASP | 427 | A | LYS | 986 | C |
| LEU | 546 | A | ASN | 978 | B |
| ASN | 919 | A | VAL | 1128 | C |
| PHE | 562 | A | VAL | 42 | B |
| SER | 975 | A | ASP | 571 | C |
| ASN | 1074 | A | GLN | 895 | B |
| ASN | 1125 | A | GLU | 918 | B |
| THR | 549 | A | ASP | 745 | B |
| LEU | 984 | A | LYS | 386 | C |
| ILE | 788 | A | GLU | 702 | C |
| ASN | 914 | A | PHE | 1121 | C |
| TYR | 38 | A | LEU | 560 | C |
| GLN | 895 | A | VAL | 705 | C |
| PHE | 562 | A | GLU | 224 | B |
| GLY | 545 | A | SER | 982 | B |
| VAL | 1129 | A | GLU | 918 | B |
| MET | 740 | A | PRO | 589 | C |
| LYS | 1038 | A | LYS | 1038 | B |
| LEU | 1141 | A | PRO | 1140 | B |
| LEU | 864 | A | CYS | 671 | C |
| THR | 791 | A | VAL | 705 | C |
| LEU | 1141 | A | GLU | 1144 | B |

| | | | | | |
|---|---|---|---|---|---|
| ASN | 709 | A | PRO | 897 | B |
| LEU | 981 | A | THR | 547 | C |
| TYR | 917 | A | PHE | 1089 | C |
| ILE | 896 | A | TYR | 707 | C |
| GLY | 566 | A | ARG | 44 | B |
| VAL | 1128 | A | GLU | 918 | B |
| GLN | 1005 | A | THR | 1006 | C |
| ASP | 985 | A | LYS | 386 | C |
| VAL | 1040 | A | LEU | 1034 | B |
| VAL | 1122 | A | GLN | 1113 | B |
| ILE | 712 | A | TYR | 904 | B |
| VAL | 976 | A | ASP | 571 | C |
| MET | 900 | A | VAL | 1094 | C |
| THR | 302 | A | THR | 761 | B |
| LEU | 861 | A | GLN | 613 | C |
| GLY | 889 | A | LYS | 1045 | C |
| GLN | 779 | A | MET | 697 | C |
| GLY | 1035 | A | VAL | 1040 | C |
| GLN | 414 | A | ASP | 985 | C |
| ARG | 646 | A | PRO | 862 | B |
| VAL | 1040 | A | GLY | 889 | B |
| ASP | 389 | A | SER | 982 | B |
| THR | 866 | A | GLY | 669 | C |
| LEU | 560 | A | GLU | 224 | B |
| LEU | 864 | A | MET | 697 | C |
| GLN | 1036 | A | VAL | 1040 | C |
| SER | 735 | A | GLN | 314 | C |
| ARG | 319 | A | ASP | 745 | B |
| PHE | 1042 | A | SER | 1030 | B |
| ASN | 282 | A | LYS | 558 | C |
| PHE | 1095 | A | MET | 900 | B |
| PRO | 561 | A | GLU | 224 | B |
| VAL | 1040 | A | GLN | 1036 | B |
| PHE | 759 | A | SER | 1003 | C |
| GLN | 965 | A | SER | 758 | B |
| TYR | 917 | A | VAL | 1129 | C |
| SER | 704 | A | LYS | 790 | B |
| LYS | 1045 | A | LYS | 786 | B |
| ASN | 317 | A | ASP | 737 | B |
| ASP | 1041 | A | ALA | 890 | B |
| GLY | 669 | A | PRO | 863 | B |

| | | | | | |
|---|---|---|---|---|---|
| GLU | 702 | A | TYR | 789 | B |
| ARG | 357 | A | PRO | 230 | B |
| GLN | 115 | A | ILE | 468 | C |
| PRO | 1090 | A | ASN | 907 | B |
| ASN | 914 | A | PHE | 1089 | C |
| SER | 711 | A | ILE | 896 | B |
| VAL | 705 | A | SER | 884 | B |
| LEU | 864 | A | CYS | 662 | C |
| PHE | 565 | A | LYS | 41 | B |
| PRO | 384 | A | ASP | 985 | B |
| MET | 697 | A | GLN | 779 | B |
| THR | 791 | A | SER | 704 | C |
| GLN | 1002 | A | GLN | 1005 | B |
| MET | 869 | A | LEU | 699 | C |
| GLN | 613 | A | LEU | 861 | B |
| GLY | 971 | A | GLN | 755 | B |
| TRP | 886 | A | ARG | 1107 | C |
| ILE | 666 | A | LEU | 864 | B |
| SER | 711 | A | GLN | 895 | B |
| SER | 1123 | A | GLU | 918 | B |
| PHE | 562 | A | ASP | 40 | B |
| ILE | 788 | A | SER | 704 | C |
| ASN | 969 | A | TYR | 756 | B |
| ALA | 890 | A | TYR | 1067 | C |
| GLU | 1031 | A | VAL | 1040 | C |
| GLN | 1002 | A | LEU | 1001 | B |
| LYS | 786 | A | LEU | 699 | C |
| GLY | 199 | A | PRO | 463 | C |
| ILE | 233 | A | ARG | 466 | C |
| GLN | 1002 | A | GLN | 1002 | C |
| PHE | 43 | A | ARG | 567 | C |
| LYS | 921 | A | ILE | 1130 | C |
| ILE | 896 | A | ALA | 713 | C |
| PRO | 862 | A | ILE | 666 | C |
| GLN | 755 | A | ALA | 972 | C |
| THR | 385 | A | LEU | 984 | B |
| PRO | 863 | A | GLY | 667 | C |
| LEU | 984 | A | SER | 383 | C |
| GLN | 563 | A | LYS | 41 | B |
| ARG | 983 | A | THR | 430 | C |
| LEU | 699 | A | GLN | 872 | B |

| | | | | | |
|---|---|---|---|---|---|
| VAL | 42 | A | ARG | 567 | C |
| LYS | 558 | A | PHE | 43 | B |
| ALA | 893 | A | GLU | 1072 | C |
| GLY | 381 | A | ILE | 973 | B |
| PHE | 592 | A | ASN | 856 | B |
| TYR | 396 | A | TYR | 200 | B |
| PRO | 862 | A | ALA | 668 | C |
| PRO | 589 | A | PHE | 855 | B |
| PRO | 897 | A | ASN | 709 | C |
| LEU | 390 | A | SER | 982 | B |
| LYS | 386 | A | SER | 982 | B |

**Table S3.** The list of the inter-protomer contacts of the protomer A in the structure of SARS-CoV-2 S-GSAS/G614 in the closed state (pdb id 7KDK).

| S protein residue name | S protein residue number | S protein chain | S protein residue name | S protein residue number | S protein chain |
|---|---|---|---|---|---|
| PHE | 562 | A | GLU | 224 | B |
| PHE | 562 | A | PRO | 225 | B |
| GLN | 755 | A | PHE | 970 | C |
| GLY | 757 | A | SER | 968 | C |
| ARG | 1039 | A | THR | 1027 | B |
| LEU | 1034 | A | VAL | 1040 | C |
| TYR | 707 | A | ILE | 794 | B |
| GLY | 700 | A | LYS | 786 | B |
| GLY | 798 | A | TYR | 707 | C |
| ALA | 713 | A | GLN | 895 | B |
| TYR | 200 | A | GLU | 516 | C |
| GLN | 895 | A | ASN | 1074 | C |
| GLY | 857 | A | ASN | 317 | C |
| ASN | 856 | A | ALA | 570 | C |
| PHE | 592 | A | THR | 859 | B |
| ASP | 994 | A | PHE | 970 | C |
| VAL | 1008 | A | GLN | 1010 | C |
| GLN | 563 | A | PHE | 43 | B |
| LYS | 1045 | A | GLY | 889 | B |
| ARG | 567 | A | VAL | 976 | B |
| CYS | 379 | A | GLU | 988 | B |
| LEU | 699 | A | ILE | 870 | B |
| ALA | 890 | A | TYR | 1047 | C |
| ILE | 1013 | A | ALA | 1016 | B |
| TYR | 707 | A | PHE | 797 | B |
| PRO | 863 | A | GLY | 667 | C |
| GLU | 702 | A | TYR | 789 | B |
| LEU | 699 | A | VAL | 785 | B |
| ALA | 1080 | A | TYR | 917 | B |
| PHE | 562 | A | VAL | 42 | B |
| SER | 711 | A | ILE | 896 | B |
| TYR | 904 | A | VAL | 1094 | C |
| GLY | 283 | A | LEU | 560 | C |
| GLU | 918 | A | PHE | 1089 | C |
| ARG | 567 | A | ARG | 44 | B |

| | | | | | |
|---|---|---|---|---|---|
| ARG | 1039 | A | ARG | 1039 | C |
| GLU | 988 | A | SER | 383 | C |
| LEU | 699 | A | LYS | 786 | B |
| PRO | 384 | A | THR | 415 | C |
| VAL | 987 | A | PRO | 412 | B |
| ILE | 1130 | A | GLY | 798 | B |
| LEU | 1145 | A | GLU | 1144 | B |
| ILE | 896 | A | ILE | 712 | C |
| GLN | 913 | A | PHE | 1089 | C |
| SER | 1003 | A | PHE | 759 | B |
| PRO | 230 | A | ARG | 357 | C |
| GLY | 700 | A | GLN | 787 | B |
| THR | 961 | A | ARG | 765 | B |
| LEU | 864 | A | PRO | 665 | C |
| VAL | 1068 | A | GLY | 891 | B |
| ALA | 892 | A | PRO | 1069 | C |
| SER | 982 | A | GLY | 545 | C |
| PHE | 464 | A | TYR | 200 | B |
| GLN | 1010 | A | GLN | 762 | B |
| PRO | 897 | A | THR | 1077 | C |
| TYR | 789 | A | SER | 704 | C |
| ASP | 1041 | A | GLY | 889 | B |
| TYR | 707 | A | ILE | 882 | B |
| GLY | 545 | A | SER | 982 | B |
| SER | 1147 | A | SER | 1147 | B |
| TYR | 707 | A | ALA | 899 | B |
| ALA | 701 | A | ILE | 788 | B |
| ASP | 979 | A | ARG | 567 | C |
| MET | 697 | A | MET | 869 | B |
| TYR | 789 | A | VAL | 705 | C |
| ILE | 468 | A | GLN | 115 | B |
| GLY | 669 | A | THR | 866 | B |
| MET | 697 | A | LEU | 865 | B |
| VAL | 382 | A | LEU | 984 | B |
| GLN | 115 | A | ILE | 468 | C |
| GLN | 1010 | A | ALA | 766 | B |
| GLY | 669 | A | MET | 869 | B |
| GLY | 971 | A | GLN | 755 | B |
| LYS | 795 | A | TYR | 707 | C |
| GLY | 232 | A | GLU | 465 | C |
| THR | 859 | A | PHE | 592 | C |

| | | | | | |
|---|---|---|---|---|---|
| GLN | 1002 | A | LEU | 1001 | B |
| ARG | 983 | A | LEU | 390 | C |
| ALA | 713 | A | LEU | 894 | B |
| ASP | 1041 | A | GLN | 784 | B |
| ASP | 985 | A | THR | 415 | B |
| GLY | 889 | A | ASP | 1041 | C |
| PRO | 589 | A | PHE | 855 | B |
| PRO | 863 | A | ALA | 668 | C |
| GLN | 1005 | A | THR | 1006 | C |
| LEU | 560 | A | THR | 284 | B |
| LEU | 861 | A | GLN | 613 | C |
| ALA | 890 | A | PRO | 1069 | C |
| ARG | 567 | A | PHE | 43 | B |
| SER | 383 | A | LEU | 984 | B |
| GLU | 918 | A | SER | 1123 | C |
| THR | 415 | A | TYR | 369 | B |
| VAL | 705 | A | SER | 884 | B |
| GLN | 1002 | A | GLN | 1002 | C |
| ASN | 914 | A | PHE | 1121 | C |
| LEU | 1145 | A | LEU | 1145 | B |
| PHE | 592 | A | LEU | 858 | B |
| ALA | 766 | A | GLN | 1010 | C |
| MET | 869 | A | ALA | 668 | C |
| ASP | 198 | A | PHE | 464 | C |
| SER | 968 | A | GLY | 757 | B |
| LYS | 786 | A | LYS | 1045 | C |
| ASP | 985 | A | LYS | 386 | C |
| MET | 869 | A | GLY | 669 | C |
| TYR | 917 | A | ILE | 1130 | C |
| TYR | 789 | A | ASN | 703 | C |
| THR | 572 | A | ASN | 856 | B |
| ASP | 198 | A | ASP | 428 | C |
| MET | 869 | A | THR | 696 | C |
| SER | 704 | A | ILE | 788 | B |
| LYS | 776 | A | LYS | 947 | C |
| ILE | 788 | A | ASN | 703 | C |
| GLN | 895 | A | VAL | 705 | C |
| ARG | 646 | A | THR | 866 | B |
| ASP | 568 | A | PHE | 855 | B |
| LEU | 864 | A | CYS | 662 | C |
| GLN | 895 | A | ALA | 706 | C |

| | | | | | |
|---|---|---|---|---|---|
| ARG | 44 | A | GLN | 563 | C |
| GLN | 895 | A | SER | 708 | C |
| ILE | 569 | A | ASN | 960 | B |
| ASP | 428 | A | ASP | 198 | B |
| GLY | 1035 | A | VAL | 1040 | C |
| ALA | 570 | A | SER | 975 | B |
| SER | 711 | A | GLN | 895 | B |
| LEU | 984 | A | VAL | 382 | C |
| GLY | 757 | A | GLN | 965 | C |
| ARG | 319 | A | ASP | 745 | B |
| GLN | 564 | A | LYS | 41 | B |
| ASP | 571 | A | SER | 967 | B |
| LEU | 966 | A | ASP | 571 | C |
| GLU | 1031 | A | PHE | 1042 | C |
| ARG | 466 | A | GLN | 115 | B |
| VAL | 705 | A | PRO | 792 | B |
| PHE | 855 | A | ASP | 568 | C |
| ALA | 570 | A | ASN | 856 | B |
| THR | 430 | A | ARG | 983 | B |
| ASP | 796 | A | TYR | 707 | C |
| PRO | 230 | A | TYR | 396 | C |
| ASP | 571 | A | ARG | 44 | B |
| ILE | 231 | A | ARG | 466 | C |
| ALA | 668 | A | PRO | 862 | B |
| ASN | 703 | A | GLN | 787 | B |
| ARG | 1107 | A | ASN | 907 | B |
| VAL | 1129 | A | TYR | 917 | B |
| LYS | 41 | A | ALA | 520 | C |
| GLN | 1002 | A | TYR | 756 | B |
| GLN | 920 | A | ILE | 1130 | C |
| LEU | 560 | A | GLY | 283 | B |
| SER | 982 | A | THR | 547 | C |
| GLU | 516 | A | ASP | 228 | B |
| GLN | 563 | A | TYR | 38 | B |
| TYR | 917 | A | VAL | 1128 | C |
| SER | 704 | A | LYS | 790 | B |
| ALA | 701 | A | GLN | 787 | B |
| ASN | 764 | A | GLN | 314 | C |
| ASN | 856 | A | THR | 572 | C |
| MET | 869 | A | LEU | 699 | C |
| ARG | 983 | A | LYS | 386 | C |

| | | | | | |
|---|---|---|---|---|---|
| MET | 869 | A | ILE | 670 | C |
| GLN | 755 | A | ALA | 972 | C |
| GLU | 224 | A | PRO | 561 | C |
| LYS | 790 | A | ASN | 703 | C |
| TYR | 396 | A | PRO | 230 | B |
| ILE | 712 | A | ILE | 896 | B |
| LEU | 1012 | A | ILE | 1013 | C |
| ILE | 788 | A | LEU | 699 | C |
| GLY | 1093 | A | GLN | 913 | B |
| PHE | 43 | A | PHE | 565 | C |
| ARG | 983 | A | GLY | 381 | C |
| GLU | 1111 | A | SER | 1123 | C |
| TRP | 886 | A | ASN | 1108 | C |
| ASN | 914 | A | PHE | 1089 | C |
| SER | 698 | A | TYR | 873 | B |
| ALA | 713 | A | ILE | 896 | B |
| ARG | 319 | A | MET | 740 | B |
| TYR | 1047 | A | THR | 887 | B |
| ASP | 571 | A | VAL | 976 | B |
| THR | 1077 | A | MET | 900 | B |
| GLN | 314 | A | ASN | 764 | B |
| HIS | 519 | A | ARG | 983 | B |
| GLY | 199 | A | GLU | 465 | C |
| GLY | 667 | A | PRO | 863 | B |
| LEU | 1141 | A | LEU | 1141 | B |
| SER | 708 | A | PRO | 897 | B |
| ILE | 714 | A | LEU | 894 | B |
| LEU | 699 | A | TYR | 873 | B |
| PRO | 862 | A | GLY | 667 | C |
| TYR | 756 | A | GLN | 1002 | C |
| GLY | 667 | A | LEU | 864 | B |
| LEU | 865 | A | LEU | 699 | C |
| GLY | 669 | A | PRO | 863 | B |
| TYR | 789 | A | GLU | 702 | C |
| VAL | 42 | A | GLN | 563 | C |
| GLN | 913 | A | PRO | 1079 | C |
| LYS | 41 | A | PHE | 565 | C |
| ALA | 1078 | A | TYR | 917 | B |
| LYS | 386 | A | ARG | 983 | B |
| GLY | 548 | A | ASN | 978 | B |
| LYS | 378 | A | GLU | 988 | B |

| | | | | | |
|---|---|---|---|---|---|
| MET | 697 | A | TYR | 873 | B |
| PHE | 1089 | A | GLU | 918 | B |
| LEU | 966 | A | ALA | 570 | C |
| ALA | 890 | A | ASP | 1041 | C |
| SER | 1123 | A | GLU | 1111 | B |
| LEU | 560 | A | TYR | 38 | B |
| LYS | 202 | A | LEU | 518 | C |
| ARG | 983 | A | VAL | 382 | C |
| PRO | 897 | A | ILE | 712 | C |
| THR | 547 | A | ASN | 978 | B |
| GLU | 516 | A | TYR | 200 | B |
| SER | 968 | A | TYR | 756 | B |
| VAL | 1094 | A | MET | 900 | B |
| SER | 975 | A | ARG | 567 | C |
| TYR | 917 | A | VAL | 1129 | C |
| GLN | 895 | A | ALA | 713 | C |
| TYR | 369 | A | ARG | 408 | C |
| ASP | 745 | A | ARG | 319 | C |
| ARG | 466 | A | ILE | 233 | B |
| PHE | 562 | A | LYS | 41 | B |
| GLU | 988 | A | CYS | 379 | C |
| GLU | 1072 | A | ALA | 893 | B |
| SER | 704 | A | TYR | 789 | B |
| ASN | 234 | A | ARG | 466 | C |
| SER | 1123 | A | GLU | 918 | B |
| GLN | 1010 | A | LEU | 1012 | B |
| LYS | 558 | A | PHE | 43 | B |
| GLN | 563 | A | ARG | 44 | B |
| LEU | 864 | A | ILE | 670 | C |
| ARG | 319 | A | ASP | 737 | B |
| VAL | 1040 | A | GLN | 1036 | B |
| MET | 900 | A | VAL | 1094 | C |
| PHE | 592 | A | MET | 740 | B |
| ARG | 355 | A | TYR | 200 | B |
| LYS | 790 | A | GLU | 702 | C |
| VAL | 42 | A | PHE | 565 | C |
| ALA | 570 | A | LYS | 964 | B |
| LEU | 894 | A | ILE | 714 | C |
| PHE | 592 | A | LYS | 854 | B |
| ARG | 1107 | A | TRP | 886 | B |
| VAL | 963 | A | ALA | 570 | C |

| | | | | | |
|---|---|---|---|---|---|
| GLN | 762 | A | TYR | 1007 | C |
| TYR | 707 | A | GLN | 895 | B |
| GLN | 787 | A | GLY | 700 | C |
| ASN | 317 | A | MET | 740 | B |
| TYR | 707 | A | GLY | 798 | B |
| TYR | 1007 | A | GLN | 762 | B |
| ILE | 569 | A | VAL | 47 | B |
| LYS | 1038 | A | GLN | 1036 | B |
| THR | 961 | A | SER | 758 | B |
| PRO | 1090 | A | GLN | 913 | B |
| ALA | 893 | A | VAL | 705 | C |
| TYR | 756 | A | PHE | 970 | C |
| GLY | 857 | A | PHE | 592 | C |
| ALA | 1016 | A | ILE | 1013 | C |
| PHE | 970 | A | PHE | 759 | B |
| ASN | 978 | A | GLY | 548 | C |
| PHE | 464 | A | GLY | 199 | B |
| ILE | 788 | A | SER | 704 | C |
| GLN | 965 | A | PHE | 759 | B |
| VAL | 705 | A | ALA | 879 | B |
| ASN | 709 | A | PRO | 897 | B |
| THR | 385 | A | ASP | 985 | B |
| VAL | 382 | A | GLU | 988 | B |
| PRO | 521 | A | LYS | 41 | B |
| ASN | 914 | A | SER | 1123 | C |
| THR | 866 | A | ALA | 668 | C |
| PRO | 1069 | A | LEU | 894 | B |
| ILE | 788 | A | ALA | 701 | C |
| VAL | 963 | A | ILE | 569 | C |
| LEU | 390 | A | SER | 982 | B |
| TYR | 873 | A | MET | 697 | C |
| CYS | 662 | A | LEU | 864 | B |
| ASN | 1074 | A | GLN | 895 | B |
| THR | 415 | A | THR | 385 | B |
| ALA | 890 | A | LYS | 1045 | C |
| SER | 383 | A | GLU | 988 | B |
| ALA | 520 | A | LYS | 41 | B |
| GLN | 1002 | A | GLN | 1005 | B |
| GLN | 755 | A | GLY | 971 | C |
| ASP | 1041 | A | LEU | 1034 | B |
| LYS | 921 | A | ILE | 1130 | C |

| | | | | | |
|---|---|---|---|---|---|
| GLU | 702 | A | ILE | 788 | B |
| LEU | 699 | A | ILE | 788 | B |
| PHE | 759 | A | THR | 1006 | C |
| ILE | 1130 | A | TYR | 917 | B |
| ASP | 427 | A | LYS | 986 | C |
| SER | 975 | A | ASP | 571 | C |
| TRP | 886 | A | ARG | 1107 | C |
| PHE | 759 | A | GLN | 1002 | C |
| ASN | 317 | A | ASP | 737 | B |
| GLY | 891 | A | LYS | 1045 | C |
| LEU | 1001 | A | GLN | 1002 | C |
| ALA | 890 | A | VAL | 1068 | C |
| VAL | 1094 | A | ILE | 896 | B |
| LEU | 546 | A | ASN | 978 | B |
| TYR | 38 | A | LEU | 560 | C |
| ARG | 983 | A | SER | 383 | C |
| VAL | 1128 | A | ASN | 919 | B |
| TYR | 873 | A | LEU | 699 | C |
| GLN | 957 | A | ARG | 765 | B |
| ARG | 1019 | A | GLU | 1017 | C |
| ALA | 372 | A | ARG | 403 | C |
| GLY | 880 | A | VAL | 705 | C |
| GLU | 988 | A | PRO | 384 | C |
| ALA | 1016 | A | ARG | 1019 | B |
| TYR | 756 | A | GLN | 965 | C |
| GLN | 762 | A | GLN | 965 | C |
| TYR | 707 | A | PRO | 897 | B |
| MET | 900 | A | PRO | 1079 | C |
| LYS | 790 | A | VAL | 705 | C |
| GLN | 613 | A | LEU | 861 | B |
| ILE | 896 | A | TYR | 707 | C |
| PHE | 565 | A | PHE | 43 | B |
| PHE | 592 | A | PHE | 855 | B |
| GLY | 798 | A | ILE | 1130 | C |
| PRO | 863 | A | GLY | 669 | C |
| ASN | 1108 | A | TRP | 886 | B |
| LYS | 790 | A | SER | 704 | C |
| ILE | 666 | A | LEU | 864 | B |
| LEU | 1141 | A | LEU | 1141 | C |
| VAL | 1129 | A | GLU | 918 | B |
| LYS | 41 | A | PHE | 562 | C |

| | | | | | |
|---|---|---|---|---|---|
| THR | 1006 | A | GLN | 762 | B |
| ASP | 571 | A | SER | 975 | B |
| ILE | 896 | A | SER | 711 | C |
| ASP | 427 | A | VAL | 987 | C |
| TYR | 1067 | A | ALA | 890 | B |
| THR | 883 | A | TYR | 707 | C |
| PRO | 862 | A | ALA | 647 | C |
| ILE | 1013 | A | THR | 1009 | B |
| VAL | 1040 | A | LEU | 1034 | B |
| ASN | 907 | A | ARG | 1107 | C |
| PRO | 715 | A | LEU | 894 | B |
| GLN | 1010 | A | VAL | 1008 | B |
| ILE | 870 | A | LEU | 699 | C |
| TYR | 200 | A | ARG | 355 | C |
| TYR | 873 | A | GLY | 700 | C |
| GLN | 563 | A | TYR | 279 | B |
| SER | 1030 | A | ASP | 1041 | C |
| ALA | 713 | A | ALA | 893 | B |
| PRO | 1079 | A | MET | 900 | B |
| SER | 968 | A | GLN | 755 | B |
| ILE | 712 | A | PRO | 897 | B |
| ASP | 985 | A | THR | 385 | C |
| GLU | 1072 | A | ALA | 892 | B |
| GLN | 755 | A | ASN | 969 | C |
| LEU | 984 | A | LYS | 386 | C |
| GLY | 232 | A | ARG | 466 | C |
| GLY | 566 | A | ARG | 44 | B |
| PHE | 797 | A | TYR | 707 | C |
| LEU | 894 | A | PRO | 1069 | C |
| LEU | 699 | A | MET | 869 | B |
| PRO | 897 | A | SER | 708 | C |
| LEU | 894 | A | ARG | 1107 | C |
| VAL | 1128 | A | TYR | 917 | B |
| PHE | 559 | A | PHE | 43 | B |
| VAL | 1040 | A | GLU | 1031 | B |
| SER | 758 | A | SER | 968 | C |
| ILE | 1013 | A | ILE | 1013 | C |
| PRO | 463 | A | GLY | 199 | B |
| THR | 1077 | A | PRO | 897 | B |
| ARG | 646 | A | PRO | 863 | B |
| GLN | 913 | A | GLY | 1093 | C |

| | | | | | |
|---|---|---|---|---|---|
| GLN | 613 | A | THR | 859 | B |
| ARG | 1091 | A | ASN | 907 | B |
| PRO | 230 | A | ARG | 355 | C |
| ILE | 569 | A | LYS | 964 | B |
| PRO | 589 | A | MET | 740 | B |
| MET | 900 | A | PHE | 1095 | C |
| ALA | 570 | A | LEU | 966 | B |
| PHE | 759 | A | GLY | 999 | C |
| LEU | 517 | A | ARG | 983 | B |
| GLN | 1002 | A | GLN | 1002 | B |
| GLY | 889 | A | VAL | 1040 | C |
| ARG | 1039 | A | GLU | 1031 | B |
| VAL | 1068 | A | ALA | 890 | B |
| GLN | 965 | A | GLN | 762 | B |
| LEU | 865 | A | GLY | 669 | C |
| GLY | 413 | A | VAL | 987 | C |
| GLY | 700 | A | TYR | 873 | B |
| MET | 740 | A | ASN | 317 | C |
| GLN | 314 | A | THR | 768 | B |
| GLY | 891 | A | GLY | 1046 | C |
| LEU | 699 | A | LEU | 865 | B |
| THR | 547 | A | SER | 982 | B |
| GLU | 1017 | A | ARG | 1019 | B |
| TYR | 1047 | A | ALA | 890 | B |
| GLN | 965 | A | SER | 758 | B |
| VAL | 705 | A | LEU | 894 | B |
| GLY | 700 | A | ILE | 788 | B |
| ALA | 701 | A | TYR | 789 | B |
| LYS | 964 | A | ILE | 569 | C |
| PHE | 1121 | A | THR | 912 | B |
| LEU | 865 | A | ALA | 668 | C |
| SER | 982 | A | LYS | 386 | C |
| SER | 982 | A | LEU | 390 | C |
| ALA | 893 | A | ALA | 713 | C |
| PRO | 862 | A | ARG | 646 | C |
| ASN | 282 | A | LYS | 558 | C |
| PHE | 1089 | A | TYR | 917 | B |
| LEU | 864 | A | ALA | 668 | C |
| GLN | 1036 | A | LYS | 1038 | C |
| ARG | 567 | A | VAL | 42 | B |
| MET | 697 | A | GLN | 779 | B |

| | | | | | |
|---|---|---|---|---|---|
| ILE | 670 | A | LEU | 864 | B |
| SER | 735 | A | GLN | 314 | C |
| PHE | 970 | A | ASP | 994 | B |
| LYS | 558 | A | ASN | 282 | B |
| GLN | 895 | A | ILE | 712 | C |
| VAL | 42 | A | HIS | 519 | C |
| ILE | 233 | A | ARG | 466 | C |
| ARG | 408 | A | TYR | 369 | B |
| GLN | 762 | A | THR | 1006 | C |
| ILE | 666 | A | PRO | 862 | B |
| PRO | 463 | A | ASP | 198 | B |
| HIS | 519 | A | ASP | 979 | B |
| GLY | 999 | A | PHE | 759 | B |
| VAL | 1040 | A | GLY | 1035 | B |
| MET | 869 | A | MET | 697 | C |
| PRO | 862 | A | ALA | 668 | C |
| LYS | 986 | A | ASP | 427 | B |
| ASP | 796 | A | ASN | 709 | C |
| PHE | 43 | A | PHE | 559 | C |
| ILE | 712 | A | GLN | 895 | B |
| VAL | 47 | A | ILE | 569 | C |
| SER | 758 | A | LYS | 964 | C |
| VAL | 1094 | A | TYR | 904 | B |
| ARG | 357 | A | PRO | 230 | B |
| ARG | 1107 | A | LEU | 894 | B |
| PHE | 43 | A | GLY | 566 | C |
| ALA | 647 | A | PRO | 862 | B |
| LEU | 858 | A | PHE | 592 | C |
| THR | 1009 | A | THR | 1009 | C |
| GLU | 702 | A | LYS | 790 | B |
| ASN | 978 | A | LEU | 546 | C |
| PHE | 898 | A | TYR | 707 | C |
| TYR | 38 | A | GLN | 563 | C |
| PRO | 412 | A | VAL | 987 | C |
| GLU | 465 | A | ILE | 233 | B |
| SER | 514 | A | TYR | 200 | B |
| GLN | 784 | A | ASP | 1041 | C |
| LEU | 1141 | A | PRO | 1140 | B |
| GLY | 669 | A | LEU | 864 | B |
| ARG | 1107 | A | TYR | 904 | B |
| VAL | 705 | A | TYR | 789 | B |

| | | | | | |
|---|---|---|---|---|---|
| LEU | 1145 | A | LEU | 1141 | B |
| THR | 883 | A | VAL | 705 | C |
| ALA | 879 | A | VAL | 705 | C |
| PRO | 897 | A | ASN | 709 | C |
| THR | 696 | A | MET | 869 | B |
| LEU | 560 | A | PHE | 43 | B |
| GLN | 563 | A | ASP | 40 | B |
| ASP | 796 | A | SER | 708 | C |
| LYS | 41 | A | GLN | 564 | C |
| PRO | 665 | A | LEU | 864 | B |
| ASP | 228 | A | GLU | 516 | C |
| PRO | 384 | A | GLU | 988 | B |
| GLU | 1017 | A | GLU | 773 | B |
| ASP | 979 | A | LEU | 546 | C |
| VAL | 42 | A | PHE | 562 | C |
| SER | 758 | A | GLN | 965 | C |
| LYS | 386 | A | SER | 982 | B |
| ALA | 668 | A | MET | 869 | B |
| VAL | 382 | A | ARG | 983 | B |
| GLU | 1072 | A | LEU | 894 | B |
| TYR | 396 | A | TYR | 200 | B |
| ASP | 40 | A | HIS | 519 | C |
| THR | 1009 | A | THR | 1009 | B |
| LEU | 865 | A | MET | 697 | C |
| TYR | 369 | A | GLY | 416 | C |
| PHE | 970 | A | GLN | 755 | B |
| GLU | 1031 | A | VAL | 1040 | C |
| CYS | 671 | A | LEU | 864 | B |
| TYR | 917 | A | PHE | 1089 | C |
| VAL | 976 | A | THR | 572 | C |
| ARG | 1019 | A | ILE | 1013 | C |
| PHE | 970 | A | GLY | 757 | B |
| ARG | 646 | A | PRO | 862 | B |
| ILE | 788 | A | GLY | 700 | C |
| PRO | 1069 | A | ALA | 892 | B |
| ILE | 712 | A | TYR | 904 | B |
| GLN | 779 | A | MET | 697 | C |
| PHE | 592 | A | ASN | 856 | B |
| ARG | 319 | A | THR | 739 | B |
| GLN | 563 | A | GLY | 283 | B |
| VAL | 976 | A | ASP | 571 | C |

| | | | | | |
|---|---|---|---|---|---|
| PRO | 426 | A | ASP | 198 | B |
| THR | 385 | A | THR | 415 | C |
| ALA | 890 | A | GLY | 1046 | C |
| SER | 708 | A | GLN | 895 | B |
| ASN | 703 | A | TYR | 789 | B |
| GLY | 667 | A | PRO | 862 | B |
| GLY | 1124 | A | GLU | 918 | B |
| GLY | 669 | A | LEU | 865 | B |
| ALA | 892 | A | GLU | 1072 | C |
| PHE | 759 | A | GLN | 965 | C |
| TYR | 904 | A | ARG | 1107 | C |
| VAL | 976 | A | ARG | 567 | C |
| ARG | 466 | A | THR | 167 | B |
| PHE | 43 | A | LYS | 558 | C |
| THR | 1006 | A | PHE | 759 | B |
| GLY | 566 | A | PHE | 43 | B |
| THR | 1027 | A | ARG | 1039 | C |
| GLN | 1036 | A | VAL | 1040 | C |
| ILE | 896 | A | SER | 708 | C |
| VAL | 705 | A | LYS | 790 | B |
| LYS | 557 | A | PHE | 43 | B |
| THR | 887 | A | ASN | 1108 | C |
| SER | 967 | A | ASP | 571 | C |
| SER | 968 | A | SER | 758 | B |
| GLU | 988 | A | VAL | 382 | C |
| TYR | 789 | A | ALA | 701 | C |
| LEU | 984 | A | SER | 383 | C |
| GLY | 891 | A | PRO | 1069 | C |
| LEU | 894 | A | PRO | 715 | C |
| LYS | 964 | A | SER | 758 | B |
| LEU | 864 | A | MET | 697 | C |
| ILE | 569 | A | SER | 967 | B |
| ASP | 198 | A | PRO | 463 | C |
| SER | 967 | A | ALA | 570 | C |
| PHE | 464 | A | ASP | 198 | B |
| ILE | 1130 | A | LYS | 921 | B |
| ARG | 1107 | A | GLN | 913 | B |
| PHE | 592 | A | GLY | 857 | B |
| TYR | 707 | A | PHE | 898 | B |
| GLN | 895 | A | TYR | 707 | C |
| VAL | 705 | A | GLN | 895 | B |

| | | | | | |
|---|---|---|---|---|---|
| GLY | 1046 | A | ALA | 890 | B |
| LEU | 390 | A | ARG | 983 | B |
| PHE | 1042 | A | GLU | 1031 | B |
| GLU | 918 | A | VAL | 1128 | C |
| VAL | 705 | A | THR | 883 | B |
| GLY | 889 | A | GLY | 1046 | C |
| THR | 912 | A | PHE | 1121 | C |
| VAL | 987 | A | ASP | 427 | B |
| MET | 900 | A | THR | 1077 | C |
| LEU | 894 | A | ASN | 1108 | C |
| PRO | 897 | A | SER | 711 | C |
| ASP | 40 | A | GLN | 563 | C |
| VAL | 1128 | A | GLU | 918 | B |
| GLN | 613 | A | SER | 735 | B |
| ILE | 1013 | A | ILE | 1013 | B |
| GLN | 563 | A | VAL | 42 | B |
| HIS | 519 | A | VAL | 42 | B |
| VAL | 1040 | A | GLY | 889 | B |
| THR | 1006 | A | GLN | 1005 | B |
| ILE | 896 | A | VAL | 1094 | C |
| TYR | 279 | A | GLN | 563 | C |
| GLN | 755 | A | SER | 968 | C |
| LYS | 557 | A | SER | 45 | B |
| GLN | 563 | A | LYS | 41 | B |
| ILE | 1013 | A | ARG | 1019 | B |
| ALA | 890 | A | TYR | 1067 | C |
| SER | 758 | A | THR | 961 | C |
| ASP | 198 | A | PRO | 426 | C |
| PRO | 792 | A | VAL | 705 | C |
| LEU | 1145 | A | SER | 1147 | B |
| TYR | 707 | A | PRO | 792 | B |
| GLN | 913 | A | PRO | 1090 | C |
| TYR | 1047 | A | TRP | 886 | B |
| GLY | 1093 | A | TYR | 904 | B |
| GLN | 314 | A | SER | 735 | B |
| GLY | 1046 | A | GLY | 891 | B |
| PHE | 970 | A | TYR | 756 | B |
| PHE | 43 | A | ARG | 567 | C |
| PRO | 792 | A | TYR | 707 | C |
| ARG | 1039 | A | ARG | 1039 | B |
| LYS | 386 | A | LEU | 981 | B |

| | | | | | |
|---|---|---|---|---|---|
| GLN | 872 | A | LEU | 699 | C |
| ASN | 856 | A | ASP | 568 | C |
| GLN | 1002 | A | THR | 998 | B |
| ASN | 710 | A | PRO | 897 | B |
| ASP | 1041 | A | ALA | 890 | B |
| ARG | 466 | A | GLY | 232 | B |
| GLN | 314 | A | LEU | 861 | B |
| PRO | 1079 | A | TYR | 917 | B |
| GLY | 381 | A | ARG | 983 | B |
| TYR | 917 | A | ALA | 1080 | C |
| LEU | 864 | A | GLY | 669 | C |
| ARG | 983 | A | THR | 430 | C |
| SER | 383 | A | ARG | 983 | B |
| GLY | 1046 | A | GLY | 889 | B |
| LEU | 864 | A | ILE | 666 | C |
| ASP | 1146 | A | SER | 1147 | B |
| LEU | 864 | A | GLY | 667 | C |
| ILE | 882 | A | TYR | 707 | C |
| THR | 739 | A | ARG | 319 | C |
| LEU | 1012 | A | GLN | 1010 | C |
| GLN | 787 | A | ASN | 703 | C |
| LEU | 1141 | A | GLU | 1144 | B |
| ASN | 978 | A | THR | 547 | C |
| GLY | 381 | A | LEU | 984 | B |
| LYS | 41 | A | GLN | 563 | C |
| THR | 866 | A | GLY | 669 | C |
| MET | 900 | A | ALA | 1078 | C |
| GLN | 965 | A | TYR | 756 | B |
| PHE | 565 | A | LYS | 41 | B |
| ALA | 668 | A | THR | 866 | B |
| LYS | 41 | A | HIS | 519 | C |
| TYR | 170 | A | ARG | 357 | C |
| PRO | 225 | A | PHE | 562 | C |
| ALA | 668 | A | PRO | 863 | B |
| ARG | 983 | A | LEU | 517 | C |
| ASN | 907 | A | ARG | 1091 | C |
| ARG | 44 | A | GLY | 566 | C |
| ILE | 794 | A | TYR | 707 | C |
| SER | 1123 | A | ASN | 914 | B |
| TYR | 200 | A | PHE | 464 | C |
| ARG | 403 | A | ALA | 372 | B |

| | | | | | |
|---|---|---|---|---|---|
| GLU | 918 | A | GLY | 1124 | C |
| PHE | 43 | A | LYS | 557 | C |
| SER | 884 | A | VAL | 705 | C |
| PHE | 464 | A | ILE | 233 | B |
| ASN | 703 | A | ILE | 788 | B |
| LEU | 894 | A | GLU | 1072 | C |
| PHE | 1121 | A | ASN | 914 | B |
| GLN | 762 | A | THR | 961 | C |
| VAL | 705 | A | GLY | 880 | B |
| LYS | 854 | A | PHE | 592 | C |
| ILE | 712 | A | MET | 900 | B |
| PHE | 1089 | A | GLN | 913 | B |
| GLU | 702 | A | GLN | 787 | B |
| LYS | 1045 | A | LEU | 1034 | B |
| THR | 887 | A | TYR | 1047 | C |
| TYR | 200 | A | SER | 514 | C |
| TYR | 369 | A | THR | 415 | C |
| GLU | 224 | A | PHE | 562 | C |
| THR | 1009 | A | ILE | 1013 | C |
| ASN | 969 | A | GLN | 755 | B |
| PHE | 464 | A | GLY | 232 | B |
| VAL | 1040 | A | SER | 1030 | B |
| SER | 1030 | A | VAL | 1040 | C |
| GLY | 891 | A | VAL | 1068 | C |
| ALA | 570 | A | SER | 967 | B |
| THR | 961 | A | GLN | 762 | B |
| GLN | 913 | A | PHE | 1121 | C |
| PHE | 855 | A | PRO | 589 | C |
| GLU | 1031 | A | ARG | 1039 | C |
| LEU | 894 | A | VAL | 705 | C |
| LEU | 894 | A | ALA | 713 | C |
| ASN | 317 | A | ASP | 745 | B |
| SER | 708 | A | ASP | 796 | B |
| LYS | 964 | A | ALA | 570 | C |
| ARG | 765 | A | GLN | 957 | C |
| ILE | 569 | A | VAL | 963 | B |
| GLU | 465 | A | GLY | 232 | B |
| PHE | 1121 | A | GLN | 913 | B |
| MET | 697 | A | LEU | 864 | B |
| PHE | 855 | A | PHE | 592 | C |
| GLN | 1005 | A | GLN | 1002 | C |

| | | | | | |
|---|---|---|---|---|---|
| ASN | 703 | A | LYS | 790 | B |
| ASP | 571 | A | HIS | 49 | B |
| VAL | 42 | A | ARG | 567 | C |
| ARG | 765 | A | THR | 961 | C |
| ARG | 357 | A | TYR | 170 | B |
| LYS | 1045 | A | GLN | 784 | B |
| GLY | 232 | A | PHE | 464 | C |
| ILE | 1013 | A | LEU | 1012 | B |
| LEU | 984 | A | GLY | 381 | C |
| PHE | 759 | A | SER | 1003 | C |
| GLU | 988 | A | LYS | 378 | C |
| THR | 415 | A | ASP | 985 | C |
| PRO | 1069 | A | GLY | 891 | B |
| GLU | 773 | A | GLU | 1017 | C |
| PHE | 562 | A | ASP | 40 | B |
| LEU | 1034 | A | ASP | 1041 | C |
| SER | 708 | A | ILE | 896 | B |
| ALA | 570 | A | VAL | 963 | B |
| VAL | 987 | A | GLY | 413 | B |
| LYS | 786 | A | ALA | 701 | C |
| PHE | 565 | A | VAL | 42 | B |
| VAL | 42 | A | GLY | 566 | C |
| PRO | 1079 | A | GLN | 913 | B |
| MET | 740 | A | PHE | 592 | C |
| ASN | 919 | A | VAL | 1128 | C |
| PRO | 897 | A | ASN | 710 | C |
| TYR | 756 | A | SER | 968 | C |
| PRO | 1069 | A | ALA | 890 | B |
| TYR | 707 | A | ASP | 796 | B |
| ALA | 1078 | A | MET | 900 | B |
| GLY | 757 | A | PHE | 970 | C |
| ASN | 709 | A | ASP | 796 | B |
| VAL | 785 | A | LEU | 699 | C |
| ASN | 370 | A | LYS | 417 | C |
| GLN | 895 | A | SER | 711 | C |
| ASP | 228 | A | LEU | 518 | C |
| PHE | 43 | A | GLN | 563 | C |
| GLU | 1144 | A | LEU | 1141 | C |
| TYR | 200 | A | TYR | 396 | C |
| LEU | 560 | A | ASN | 282 | B |
| GLU | 918 | A | VAL | 1129 | C |

| | | | | | |
|---|---|---|---|---|---|
| LYS | 417 | A | ASN | 370 | B |
| TYR | 505 | A | ALA | 372 | B |
| GLN | 965 | A | GLY | 757 | B |
| VAL | 705 | A | ALA | 893 | B |
| THR | 167 | A | ARG | 466 | C |
| ASP | 737 | A | ARG | 319 | C |
| GLU | 224 | A | LEU | 560 | C |
| ALA | 706 | A | GLN | 895 | B |
| TYR | 707 | A | ILE | 896 | B |
| GLY | 889 | A | LYS | 1045 | C |
| LEU | 546 | A | ASP | 979 | B |
| PHE | 1089 | A | ASN | 914 | B |
| GLY | 283 | A | GLN | 563 | C |
| ASP | 568 | A | ASN | 856 | B |
| ASN | 856 | A | PHE | 592 | C |
| LYS | 786 | A | GLY | 700 | C |
| LYS | 386 | A | LEU | 984 | B |
| ILE | 1130 | A | GLN | 920 | B |
| SER | 711 | A | PRO | 897 | B |
| ARG | 355 | A | PRO | 230 | B |
| ASP | 737 | A | ASN | 317 | C |
| GLN | 787 | A | ALA | 701 | C |
| GLY | 381 | A | ILE | 973 | B |
| ALA | 899 | A | TYR | 707 | C |
| TRP | 886 | A | TYR | 1047 | C |
| GLU | 1144 | A | LEU | 1145 | C |
| PRO | 897 | A | TYR | 707 | C |
| LEU | 981 | A | LYS | 386 | C |
| GLY | 566 | A | VAL | 42 | B |
| ALA | 972 | A | GLN | 755 | B |
| ASP | 40 | A | PHE | 562 | C |
| ILE | 468 | A | GLU | 132 | B |
| ALA | 701 | A | LYS | 786 | B |
| ASP | 985 | A | SER | 383 | C |
| TYR | 873 | A | SER | 698 | C |
| TYR | 707 | A | THR | 883 | B |
| LEU | 699 | A | GLN | 872 | B |
| GLY | 199 | A | PRO | 463 | C |
| LEU | 546 | A | VAL | 976 | B |
| SER | 383 | A | ASP | 985 | B |
| ILE | 788 | A | GLU | 702 | C |

| | | | | | |
|---|---|---|---|---|---|
| THR | 768 | A | GLN | 314 | C |
| THR | 572 | A | VAL | 976 | B |
| SER | 45 | A | LYS | 557 | C |
| ALA | 668 | A | LEU | 864 | B |
| GLN | 787 | A | GLU | 702 | C |
| TYR | 917 | A | PRO | 1079 | C |
| ASP | 389 | A | SER | 982 | B |
| ASN | 960 | A | ILE | 569 | C |
| ARG | 466 | A | ILE | 231 | B |
| HIS | 519 | A | LYS | 41 | B |
| MET | 740 | A | PRO | 589 | C |
| MET | 740 | A | ARG | 319 | C |
| ASP | 1041 | A | SER | 1030 | B |
| GLY | 199 | A | PHE | 464 | C |
| MET | 900 | A | ILE | 712 | C |
| THR | 998 | A | GLN | 1002 | C |
| TRP | 353 | A | GLY | 232 | B |
| PHE | 759 | A | PHE | 970 | C |
| GLU | 1017 | A | LYS | 776 | B |
| PHE | 43 | A | LEU | 560 | C |
| ILE | 896 | A | ALA | 713 | C |
| LYS | 1045 | A | ALA | 890 | B |
| PHE | 562 | A | TYR | 38 | B |
| GLN | 1002 | A | PHE | 759 | B |
| ILE | 973 | A | GLY | 381 | C |
| TYR | 904 | A | ILE | 712 | C |
| GLN | 762 | A | GLN | 1010 | C |
| TYR | 38 | A | PHE | 562 | C |
| SER | 967 | A | ILE | 569 | C |
| TYR | 904 | A | GLY | 1093 | C |
| ILE | 233 | A | PHE | 464 | C |
| LYS | 386 | A | ASP | 985 | B |
| LEU | 864 | A | CYS | 671 | C |
| LYS | 786 | A | LEU | 699 | C |
| ARG | 44 | A | ARG | 567 | C |
| PHE | 1095 | A | MET | 900 | B |

**Table S4.** The list of the inter-protomer contacts of the protomer A in the structure of SARS-CoV-2 S-GSAS/G614 in the open state (pdb id 7KDL).

| S protein residue name | S protein residue number | S protein chain | S protein residue name | S protein residue number | S protein chain |
|---|---|---|---|---|---|
| VAL | 1040 | A | GLN | 1036 | B |
| VAL | 1068 | A | GLY | 891 | B |
| ILE | 788 | A | SER | 704 | C |
| LEU | 966 | A | ALA | 570 | C |
| VAL | 785 | A | LEU | 699 | C |
| ILE | 870 | A | LEU | 699 | C |
| SER | 758 | A | LYS | 964 | C |
| TYR | 380 | A | ILE | 973 | B |
| GLN | 762 | A | THR | 1006 | C |
| ASN | 710 | A | PRO | 897 | B |
| VAL | 42 | A | PHE | 562 | C |
| THR | 430 | A | ARG | 983 | B |
| LYS | 557 | A | SER | 45 | B |
| SER | 383 | A | LEU | 984 | B |
| GLU | 918 | A | SER | 1123 | C |
| GLN | 1036 | A | VAL | 1040 | C |
| PHE | 759 | A | GLN | 965 | C |
| THR | 696 | A | MET | 869 | B |
| PHE | 759 | A | GLN | 1002 | C |
| GLN | 314 | A | SER | 735 | B |
| GLY | 999 | A | PHE | 759 | B |
| GLN | 755 | A | SER | 968 | C |
| ASP | 1041 | A | GLN | 784 | B |
| GLN | 414 | A | VAL | 987 | C |
| SER | 968 | A | GLN | 755 | B |
| TYR | 707 | A | ILE | 896 | B |
| THR | 859 | A | GLN | 613 | C |
| THR | 883 | A | TYR | 707 | C |
| SER | 1030 | A | ASP | 1041 | C |
| ASP | 1041 | A | GLY | 889 | B |
| GLN | 314 | A | LEU | 861 | B |
| MET | 869 | A | GLY | 669 | C |
| GLN | 895 | A | ALA | 713 | C |
| ALA | 890 | A | VAL | 1068 | C |
| GLY | 232 | A | GLU | 465 | C |

| | | | | | |
|---|---|---|---|---|---|
| TYR | 38 | A | PHE | 562 | C |
| ARG | 1091 | A | ASP | 1118 | B |
| LEU | 560 | A | TYR | 38 | B |
| GLN | 784 | A | ASP | 1041 | C |
| ILE | 569 | A | VAL | 47 | B |
| GLU | 702 | A | GLN | 787 | B |
| GLN | 1002 | A | GLN | 1002 | C |
| PHE | 43 | A | LEU | 560 | C |
| GLN | 913 | A | PRO | 1090 | C |
| ALA | 892 | A | GLU | 1072 | C |
| ARG | 44 | A | ARG | 567 | C |
| GLU | 1144 | A | LEU | 1145 | C |
| GLU | 988 | A | SER | 383 | C |
| TYR | 279 | A | GLN | 563 | C |
| GLY | 669 | A | LEU | 864 | B |
| ALA | 890 | A | TYR | 1067 | C |
| LEU | 864 | A | CYS | 662 | C |
| PRO | 521 | A | LYS | 41 | B |
| GLY | 757 | A | GLN | 965 | C |
| SER | 758 | A | THR | 961 | C |
| ASP | 979 | A | ARG | 567 | C |
| TYR | 904 | A | VAL | 1094 | C |
| TYR | 707 | A | GLY | 798 | B |
| PRO | 384 | A | GLU | 988 | B |
| PRO | 897 | A | ASN | 709 | C |
| GLY | 1046 | A | GLY | 889 | B |
| TYR | 917 | A | VAL | 1129 | C |
| VAL | 705 | A | PRO | 792 | B |
| THR | 887 | A | TYR | 1047 | C |
| PHE | 970 | A | GLN | 755 | B |
| ASN | 703 | A | TYR | 789 | B |
| SER | 383 | A | GLU | 988 | B |
| ALA | 892 | A | PRO | 1069 | C |
| PHE | 559 | A | PHE | 43 | B |
| PRO | 230 | A | ARG | 355 | C |
| ALA | 570 | A | LEU | 966 | B |
| ILE | 569 | A | VAL | 963 | B |
| GLN | 564 | A | PHE | 43 | B |
| PRO | 862 | A | ALA | 668 | C |
| ALA | 899 | A | TYR | 707 | C |
| THR | 1009 | A | THR | 1009 | C |

| | | | | | |
|---|---|---|---|---|---|
| ASN | 978 | A | THR | 547 | C |
| CYS | 379 | A | GLU | 988 | B |
| GLY | 594 | A | THR | 859 | B |
| LEU | 699 | A | LYS | 786 | B |
| LEU | 517 | A | ARG | 983 | B |
| GLN | 787 | A | ALA | 701 | C |
| TYR | 707 | A | ALA | 899 | B |
| ILE | 896 | A | ALA | 713 | C |
| ILE | 1013 | A | LEU | 1012 | B |
| GLY | 199 | A | ARG | 355 | C |
| TYR | 1047 | A | TRP | 886 | B |
| ILE | 233 | A | GLU | 465 | C |
| VAL | 1094 | A | ILE | 896 | B |
| VAL | 976 | A | LEU | 546 | C |
| GLN | 965 | A | TYR | 756 | B |
| ASP | 979 | A | LEU | 546 | C |
| LEU | 1141 | A | LEU | 1141 | B |
| LYS | 386 | A | LEU | 981 | B |
| ASN | 703 | A | LYS | 790 | B |
| LEU | 1145 | A | LEU | 1145 | B |
| ASP | 1139 | A | GLU | 1144 | B |
| GLY | 413 | A | LYS | 986 | C |
| ARG | 983 | A | LYS | 386 | C |
| LEU | 865 | A | MET | 697 | C |
| ALA | 570 | A | ASN | 856 | B |
| THR | 547 | A | SER | 982 | B |
| GLU | 1072 | A | LEU | 894 | B |
| TRP | 886 | A | TYR | 1047 | C |
| GLN | 913 | A | ARG | 1107 | C |
| GLN | 1142 | A | GLU | 1144 | B |
| GLN | 895 | A | SER | 708 | C |
| PHE | 855 | A | PRO | 589 | C |
| ASP | 796 | A | TYR | 707 | C |
| THR | 385 | A | THR | 415 | C |
| GLY | 1093 | A | GLN | 913 | B |
| LEU | 894 | A | GLU | 1072 | C |
| ASN | 856 | A | ASP | 568 | C |
| MET | 900 | A | PRO | 1079 | C |
| PRO | 863 | A | ALA | 668 | C |
| GLY | 857 | A | PHE | 592 | C |
| ASP | 427 | A | GLU | 990 | C |

| | | | | | |
|---|---|---|---|---|---|
| GLU | 773 | A | GLU | 1017 | C |
| ASP | 985 | A | THR | 385 | C |
| ALA | 570 | A | LYS | 964 | B |
| GLN | 1002 | A | TYR | 756 | B |
| LEU | 894 | A | ALA | 713 | C |
| LEU | 981 | A | LYS | 386 | C |
| GLY | 669 | A | MET | 869 | B |
| TYR | 904 | A | ARG | 1107 | C |
| GLY | 798 | A | TYR | 707 | C |
| VAL | 705 | A | THR | 883 | B |
| GLY | 798 | A | ILE | 1130 | C |
| LEU | 894 | A | ILE | 714 | C |
| VAL | 1128 | A | ASN | 919 | B |
| VAL | 963 | A | ASP | 568 | C |
| LYS | 790 | A | ASN | 703 | C |
| GLN | 314 | A | THR | 768 | B |
| VAL | 705 | A | ALA | 879 | B |
| ASP | 198 | A | ASP | 428 | C |
| PHE | 1089 | A | ASN | 914 | B |
| HIS | 519 | A | VAL | 42 | B |
| THR | 588 | A | PHE | 855 | B |
| VAL | 705 | A | GLN | 895 | B |
| MET | 740 | A | CYS | 590 | C |
| PRO | 412 | A | VAL | 987 | C |
| ARG | 1039 | A | ARG | 1039 | C |
| ASP | 228 | A | TYR | 396 | C |
| ARG | 1107 | A | TRP | 886 | B |
| SER | 704 | A | ILE | 788 | B |
| TYR | 1047 | A | ALA | 890 | B |
| VAL | 1094 | A | TYR | 904 | B |
| ARG | 765 | A | THR | 961 | C |
| ARG | 765 | A | GLN | 957 | C |
| ILE | 231 | A | ARG | 355 | C |
| SER | 383 | A | ASP | 985 | B |
| THR | 961 | A | GLN | 762 | B |
| ILE | 1013 | A | ALA | 1016 | B |
| PRO | 665 | A | LEU | 864 | B |
| GLN | 613 | A | THR | 859 | B |
| LEU | 1141 | A | LEU | 1141 | C |
| ASP | 994 | A | PHE | 970 | C |
| LEU | 560 | A | ASN | 282 | B |

| | | | | | |
|---|---|---|---|---|---|
| TYR | 200 | A | SER | 514 | C |
| ASN | 856 | A | PHE | 592 | C |
| SER | 1003 | A | PHE | 759 | B |
| ASN | 282 | A | LYS | 558 | C |
| GLU | 1072 | A | ALA | 893 | B |
| GLY | 889 | A | LYS | 1045 | C |
| GLN | 787 | A | ASN | 703 | C |
| GLN | 787 | A | GLY | 700 | C |
| ILE | 712 | A | MET | 900 | B |
| TYR | 873 | A | LEU | 699 | C |
| GLN | 762 | A | THR | 961 | C |
| LEU | 966 | A | ASP | 571 | C |
| GLU | 918 | A | VAL | 1128 | C |
| TYR | 200 | A | ARG | 355 | C |
| PHE | 592 | A | THR | 859 | B |
| ARG | 1019 | A | GLU | 1017 | C |
| LYS | 1045 | A | GLY | 889 | B |
| GLN | 762 | A | TYR | 1007 | C |
| PRO | 1069 | A | ALA | 890 | B |
| VAL | 963 | A | ILE | 569 | C |
| LYS | 558 | A | PHE | 43 | B |
| TYR | 707 | A | ILE | 882 | B |
| GLY | 889 | A | ASP | 1041 | C |
| ASN | 317 | A | GLY | 857 | B |
| PHE | 43 | A | LYS | 557 | C |
| PHE | 562 | A | LYS | 41 | B |
| ILE | 1013 | A | ILE | 1013 | C |
| VAL | 382 | A | GLU | 988 | B |
| GLU | 1111 | A | VAL | 1122 | C |
| PHE | 562 | A | PRO | 225 | B |
| VAL | 1094 | A | MET | 900 | B |
| PRO | 862 | A | GLY | 667 | C |
| GLY | 566 | A | VAL | 42 | B |
| LEU | 560 | A | PHE | 43 | B |
| ALA | 701 | A | ILE | 788 | B |
| THR | 1027 | A | ARG | 1039 | C |
| THR | 961 | A | SER | 758 | B |
| ARG | 983 | A | GLY | 381 | C |
| THR | 167 | A | ARG | 466 | C |
| SER | 982 | A | ASP | 389 | C |
| THR | 547 | A | ASP | 979 | B |

| | | | | | |
|---|---|---|---|---|---|
| MET | 900 | A | THR | 1077 | C |
| PRO | 1079 | A | GLN | 913 | B |
| LYS | 1045 | A | ALA | 890 | B |
| GLN | 1010 | A | LEU | 1012 | B |
| GLY | 667 | A | PRO | 863 | B |
| PHE | 898 | A | TYR | 707 | C |
| SER | 1123 | A | ASN | 914 | B |
| ILE | 712 | A | TYR | 904 | B |
| ILE | 973 | A | GLY | 381 | C |
| PHE | 759 | A | GLY | 999 | C |
| LEU | 1145 | A | LEU | 1141 | B |
| ASP | 1041 | A | SER | 1030 | B |
| GLY | 880 | A | VAL | 705 | C |
| PHE | 43 | A | LYS | 558 | C |
| THR | 1006 | A | GLN | 1005 | B |
| ASN | 914 | A | PHE | 1121 | C |
| ARG | 983 | A | VAL | 382 | C |
| THR | 883 | A | VAL | 705 | C |
| LEU | 1034 | A | ASP | 1041 | C |
| GLY | 1046 | A | ALA | 890 | B |
| LYS | 790 | A | VAL | 705 | C |
| GLU | 918 | A | VAL | 1129 | C |
| GLY | 566 | A | PHE | 43 | B |
| LYS | 786 | A | GLY | 700 | C |
| LEU | 560 | A | GLY | 283 | B |
| ASP | 40 | A | PHE | 562 | C |
| LYS | 41 | A | ALA | 520 | C |
| PHE | 970 | A | PHE | 759 | B |
| LEU | 894 | A | ARG | 1107 | C |
| PRO | 1069 | A | ALA | 892 | B |
| MET | 869 | A | THR | 696 | C |
| TYR | 200 | A | TYR | 396 | C |
| LEU | 1012 | A | GLN | 1010 | C |
| PHE | 1121 | A | GLN | 913 | B |
| GLN | 564 | A | VAL | 42 | B |
| ASP | 40 | A | GLN | 563 | C |
| ARG | 1039 | A | GLU | 1031 | B |
| SER | 698 | A | TYR | 873 | B |
| GLN | 563 | A | LYS | 41 | B |
| ALA | 1078 | A | MET | 900 | B |
| GLU | 1144 | A | LEU | 1141 | C |

| | | | | | |
|---|---|---|---|---|---|
| SER | 967 | A | ILE | 569 | C |
| SER | 975 | A | ALA | 570 | C |
| ALA | 701 | A | GLN | 787 | B |
| GLN | 957 | A | ARG | 765 | B |
| TYR | 707 | A | LYS | 795 | B |
| ALA | 570 | A | VAL | 963 | B |
| GLU | 1031 | A | VAL | 1040 | C |
| LYS | 557 | A | PHE | 43 | B |
| GLU | 1072 | A | ALA | 892 | B |
| PHE | 43 | A | GLN | 563 | C |
| THR | 859 | A | GLY | 594 | C |
| ALA | 713 | A | LEU | 894 | B |
| GLY | 413 | A | VAL | 987 | C |
| VAL | 382 | A | LEU | 984 | B |
| VAL | 1040 | A | SER | 1030 | B |
| ALA | 1016 | A | ILE | 1013 | C |
| VAL | 47 | A | ILE | 569 | C |
| VAL | 382 | A | ARG | 983 | B |
| ILE | 882 | A | TYR | 707 | C |
| PRO | 863 | A | GLY | 669 | C |
| GLY | 891 | A | GLY | 1046 | C |
| SER | 967 | A | ALA | 570 | C |
| THR | 302 | A | THR | 761 | B |
| GLY | 891 | A | PRO | 1069 | C |
| PHE | 43 | A | PHE | 565 | C |
| ASN | 969 | A | GLN | 755 | B |
| ALA | 668 | A | LEU | 864 | B |
| PHE | 562 | A | TYR | 38 | B |
| ILE | 788 | A | LEU | 699 | C |
| GLU | 1092 | A | GLN | 913 | B |
| GLY | 283 | A | GLN | 563 | C |
| ARG | 1107 | A | LEU | 894 | B |
| GLN | 965 | A | GLN | 762 | B |
| TYR | 396 | A | PRO | 230 | B |
| TYR | 707 | A | GLN | 895 | B |
| SER | 1030 | A | VAL | 1040 | C |
| GLN | 895 | A | SER | 711 | C |
| MET | 697 | A | LEU | 865 | B |
| PHE | 43 | A | PHE | 559 | C |
| ALA | 668 | A | LEU | 865 | B |
| ILE | 670 | A | LEU | 864 | B |

| | | | | | |
|---|---|---|---|---|---|
| LEU | 699 | A | ILE | 788 | B |
| LEU | 699 | A | VAL | 785 | B |
| THR | 1009 | A | ILE | 1013 | C |
| LEU | 984 | A | SER | 383 | C |
| LEU | 861 | A | GLN | 613 | C |
| ASP | 994 | A | ARG | 995 | C |
| LEU | 894 | A | VAL | 705 | C |
| LYS | 921 | A | ILE | 1130 | C |
| MET | 900 | A | PHE | 1095 | C |
| TYR | 707 | A | THR | 883 | B |
| ALA | 899 | A | PRO | 1079 | C |
| SER | 758 | A | GLN | 965 | C |
| LEU | 699 | A | GLN | 872 | B |
| LYS | 41 | A | GLN | 564 | C |
| LEU | 1034 | A | VAL | 1040 | C |
| ILE | 794 | A | TYR | 707 | C |
| MET | 697 | A | TYR | 873 | B |
| SER | 982 | A | THR | 547 | C |
| TYR | 917 | A | ILE | 1130 | C |
| LEU | 858 | A | PHE | 592 | C |
| LEU | 984 | A | GLY | 381 | C |
| ALA | 701 | A | TYR | 789 | B |
| GLU | 1031 | A | ARG | 1039 | C |
| TYR | 756 | A | GLN | 1002 | C |
| GLY | 669 | A | PRO | 863 | B |
| TYR | 707 | A | ILE | 794 | B |
| GLN | 1005 | A | THR | 1006 | C |
| GLY | 971 | A | TYR | 756 | B |
| GLU | 1111 | A | SER | 1123 | C |
| PRO | 792 | A | TYR | 707 | C |
| ASP | 1041 | A | ALA | 890 | B |
| GLU | 1017 | A | GLU | 773 | B |
| PHE | 565 | A | PHE | 43 | B |
| GLN | 1002 | A | GLN | 1005 | B |
| ALA | 668 | A | PRO | 863 | B |
| ASP | 796 | A | SER | 708 | C |
| ARG | 357 | A | PRO | 230 | B |
| SER | 735 | A | GLN | 314 | C |
| VAL | 963 | A | ALA | 570 | C |
| ALA | 890 | A | GLY | 1046 | C |
| LEU | 894 | A | PRO | 1069 | C |

| | | | | | |
|---|---|---|---|---|---|
| ARG | 983 | A | LEU | 390 | C |
| GLY | 669 | A | THR | 866 | B |
| ASN | 914 | A | SER | 1123 | C |
| GLN | 564 | A | LYS | 41 | B |
| PHE | 1089 | A | TYR | 917 | B |
| PRO | 897 | A | SER | 711 | C |
| TYR | 707 | A | PHE | 898 | B |
| GLY | 381 | A | ILE | 973 | B |
| ARG | 567 | A | ARG | 44 | B |
| GLY | 1046 | A | GLY | 891 | B |
| ILE | 231 | A | PHE | 464 | C |
| ASN | 919 | A | VAL | 1128 | C |
| ASP | 1041 | A | LEU | 1034 | B |
| LEU | 1145 | A | GLU | 1144 | B |
| ARG | 1107 | A | GLN | 913 | B |
| SER | 982 | A | LEU | 390 | C |
| GLY | 1124 | A | GLU | 918 | B |
| ALA | 893 | A | VAL | 705 | C |
| THR | 1006 | A | GLN | 762 | B |
| GLN | 755 | A | ASN | 969 | C |
| GLY | 566 | A | ARG | 44 | B |
| THR | 547 | A | LEU | 981 | B |
| VAL | 42 | A | GLN | 563 | C |
| THR | 859 | A | PHE | 592 | C |
| ALA | 570 | A | SER | 967 | B |
| ASP | 198 | A | PHE | 464 | C |
| PRO | 897 | A | TYR | 707 | C |
| ARG | 983 | A | LEU | 517 | C |
| MET | 900 | A | VAL | 1094 | C |
| VAL | 1040 | A | GLY | 889 | B |
| VAL | 42 | A | GLY | 566 | C |
| ASP | 571 | A | SER | 967 | B |
| ARG | 567 | A | PHE | 43 | B |
| THR | 549 | A | ASP | 745 | B |
| LYS | 786 | A | LEU | 699 | C |
| GLY | 381 | A | LEU | 984 | B |
| TYR | 369 | A | THR | 415 | C |
| THR | 1009 | A | THR | 1009 | B |
| ILE | 712 | A | PRO | 897 | B |
| GLN | 779 | A | MET | 697 | C |
| LYS | 790 | A | SER | 704 | C |

| | | | | | |
|---|---|---|---|---|---|
| SER | 1123 | A | GLU | 1111 | B |
| SER | 982 | A | LYS | 386 | C |
| THR | 547 | A | ASN | 978 | B |
| ILE | 1130 | A | LYS | 921 | B |
| LEU | 984 | A | LYS | 386 | C |
| VAL | 860 | A | ARG | 646 | C |
| PRO | 1079 | A | TYR | 917 | B |
| ASN | 709 | A | PRO | 897 | B |
| TYR | 756 | A | SER | 968 | C |
| LEU | 699 | A | LEU | 865 | B |
| TYR | 904 | A | ILE | 712 | C |
| SER | 708 | A | GLN | 895 | B |
| GLN | 913 | A | PHE | 1089 | C |
| ILE | 896 | A | ILE | 712 | C |
| SER | 704 | A | TYR | 789 | B |
| ALA | 706 | A | GLN | 895 | B |
| GLN | 1002 | A | THR | 998 | B |
| THR | 572 | A | ASN | 856 | B |
| LEU | 546 | A | ASP | 979 | B |
| VAL | 1040 | A | GLY | 1035 | B |
| ILE | 712 | A | ILE | 896 | B |
| ASN | 856 | A | ALA | 570 | C |
| LEU | 699 | A | ILE | 870 | B |
| GLN | 762 | A | GLN | 1010 | C |
| SER | 708 | A | PRO | 897 | B |
| GLY | 757 | A | PHE | 970 | C |
| GLY | 889 | A | GLY | 1046 | C |
| PHE | 592 | A | MET | 740 | B |
| VAL | 1128 | A | TYR | 917 | B |
| PHE | 970 | A | TYR | 756 | B |
| PRO | 897 | A | SER | 708 | C |
| ARG | 567 | A | VAL | 976 | B |
| LEU | 984 | A | VAL | 382 | C |
| ASN | 978 | A | GLY | 545 | C |
| ARG | 983 | A | SER | 383 | C |
| LYS | 41 | A | GLN | 563 | C |
| GLN | 895 | A | VAL | 705 | C |
| VAL | 42 | A | ARG | 567 | C |
| THR | 284 | A | LEU | 560 | C |
| ASP | 571 | A | VAL | 976 | B |
| MET | 869 | A | MET | 697 | C |

| | | | | | |
|---|---|---|---|---|---|
| GLN | 563 | A | TYR | 279 | B |
| GLY | 199 | A | PHE | 464 | C |
| TYR | 904 | A | GLY | 1093 | C |
| LYS | 964 | A | ILE | 569 | C |
| PRO | 1069 | A | LEU | 894 | B |
| VAL | 976 | A | THR | 572 | C |
| ALA | 890 | A | ASP | 1041 | C |
| GLN | 895 | A | ASN | 1074 | C |
| ARG | 1039 | A | ARG | 1039 | B |
| TYR | 756 | A | GLY | 971 | C |
| SER | 968 | A | GLY | 757 | B |
| PHE | 1089 | A | GLN | 913 | B |
| VAL | 1040 | A | GLU | 1031 | B |
| ILE | 1130 | A | GLY | 798 | B |
| TYR | 756 | A | ARG | 995 | C |
| SER | 375 | A | ARG | 408 | C |
| LEU | 864 | A | ILE | 666 | C |
| ASN | 317 | A | MET | 740 | B |
| ARG | 1019 | A | ALA | 1016 | C |
| SER | 758 | A | SER | 968 | C |
| VAL | 705 | A | LYS | 790 | B |
| ILE | 973 | A | TYR | 380 | C |
| LEU | 864 | A | CYS | 671 | C |
| ASN | 317 | A | ASP | 737 | B |
| PHE | 43 | A | GLY | 566 | C |
| ILE | 666 | A | LEU | 864 | B |
| ALA | 713 | A | GLN | 895 | B |
| ALA | 713 | A | ILE | 896 | B |
| LEU | 560 | A | THR | 284 | B |
| GLN | 755 | A | GLY | 971 | C |
| GLY | 700 | A | ILE | 788 | B |
| ARG | 1107 | A | TYR | 904 | B |
| THR | 859 | A | GLY | 593 | C |
| VAL | 705 | A | GLY | 880 | B |
| GLY | 757 | A | SER | 968 | C |
| LEU | 48 | A | ILE | 569 | C |
| VAL | 1128 | A | GLU | 918 | B |
| ALA | 890 | A | LYS | 1045 | C |
| ARG | 1091 | A | ASN | 907 | B |
| GLY | 413 | A | ASP | 985 | C |
| GLN | 895 | A | TYR | 707 | C |

| | | | | | |
|---|---|---|---|---|---|
| CYS | 166 | A | ILE | 468 | C |
| GLN | 563 | A | PHE | 43 | B |
| PRO | 792 | A | VAL | 705 | C |
| LEU | 864 | A | PRO | 665 | C |
| GLY | 700 | A | LYS | 786 | B |
| SER | 982 | A | LEU | 546 | C |
| LEU | 1145 | A | LEU | 1141 | C |
| GLN | 563 | A | ARG | 44 | B |
| THR | 1077 | A | MET | 900 | B |
| SER | 704 | A | LYS | 790 | B |
| GLU | 224 | A | PHE | 562 | C |
| PRO | 1090 | A | ASN | 907 | B |
| ILE | 233 | A | ARG | 466 | C |
| SER | 711 | A | PRO | 897 | B |
| ALA | 668 | A | MET | 869 | B |
| PHE | 797 | A | TYR | 707 | C |
| GLN | 321 | A | ASP | 745 | B |
| PRO | 1090 | A | GLN | 913 | B |
| GLN | 755 | A | ALA | 972 | C |
| ALA | 893 | A | GLU | 1072 | C |
| ILE | 1130 | A | TYR | 917 | B |
| LYS | 41 | A | HIS | 519 | C |
| ASN | 394 | A | PRO | 230 | B |
| PHE | 562 | A | GLU | 224 | B |
| GLN | 1010 | A | ALA | 766 | B |
| LYS | 41 | A | PHE | 565 | C |
| PHE | 1042 | A | GLU | 1031 | B |
| GLY | 971 | A | GLN | 755 | B |
| TYR | 707 | A | ASP | 796 | B |
| GLN | 965 | A | SER | 758 | B |
| ASP | 571 | A | SER | 975 | B |
| VAL | 42 | A | HIS | 519 | C |
| ALA | 893 | A | ASN | 703 | C |
| SER | 982 | A | GLY | 545 | C |
| VAL | 42 | A | PHE | 565 | C |
| VAL | 1129 | A | TYR | 917 | B |
| SER | 968 | A | SER | 758 | B |
| MET | 697 | A | MET | 869 | B |
| ARG | 319 | A | MET | 740 | B |
| PHE | 759 | A | THR | 1006 | C |
| TYR | 1007 | A | GLN | 762 | B |

| | | | | | |
|---|---|---|---|---|---|
| GLU | 224 | A | LEU | 560 | C |
| GLN | 563 | A | GLY | 283 | B |
| ILE | 788 | A | ALA | 701 | C |
| SER | 884 | A | VAL | 705 | C |
| THR | 866 | A | ALA | 668 | C |
| CYS | 662 | A | LEU | 864 | B |
| GLY | 667 | A | PRO | 862 | B |
| PHE | 592 | A | GLY | 857 | B |
| ALA | 1020 | A | ARG | 1019 | B |
| PRO | 1079 | A | ALA | 899 | B |
| GLY | 700 | A | TYR | 873 | B |
| ASP | 985 | A | LYS | 386 | C |
| PRO | 715 | A | LEU | 894 | B |
| GLN | 913 | A | GLY | 1093 | C |
| ASP | 389 | A | SER | 982 | B |
| ASP | 198 | A | PRO | 426 | C |
| ALA | 668 | A | PRO | 862 | B |
| VAL | 705 | A | SER | 884 | B |
| ASP | 985 | A | SER | 383 | C |
| LYS | 386 | A | LEU | 984 | B |
| PRO | 897 | A | ASN | 710 | C |
| ALA | 668 | A | THR | 866 | B |
| ARG | 44 | A | GLN | 563 | C |
| GLY | 889 | A | VAL | 1040 | C |
| SER | 975 | A | ASP | 571 | C |
| PRO | 862 | A | ALA | 647 | C |
| LEU | 864 | A | GLY | 667 | C |
| PHE | 1121 | A | ASN | 914 | B |
| PRO | 589 | A | PHE | 855 | B |
| ALA | 701 | A | LYS | 786 | B |
| SER | 967 | A | ASP | 571 | C |
| TRP | 886 | A | ARG | 1107 | C |
| ASP | 796 | A | ASN | 709 | C |
| LEU | 1145 | A | LEU | 1145 | C |
| ILE | 569 | A | SER | 967 | B |
| ASN | 1108 | A | TRP | 886 | B |
| GLN | 314 | A | ASP | 737 | B |
| ILE | 569 | A | LYS | 964 | B |
| ASN | 709 | A | ASP | 796 | B |
| ILE | 712 | A | GLN | 895 | B |
| PHE | 970 | A | GLY | 757 | B |

| | | | | | |
|---|---|---|---|---|---|
| LYS | 558 | A | ASN | 282 | B |
| TYR | 789 | A | SER | 704 | C |
| PRO | 589 | A | MET | 740 | B |
| GLN | 920 | A | ILE | 1130 | C |
| ILE | 788 | A | GLY | 700 | C |
| TYR | 789 | A | ALA | 701 | C |
| LEU | 1001 | A | GLN | 1002 | C |
| CYS | 671 | A | LEU | 864 | B |
| SER | 708 | A | ILE | 896 | B |
| HIS | 519 | A | ASP | 40 | B |
| MET | 740 | A | PHE | 592 | C |
| ASP | 40 | A | HIS | 519 | C |
| ASN | 1074 | A | GLN | 895 | B |
| LYS | 790 | A | GLU | 702 | C |
| PHE | 759 | A | SER | 1003 | C |
| TRP | 886 | A | ASN | 1108 | C |
| ASP | 737 | A | ASN | 317 | C |
| ASP | 745 | A | ARG | 319 | C |
| GLY | 283 | A | LEU | 560 | C |
| GLY | 232 | A | ARG | 466 | C |
| SER | 1123 | A | GLU | 918 | B |
| GLU | 918 | A | GLY | 1124 | C |
| GLN | 563 | A | VAL | 42 | B |
| GLY | 199 | A | PRO | 463 | C |
| VAL | 705 | A | TYR | 789 | B |
| GLN | 1010 | A | VAL | 1008 | B |
| TYR | 917 | A | PHE | 1089 | C |
| TYR | 873 | A | SER | 698 | C |
| TYR | 1067 | A | ALA | 890 | B |
| SER | 711 | A | GLN | 895 | B |
| ALA | 890 | A | PRO | 1069 | C |
| TYR | 917 | A | PRO | 1079 | C |
| PRO | 1079 | A | MET | 900 | B |
| TYR | 789 | A | ASN | 703 | C |
| LEU | 390 | A | SER | 982 | B |
| GLY | 199 | A | GLU | 465 | C |
| LYS | 964 | A | SER | 758 | B |
| TYR | 707 | A | PRO | 897 | B |
| GLN | 965 | A | GLY | 757 | B |
| SER | 1147 | A | LEU | 1145 | C |
| GLY | 232 | A | PHE | 464 | C |

| | | | | | |
|---|---|---|---|---|---|
| ARG | 44 | A | GLY | 566 | C |
| GLN | 1002 | A | LEU | 1001 | B |
| ILE | 788 | A | GLU | 702 | C |
| VAL | 1040 | A | LEU | 1034 | B |
| THR | 768 | A | GLN | 314 | C |
| PHE | 1095 | A | MET | 900 | B |
| GLN | 787 | A | GLU | 702 | C |
| LEU | 981 | A | THR | 547 | C |
| GLU | 918 | A | PHE | 1089 | C |
| GLY | 548 | A | ASN | 978 | B |
| PHE | 562 | A | ASP | 40 | B |
| TYR | 756 | A | GLN | 965 | C |
| GLU | 702 | A | ILE | 788 | B |
| MET | 740 | A | ARG | 319 | C |
| GLU | 702 | A | TYR | 789 | B |
| GLY | 381 | A | ARG | 983 | B |
| ILE | 666 | A | PRO | 862 | B |
| LYS | 386 | A | ASP | 985 | B |
| GLN | 563 | A | ASP | 40 | B |
| PHE | 1121 | A | THR | 912 | B |
| GLY | 669 | A | LEU | 865 | B |
| VAL | 1129 | A | GLU | 918 | B |
| ARG | 1107 | A | ASN | 907 | B |
| LEU | 894 | A | PRO | 715 | C |
| TYR | 707 | A | PHE | 797 | B |
| LEU | 699 | A | MET | 869 | B |
| LYS | 386 | A | ARG | 983 | B |
| LEU | 865 | A | ALA | 668 | C |
| HIS | 519 | A | LYS | 41 | B |
| LEU | 1141 | A | GLU | 1144 | B |
| ALA | 890 | A | TYR | 1047 | C |
| GLN | 965 | A | PHE | 759 | B |
| GLN | 613 | A | LEU | 861 | B |
| ASN | 282 | A | LEU | 560 | C |
| GLN | 895 | A | ILE | 712 | C |
| ILE | 231 | A | ARG | 466 | C |
| LYS | 786 | A | ALA | 701 | C |
| GLN | 1002 | A | GLN | 1002 | B |
| TYR | 789 | A | VAL | 705 | C |
| GLY | 700 | A | GLN | 787 | B |
| MET | 869 | A | ALA | 668 | C |

| | | | | | |
|---|---|---|---|---|---|
| PRO | 863 | A | GLY | 667 | C |
| TYR | 200 | A | PHE | 464 | C |
| ASN | 703 | A | GLN | 787 | B |
| VAL | 705 | A | LEU | 894 | B |
| ASP | 198 | A | PRO | 463 | C |
| LYS | 41 | A | PHE | 562 | C |
| GLY | 891 | A | VAL | 1068 | C |
| ILE | 896 | A | SER | 708 | C |
| GLN | 1005 | A | GLN | 1002 | C |
| MET | 697 | A | LEU | 864 | B |
| VAL | 976 | A | ARG | 567 | C |
| TYR | 38 | A | GLN | 563 | C |
| PHE | 759 | A | PHE | 970 | C |
| VAL | 976 | A | ASP | 571 | C |
| ILE | 714 | A | LEU | 894 | B |
| ASN | 703 | A | ILE | 788 | B |
| TYR | 873 | A | GLY | 700 | C |
| ILE | 896 | A | SER | 711 | C |
| ARG | 567 | A | VAL | 42 | B |
| ILE | 1013 | A | THR | 1009 | B |
| LEU | 865 | A | GLY | 669 | C |
| ASN | 978 | A | GLY | 548 | C |
| LYS | 1038 | A | LYS | 1038 | C |
| PRO | 230 | A | TYR | 396 | C |
| ALA | 647 | A | PRO | 862 | B |
| LEU | 865 | A | LEU | 699 | C |
| LYS | 558 | A | GLY | 283 | B |
| LEU | 864 | A | ILE | 670 | C |
| LEU | 390 | A | ARG | 983 | B |
| ALA | 879 | A | VAL | 705 | C |
| ARG | 1091 | A | GLN | 913 | B |
| TYR | 756 | A | PHE | 970 | C |
| ILE | 1013 | A | ILE | 1013 | B |
| THR | 961 | A | ARG | 765 | B |
| TYR | 917 | A | ALA | 1080 | C |
| THR | 572 | A | VAL | 976 | B |
| ILE | 788 | A | ASN | 703 | C |
| THR | 866 | A | GLY | 669 | C |
| GLY | 1093 | A | TYR | 904 | B |
| GLN | 872 | A | LEU | 699 | C |
| GLN | 913 | A | PHE | 1121 | C |

| | | | | | |
|---|---|---|---|---|---|
| MET | 900 | A | ALA | 1078 | C |
| THR | 385 | A | ASP | 985 | B |
| PHE | 592 | A | ASN | 856 | B |
| SER | 383 | A | ARG | 983 | B |
| PRO | 897 | A | THR | 1077 | C |
| VAL | 976 | A | ALA | 570 | C |
| MET | 869 | A | LEU | 699 | C |
| ILE | 569 | A | LEU | 48 | B |
| TYR | 38 | A | LEU | 560 | C |
| GLN | 762 | A | GLN | 965 | C |
| GLN | 1002 | A | PHE | 759 | B |
| PRO | 1069 | A | GLY | 891 | B |
| MET | 697 | A | GLN | 779 | B |
| PRO | 862 | A | ARG | 646 | C |
| ARG | 646 | A | PRO | 862 | B |
| VAL | 1068 | A | ALA | 890 | B |
| PRO | 897 | A | ILE | 712 | C |
| GLN | 913 | A | PRO | 1079 | C |
| TYR | 917 | A | VAL | 1128 | C |
| TYR | 789 | A | GLU | 702 | C |
| PHE | 1089 | A | GLU | 918 | B |
| MET | 900 | A | ASN | 709 | C |
| ILE | 896 | A | TYR | 707 | C |
| GLU | 868 | A | LEU | 699 | C |
| THR | 912 | A | PHE | 1121 | C |
| PHE | 592 | A | PHE | 855 | B |
| ASN | 907 | A | ARG | 1107 | C |
| SER | 968 | A | TYR | 756 | B |
| TYR | 873 | A | MET | 697 | C |
| SER | 708 | A | ASP | 796 | B |
| PHE | 43 | A | ARG | 567 | C |
| GLY | 545 | A | SER | 982 | B |
| LEU | 699 | A | GLN | 787 | B |
| LYS | 964 | A | ALA | 570 | C |
| ILE | 1130 | A | GLN | 920 | B |
| GLU | 1017 | A | ARG | 1019 | B |
| ASN | 856 | A | THR | 572 | C |
| ASN | 234 | A | ARG | 466 | C |
| GLN | 755 | A | PHE | 970 | C |
| LEU | 864 | A | GLY | 669 | C |
| THR | 998 | A | GLN | 1002 | C |

| | | | | | |
|---|---|---|---|---|---|
| MET | 900 | A | ILE | 712 | C |
| SER | 45 | A | LYS | 557 | C |
| VAL | 705 | A | ALA | 893 | B |
| GLN | 895 | A | ALA | 706 | C |
| LEU | 699 | A | TYR | 873 | B |
| ASN | 914 | A | PHE | 1089 | C |
| THR | 1077 | A | PRO | 897 | B |
| TYR | 707 | A | PRO | 792 | B |
| ARG | 319 | A | ASP | 745 | B |
| LEU | 1012 | A | ILE | 1013 | C |
| LEU | 864 | A | ALA | 668 | C |
| GLN | 563 | A | TYR | 38 | B |
| ARG | 1019 | A | ILE | 1013 | C |
| SER | 711 | A | ILE | 896 | B |
| PHE | 565 | A | LYS | 41 | B |
| LYS | 1038 | A | LYS | 1038 | B |
| GLY | 667 | A | LEU | 864 | B |
| GLU | 702 | A | LYS | 790 | B |
| TYR | 1047 | A | THR | 887 | B |
| THR | 1006 | A | PHE | 759 | B |
| PHE | 565 | A | VAL | 42 | B |
| GLN | 1010 | A | GLN | 762 | B |
| LEU | 864 | A | MET | 697 | C |
| ARG | 1039 | A | THR | 1027 | B |
| PHE | 855 | A | PHE | 592 | C |
| GLY | 1035 | A | VAL | 1040 | C |
| ALA | 520 | A | LYS | 41 | B |
| ARG | 983 | A | THR | 430 | C |
| PRO | 225 | A | PHE | 562 | C |
| LYS | 386 | A | SER | 982 | B |
